# Uncovering new drug properties in target-based drug-drug similarity networks

**DOI:** 10.1101/2020.03.12.988600

**Authors:** Lucreţia Udrescu, Paul Bogdan, Aimée Chiş, Ioan Ovidiu Sîrbu, Alexandru Topîrceanu, Renata-Maria Văruţ, Mihai Udrescu

## Abstract

Despite recent advances in bioinformatics, systems biology, and machine learning, the accurate prediction of drug properties remains an open problem. Indeed, because the biological environment is a complex system, the traditional approach – based on knowledge about the chemical structures – cannot fully explain the nature of interactions between drugs and biological targets. Consequently, in this paper, we propose an unsupervised machine learning approach that uses the information we know about drug-target interactions to infer drug properties. To this end, we define drug similarity based on drug-target interactions and build a weighted Drug-Drug Similarity Network according to the drug-drug similarity relationships. Using an energy-model network layout, we generate drug communities that are associated with specific, dominant drug properties. DrugBank confirms the properties of 59.52% of the drugs in these communities, and 26.98% are existing drug repositioning hints we reconstruct with our DDSN approach. The remaining 13.49% of the drugs seem not to match the dominant pharmacologic property; thus, we consider them as drug repurposing hints. The resources required to test all these repurposing hints are considerable. Therefore we introduce a mechanism of prioritization based on the betweenness/degree node centrality. By using betweenness/degree as an indicator of drug repurposing potential, we select Azelaic acid and Meprobamate as a possible antineoplastic and antifungal, respectively. Finally, we use a test procedure, based on molecular docking, to further analyze the repurposing of Azelaic acid and Meprobamate.

## Introduction

Conventional drug design has become expensive and cumbersome, as it requires large amounts of resources and faces serious challenges^1, 2^. Consequently, although the number of new FDA drug applications (NDAs) has significantly increased during the last decade – due to a spectacular accumulation of multi-omics data and appearance of increasingly complex bioinformatics tools – the number of approved drugs has only marginally grown (see Figure 1)^3, 4^, calling for more robust alternative strategies^5^.

**Figure 1.**
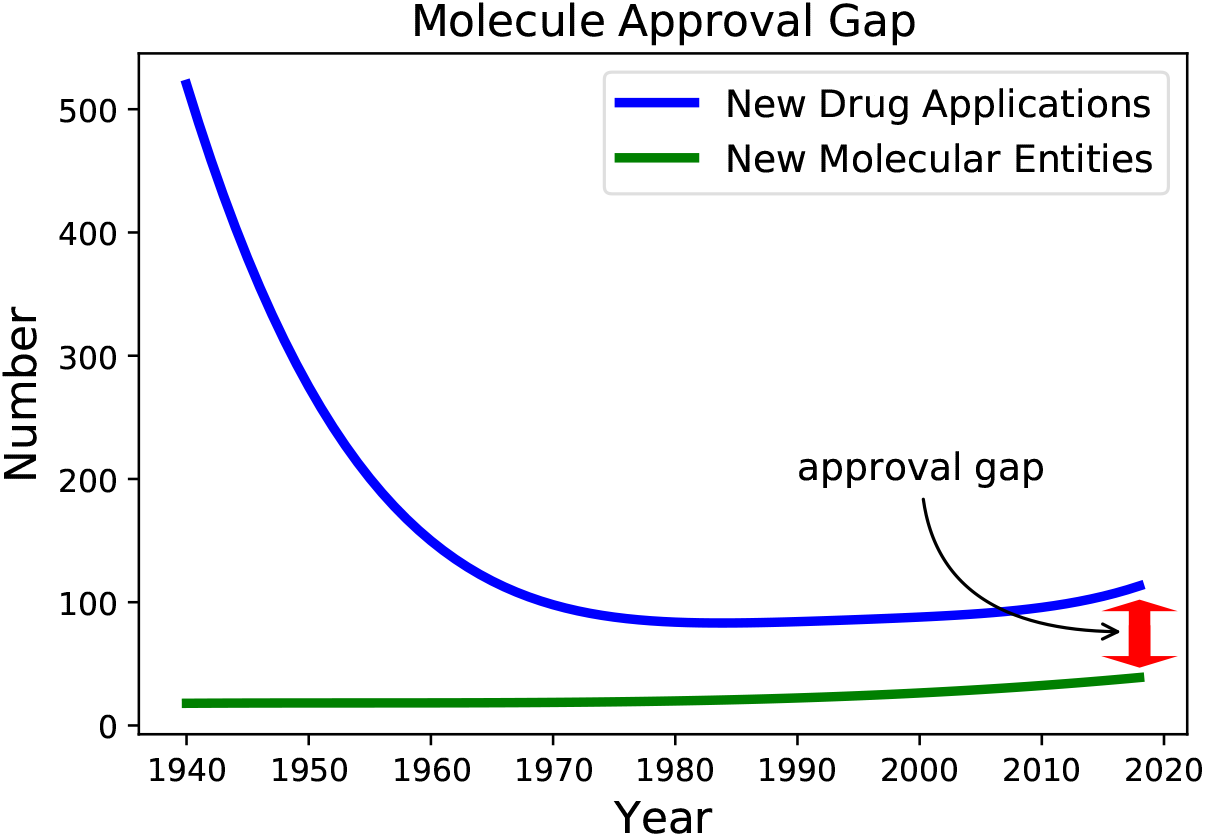
The evolution of New Drug Applications (NDAs) and New Molecular Entities (NME) during the period 1940-2017, according to FDA reports. We used the FDA’s annual reports data (https://www.fda.gov) and removed local oscillations by plotting a polynomial data fitting.

One of the most effective alternative strategies is *drug repositioning* (or *drug repurposing*)^6, 7^, namely finding new pharmaceutical functions for already used drugs. The extensive medical and pharmaceutical experience reveals a surprising propensity towards multiple indications for many drugs^8^, and the examples of successful drug repositioning are steadily accumulating. Out of the 90 newly approved drugs in 2016 (a 10% decrease from 2015), 25% are repositionings in terms of formulations, combinations, and indications^4^. Furthermore, drug repositioning reduces the incurred research and development (R&D) time and costs, as well as medication risks^8, 9^.

The recent developments confirm computational methods as powerful tools for drug repositioning:

- The trivialization/spread of omics analytical approaches have generated significant volumes of useful multi-omics data (genomics, proteomics, metabolomics, and others)^10, 11^.
- The ubiquity of digitalization in everyday life, including social media, has tremendously expanded the amplitude of the process of gathering data on drug-drug interactions and drug side-effects^12, 13^.
- The recent developments in physics, computer science, and computer engineering have created efficient methods and technologies for data exploration and mining, such as complex network analysis, machine learning, or deep learning^11, 14–18^.

Complex network analysis has proven to be a useful tool for predicting unaccounted drug-target interactions. Indeed, several state-of-the-art network-based computational drug repurposing approaches use data on confirmed drug-target interactions to predict new such interactions, thus leading to new repositioning hints^19, 20^. Some approaches build drug-drug similarity networks, where the similarity is defined based on transcriptional responses^21, 22^. These repositioning approaches analyze the network parameters and the node centrality distributions in either drug-drug or drug-target networks, using statistical analysis^10, 11, 23, 24^ and machine learning (i.e., graph convolutional networks)^25–28^ to link certain drugs to new pharmacological properties. However, conventional statistics can be misleading when used to predict extreme centrality values, such as degree and betweenness (which particularly indicate nodes/drugs with a high potential for repositioning)^29^.

To overcome these challenges, we developed a novel, network-based, computational approach to drug repositioning. To this end, we build a weighted *drug-drug network*, i.e., a complex network where the nodes are drugs, and the weighted links represent relationships between drugs, using information from the accurate DrugBank^30^. In our drug-drug similarity network (DDSN), a link is placed between two drugs if their interaction with at least one target is of the same type (either agonistic or antagonistic). The link weight represents the number of biological targets that interact in the same way with the two drugs. Figure 2 illustrates the building of the DDSN with information on drug-target interactions.

**Figure 2.**
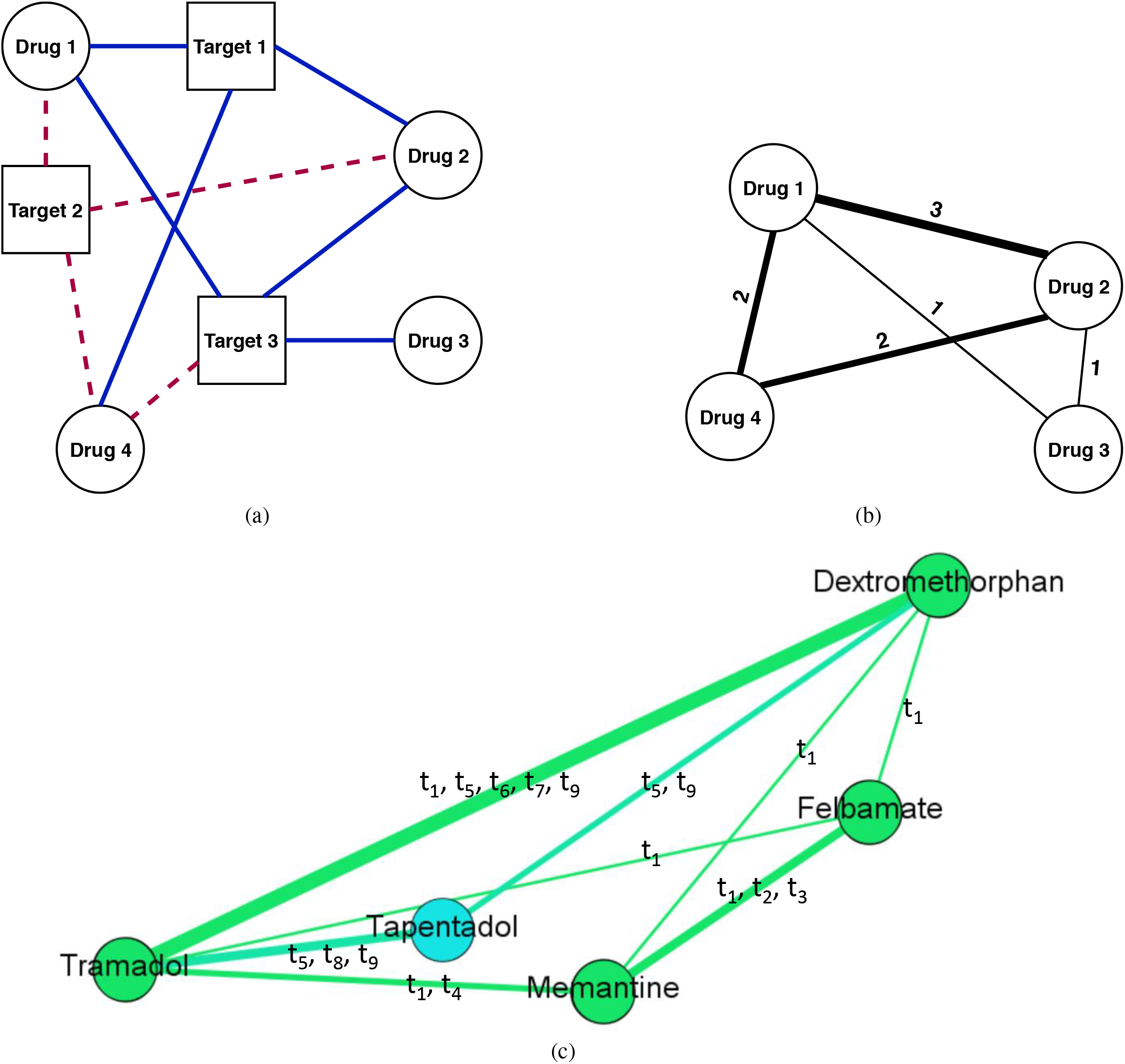
An illustrative example of using drug-target interaction information to build a weighted drug-drug similarity network. In panel (a), we consider the drug-target interactions between four drugs (i.e., round nodes labeled 1 to 4) and three targets (i.e., square nodes labeled 1 to 3). The dashed red links represent agonist drug-target interactions, whereas the solid blue links represent antagonist drug-target interactions. In panel (b), we show the DDSN corresponding to the interactions in (a). For instance, a link of weight 3 connects the nodes 1 and 2 because Drug 1 and Drug 2 interact in the same way for the three targets, i.e., agonist on Target 2 and antagonist on Targets 1 and 3. Also, a link with weight 2 connects Drug 2 and Drug 4 because they both interact agonistically on Target 2 and antagonistically on Target 1, but they do not interact in the same way with Target 3. In panel (c), we show a DDSN sub-network example, according to drug-target interactions from DrugBank 4.2, containing drugs Dextromethorphan, Felbamate, Tapentadol, Tramadol, and Memantine. We shape the link thickness according to the weight and specify the list of common targets for each link. The weight equals the number of targets in the list, where *t*_1_ = Glutamate receptor ionotropic NMDA 3A, *t*_2_ = Glutamate receptor ionotropic NMDA 2A, *t*_3_ = Glutamate receptor ionotropic NMDA 2B, *t*_4_ = Alpha-7 nicotinic cholinergic receptor subunit, *t*_5_ = Mu-type opioid receptor, *t*_6_ = Kappa-type opioid receptor, *t*_7_ = Delta-type opioid receptor, *t*_8_ = Sodium-dependent noradrenaline transporter, and *t*_9_ = Sodium-dependent serotonin transporter.

Our methodology for analyzing the drug-drug similarity network (DDSN) consists of the following steps:

1. Generate (using the Force Atlas 2 layout and modularity classes)^31, 32^ both topological clusters, and network communities.
2. Relate each cluster and each community to a pharmacological property or pharmacological action (i.e., label communities and clusters according to the dominant property or pharmacological action), using expert analysis.
3. Identify and select (by betweenness divided by degree, 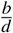) within each topological cluster/modularity class community, the top drugs not compliant with the cluster/community label.
4. Validate the hinted repositionings by searching the new versions of DrugBank, the electronic records containing the relevant scientific literature (for merely reconstructed repositionings), and by analyzing molecular docking parameters^33^ for previously unaccounted repositionings.

## Results

### Building the DDSN

We built our DDSN as a weighted graph *G* = (*V, E*) with |*V*| = *n* vertices (nodes) representing drugs and |*E*| = *n_E_* edges representing drug-drug similarity relationships based on drug-target interactions. The weight associated with an edge *e_ij_* ∈ *E* represents the degree of *target action similarity* between drugs *v_i_* and *v_j_*, and it is equal with the number of *common biological targets* for *v_i_* and *v_j_*. If *e_ij_* = 0, then there is no target similarity between *v_i_* and *v_j_*, therefore no edge between these nodes. A *common biological target* is a target *t_k_* ∈ *T* (*T* is the set of targets) on which drugs *v_i_* and *v_j_* act in the same way, either both agonistically or both antagonistically. Within this framework, we build the DDSN graph *G* using drug-target interaction information from Drug Bank 4.2^30^. We base our analysis on the largest connected component of the DDSN, consisting of |*V*| = 1008 drugs/nodes and |*E*| = 17963 links resulted from the analysis of the drug-target interactions with |*T*| = 516 targets.

To mine the DDSN topological complexity, we identified the drug clusters (or communities) using both the modularity^32^ and the force-directed, energy-based layout Force Atlas 2^31^ algorithms. The two clustering techniques are compatible^34^; however, the energy-based force-directed layout clustering offers more information about the relationship between clusters and acts as an efficient classifier^35^. In the case of DDSN, the clusters correspond to drug communities *C_x_*, *x* ∈ ℕ^*^, such that 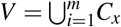. Using the constructed DDSN from Drug Bank 4.2 and expert analysis, we label each cluster according to its dominant property (i.e., the property that better describes the majority of drugs in the cluster – see supplementary file *SupplementaryDDSN.xlsx* for a detailed proof), which may represent a specific mechanism of pharmacologic action, a specifically targeted disease, or a targeted organ. Figure 3 illustrates the resulted DDSN, where the node colors identify the distinct modularity clusters. We assessed the ability of our method to uncover new repositionings by confronting our results with the latest (version 5.1.4) Drug Bank and with data compiled from interrogating scientific literature databases.

**Figure 3.**
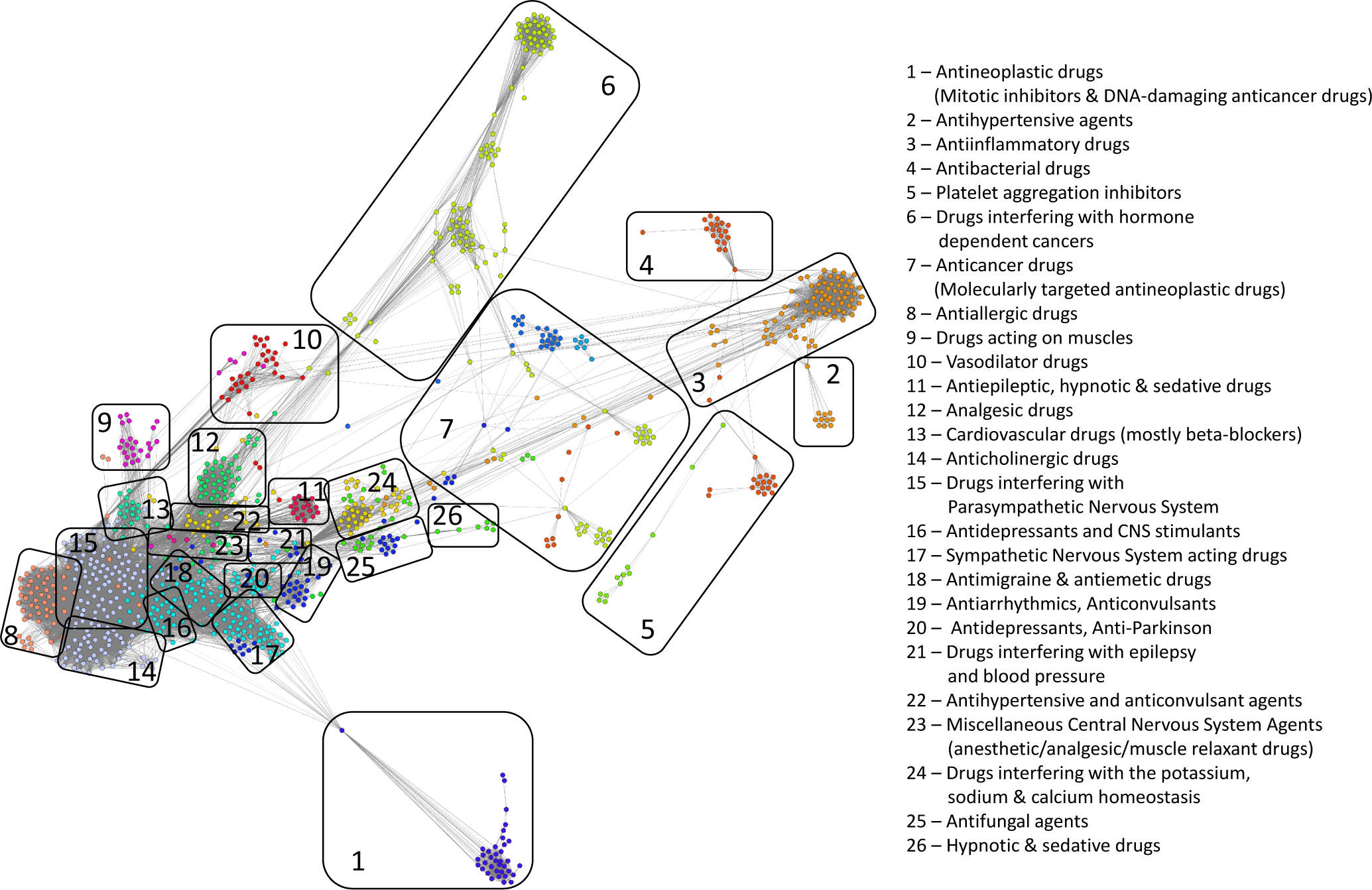
The drug-drug similarity network, where nodes represent drugs and links represent drug-drug similarity relationships based on drug-target interaction behavior. The layout is Force Atlas 2^31^ and the distinct node colors identify the modularity classes that define the drug clusters. We identify the 26 topological clusters with rounded rectangles and provide the functional descriptions for each of them.

When using network clustering, if a drug does not comply with the community/cluster label, then this indicates a possible repurposing^36^. We labeled the clusters using the drug properties listed by DrugBank or reported in the literature, such that the dominant property or properties (i.e., properties found in more than 50% of the drugs in the community) give the name of the community, as indicated in Tables 1 and 2. According to Tables 1 and 2 (column *Literature* [%]), our DDSN computational approach recovers/reconstructs a significant number of drug repurposings reported in the literature (see the supplementary file *SupplementaryDDSN.xlsx* for detailed confirmation literature lists, including some recent repurposing confirmations), namely 26.98% of the 1008 drugs in DDSN (last line in Table 2, summarizing the confirmation results).

**Table 1.** Confirmation of drug community properties and drug repurposing hints. Each table line is dedicated to a distinct topological community *C_x_* (with 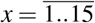), by specifying the dominant property(-ies) resulted from the pharmacological expert analysis (column *Properties*), the number of nodes/drugs in community *C_x_* (column *Nodes* [#]), the percentage of drugs with the properties confirmed by DrugBank (column *DrugBank* [%]), the percentage of drugs with the predicted properties confirmed by the literature (column *Literature* [%]), the percentage of drugs with not yet confirmed predicted properties (column *Not confirmed* [%]), and the drugs we propose for repositioning, representing predictions not confirmed yet but with non-zero betweeness/degree in the DDSN (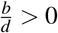, in column *Hints*).

| $C_x$ | Properties | Nodes [#] | DrugBank [%] | Literature [%] | Not confirmed [%] | Hints |
| --- | --- | --- | --- | --- | --- | --- |
| 1 | Antineoplastic (mitotic inhibitors and DNA-damaging) | 37 | 40.54 | 37.84 | 21.62 | Besifloxacin<br>Pefloxacin<br>Norfloxacin<br>Ofloxacin |
| 2 | Antihypertensive (sartans) | 10 | 100 | 0 | 0 | – |
| 3 | Anti-inflammatory | 84 | 65.48 | 28.57 | 5.95 | Glipizide |
| 4 | Antibacterial tetracyclines and Aminoglycosides | 20 | 95.00 | 0 | 5.00 | Plerixafor |
| 5 | Platelet aggregation inhibitor | 29 | 10.34 | 82.76 | 6.90 | – |
| 6 | Interfering with hormone-dependent cancers | 93 | 26.88 | 65.59 | 7.53 | Azelaic ac. |
| 7 | Anticancer (molecularly targeted) | 92 | 23.91 | 50.00 | 26.09 | Suramin<br>Acetohydroxamic ac.<br>Glyburide<br>Gliquidone<br>Tolbutamide |
| 8 | Anti-allergic | 51 | 86.27 | 11.76 | 1.96 | Butriptyline |
| 9 | Acting on muscles | 25 | 72.00 | 16.00 | 12.00 | – |
| 10 | Vasodilator | 37 | 48.65 | 24.32 | 27.03 | Tofisopam<br>Mefloquine<br>Oxtriphylline<br>Enprofylline<br>Roflumilast<br>Aminophylline |
| 11 | Antiepileptic, hypnotic, and sedative | 19 | 84.21 | 10.53 | 5.26 | Barbituric ac. deriv. |
| 12 | Analgesic and used in opiate withdrawal & side-effects | 46 | 89.13 | 8.70 | 2.17 | – |
| 13 | Antihypertensive, anti-arrhythmic, anti-angina (mostly beta-blockers) | 26 | 92.31 | 3.85 | 3.85 | – |
| 14 | Anticholinergic | 53 | 100 | 0 | 0 | – |
| 15 | Interfering with Parasympathetic Nervous System | 97 | 42.27 | 42.27 | 15.46 | Doxazosin<br>Terazosin<br>Prazosin<br>Paliperidone<br>Aripiprazole<br>Fenoldopam<br>Dapiprazole<br>Alfuzosin<br>Tamsulosin<br>Silodosin<br>Amisulpiride<br>Carphenazine<br>Acetophenazine |

**Table 2.** Confirmation of drug community properties and drug repurposing hints. Each table line is dedicated to a distinct topological community *C_x_* (with 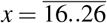), as well as the last line for the entire DDSN, by specifying the dominant property(-ies) resulted from the pharmacological expert analysis (column *Properties*), the number of nodes/drugs in community *C_x_* (column *Nodes* [#]), the percentage of drugs with the properties confirmed by DrugBank (column *DrugBank* [%]), the percentage of drugs with the predicted properties confirmed by the literature (column *Literature* [%]), the percentage of drugs with not yet confirmed predicted properties (column *Not confirmed* [%]), and the drugs we propose for repositioning, representing predictions not confirmed yet but with non-zero betweeness/degree in the DDSN (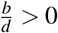, in column *Hints*).

| $C_x$ | Properties | Nodes [#] | DrugBank [%] | Literature [%] | Not confirmed [%] | Hints |
| --- | --- | --- | --- | --- | --- | --- |
| 16 | Antidepressant and Central Nervous System stimulant | 26 | 92.31 | 7.69 | 0 | – |
| 17 | Sympathetic Nervous System acting | 61 | 85.25 | 8.20 | 6.56 | – |
| 18 | Antimigraine and antiemetic | 26 | 42.31 | 26.92 | 30.77 | Captodiamine<br>Ropinirole<br>MDMA<br>Dofetilide<br>Rotigotine<br>L-DOPA |
| 19 | Antiarrhythmic and anticonvulsant | 24 | 66.67 | 12.50 | 20.83 | Acarbose<br>Hexylcaine |
| 20 | Antidepressant and anti-Parkinson | 21 | 57.14 | 14.29 | 28.57 | Quinidine<br>Propafenone<br>Cinchocaine<br>MMDA<br>Aprindine |
| 21 | Interfering with epilepsy and blood pressure | 12 | 41.67 | 25.00 | 33.33 | Miconazole<br>Quinidine barbiturate |
| 22 | Antihypertensive and anticonvulsant | 20 | 80.00 | 15.00 | 5.00 | – |
| 23 | Anesthetic, analgesic, and muscle relaxant | 19 | 73.68 | 5.26 | 21.05 | Halofantrine<br>Ibutilide<br>Pentolinium |
| 24 | Interfering with K, Na, Ca homeostasis | 51 | 50.98 | 13.73 | 35.29 | Progabide<br>Bethanidine<br>Ellagic ac.<br>Vigabatrin<br>Ethinamate |
| 25 | Antifungal | 22 | 59.09 | 9.09 | 31.82 | Meprobamate<br>Enflurane<br>Sevoflurane<br>Desflurane |
| 26 | Hypnotic and sedative | 7 | 100 | 0 | 0 | – |
| All | – | 1008 | 59.52 | 26.98 | 13.49 | – |

### Illustrative examples of reconstructed drug repositionings

Here, we present a few illustrative examples of reconstructed drug repositionings, as confirmed by recent and very recent literature. We provide the entire list of drug repositionings we recovered with the DDSN method and the references that prove them as such in the supplementary file *SupplementaryDDSN.xlsx*.

### Reconstructed repurposings as antineoplastic agents

The topological community 1 (i.e., *C*_1_) consists of antineoplastic drugs, mostly mitotic inhibitors (e.g., Etoposide, Teniposide, Vincristine, Vinorelbine) and DNA-damaging anticancer drugs (e.g., Doxorubicin, Valrubicin, Mitoxantrone). This community also contains fluoroquinolone antibiotics (targeting the alpha subunits of two types of bacterial topoisomerase II enzymes, namely DNA gyrase and DNA topoisomerase 4) and a few other drugs. However, DrugBank does not confirm the anticancer effects of some drugs within topological *C*_1_, yet the literature confirms them as such. For example, Colchicine, which is currently used based on its anti-inflammatory effects as an antigout drug, exhibits anticancer effects^37^; Podofilox, a drug for topical treatment of external genital warts, is a potent cytotoxic agent in chronic lymphocytic leukemia (CLL)^38^; for some fluoroquinolone drugs the literature reports anticancer effects (e.g., Enoxacin^39^, Ciprofloxacin^40^, Moxifloxacin^41^, Gatifloxacin^42^). In Figure 4, we show a zoomed detail from our DDSN, by highlighting the presence of Colchicine, Podofilox, Enoxacin, Ciprofloxacin, Moxifloxacin, Gatifloxacin in *C*_1_; such topological placement suggests their antineoplastic effect.

**Figure 4.**
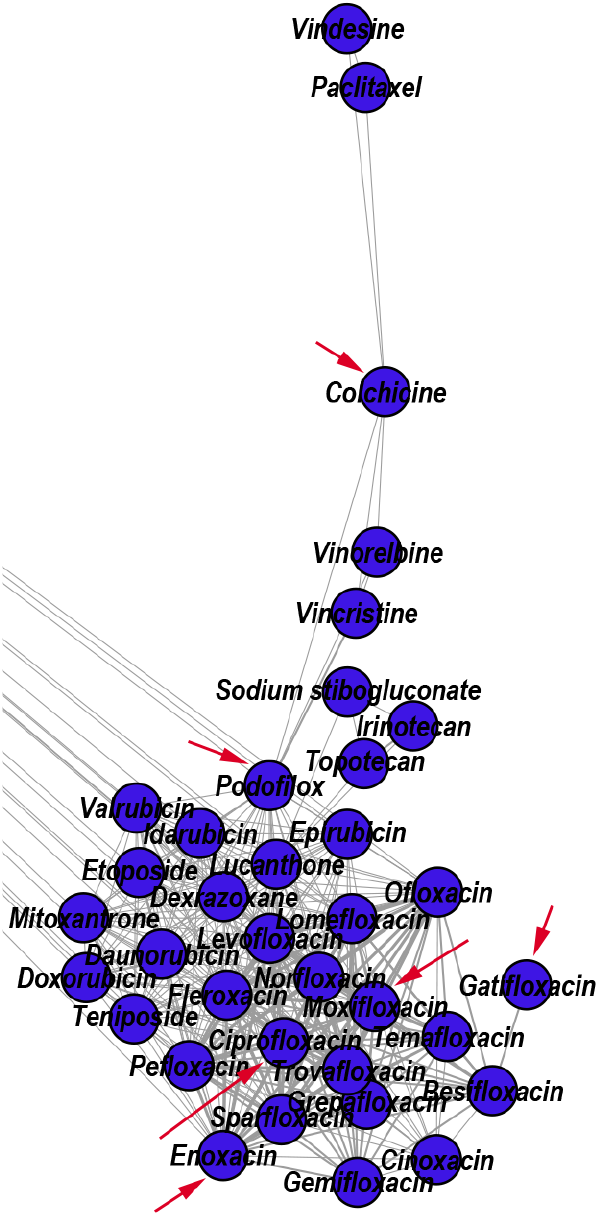
Zoomed DDSN detail of Community 1 (*C*_1_, Antineoplastic drugs – Mitotic inhibitors & DNA-damaging anticancer drugs). The red arrows indicate the reconstructed drug repositionings: Colchicine (antigout drug), Podofilox (topical antiviral), and Enoxacin, Ciprofloxacin, Moxifloxacin, Gatifloxacin (fluoroquinolone antibiotics).

The topological community *C*_6_ consists of anticancer drugs that target hormone-dependent organs (i.e., ovary, endometrium, vagina, cervix, and prostate). In this community, Progesterone has the highest value of betweenness/degree ratio, and the DrugBank database does not indicate its anticancer property. Although there are extensive epidemiological studies that link the long-term Progesterone use in oral contraceptives to breast cancer risk, this link is strengthened or weakened by various parameters, such as body weight, age, duration of use^43^, parity, age at first birth, breastfeeding, and age at menarche^44^. However, J.C. Leo et al. determined the whole genomic effect of Progesterone in PR-transfected MDA-MB-231 cells and found that Progesterone suppressed the expression of genes involved in cell proliferation and metastasis, concluding that Progesterone can exert a strong anticancer effect in hormone-independent breast cancer following Progesterone receptor (PR) reactivation^45^. Quinacrine is an antiprotozoal drug that exhibits an anticancer effect in breast cancer, because it produces apoptosis by blocking cells in S-phase, induces DNA damage, and inhibits topoisomerase activity^46^; indeed, reference^47^ recommends the clinical trial test of Quinacrine for the treatment of patients with androgen-independent prostate cancer. The antineoplastic drug Mimosineattenuates cell proliferation of prostate carcinoma cells in vitro^48^. Figure 5 provides a zoomed detail (i.e., focused view) of the DDSN that highlights the presence of Mimosine (an experimental antineoplastic which inhibits DNA replication) in *C*_6_; this indicates that Mimosine has effects in hormone-dependent cancers.

**Figure 5.**
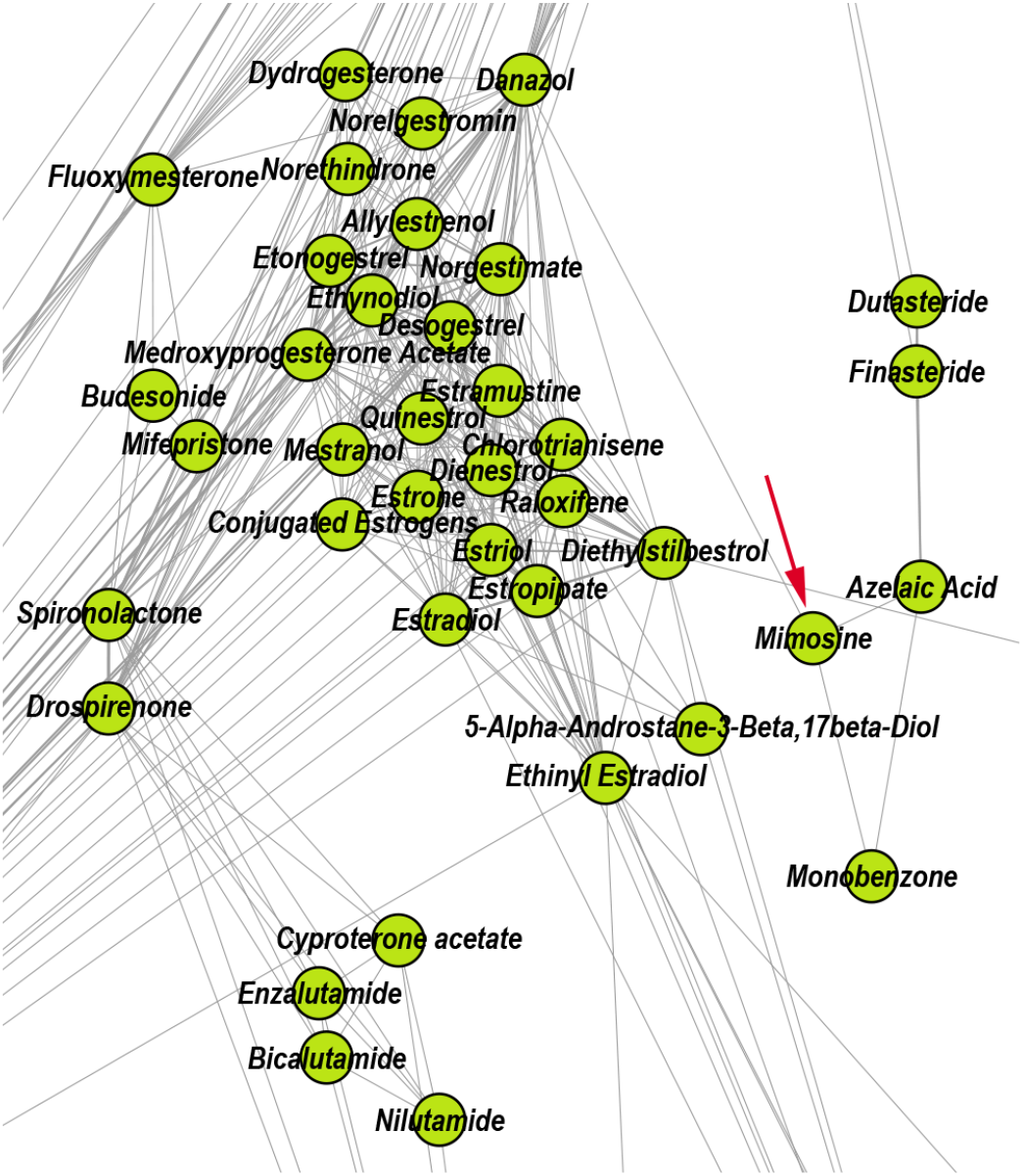
Zoomed DDSN detail of community *C*_6_ (Drugs interfering with hormone dependent cancers). The red arrow indicate the reconstructed drug repositioning: Mimosine – an experimental antineoplastic which inhibits DNA replication – also has effects in cancers affecting hormone-dependent organs.

### Reconstructed repurposings as anti-inflammatory drugs

According to the properties listed in DrugBank, the topological community *C*_3_ includes drugs that exert anti-inflammatory effects via different mechanisms: non-steroidal anti-inflammatory drugs (e.g., Diclofenac, Ibuprofen, and Acetylsalicylic acid), the antirheumatic agent Auranofin, hypoglycemic drugs (e.g., Rosiglitazone, Troglitazone), and the antihypertensive drug Telmisartan. Moreover, the literature confirms that 28.57% of drugs within this community also present anti-inflammatory effects, even if they are not listed as anti-inflammatory in DrugBank. Here, we present the example of the versatile molecule of Fenofibrate, which reduces the systemic inflammation independent of its lipid regulation effects, with cardiovascular benefits in high-risk^49^ and rheumatoid arthritis patients^50^. Another illustrative example is that of Amiloride, which inhibits the activation of the dendritic cells and ameliorates the inflammation besides its diuretic effects, thus having benefits for hypertensive patients^51^. Figure 6 shows a zoomed DDSN detail, which highlights the presence of Fenofibrate and Amiloride in *C*_3_; this may indicate that the highlighted drugs also have anti-inflammatory effects.

**Figure 6.**
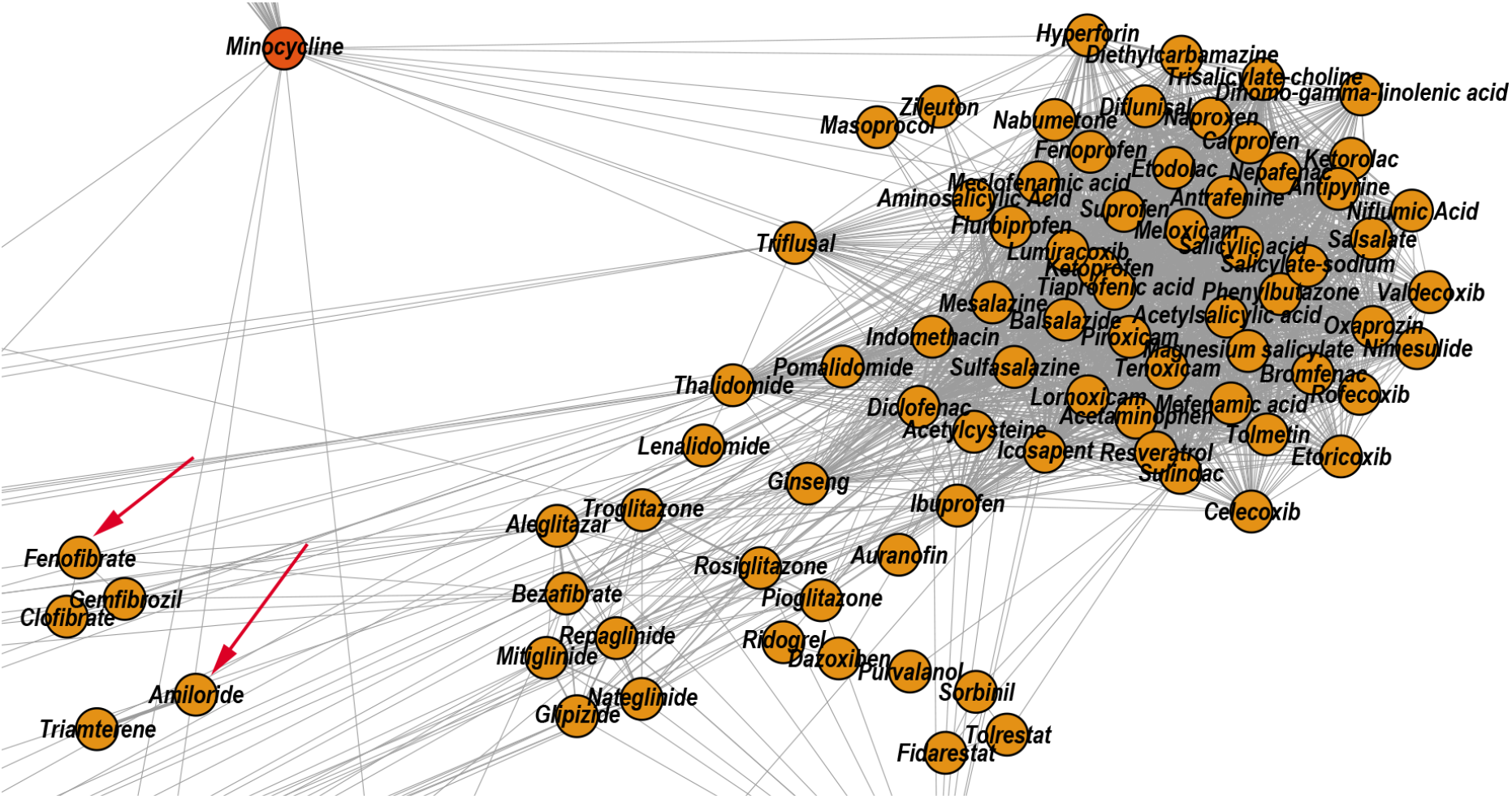
Zoomed DDSN detail of community *C*_3_ (Anti-inflammatory drugs). The red arrows indicate the reconstructed drug repositionings as anti-inflammatory drugs: Fenofibrate (a lipid modifying drug) and Amiloride (a diuretic).

### Reconstructed repurposings as antifungal drugs

The topological community *C*_25_ includes 22 drugs. According to DrugBank, 13 out of these 22 drugs have antifungal properties, and 9 drugs act on the central nervous system (i.e., general anesthetics, sedative-hypnotics, and antiepileptics). DrugBank lists Isoflurane and Methoxyflurane as general anesthetic drugs. However, A. Giorgi et al. performed in vitro tests to investigate the antibacterial and antifungal effects of common anesthetic gases, and they found that Methoxyflurane and Isoflurane have excellent inhibitory effects on cultures of Klebsiella pneumoniae and Candida albicans^52^. Using *in vitro* experiments, V.M. Barodka et al. also found that the liquid formulation of Isoflurane has a better ani-Candida activity in comparison with the antifungal Amphotericin B^53^. Figure 7 shows a zoomed DDSN detail that highlights the presence of Isoflurane and Methoxyflurane in *C*_25_; this indicates that the highlighted drugs may also have antifungal effects.

**Figure 7.**
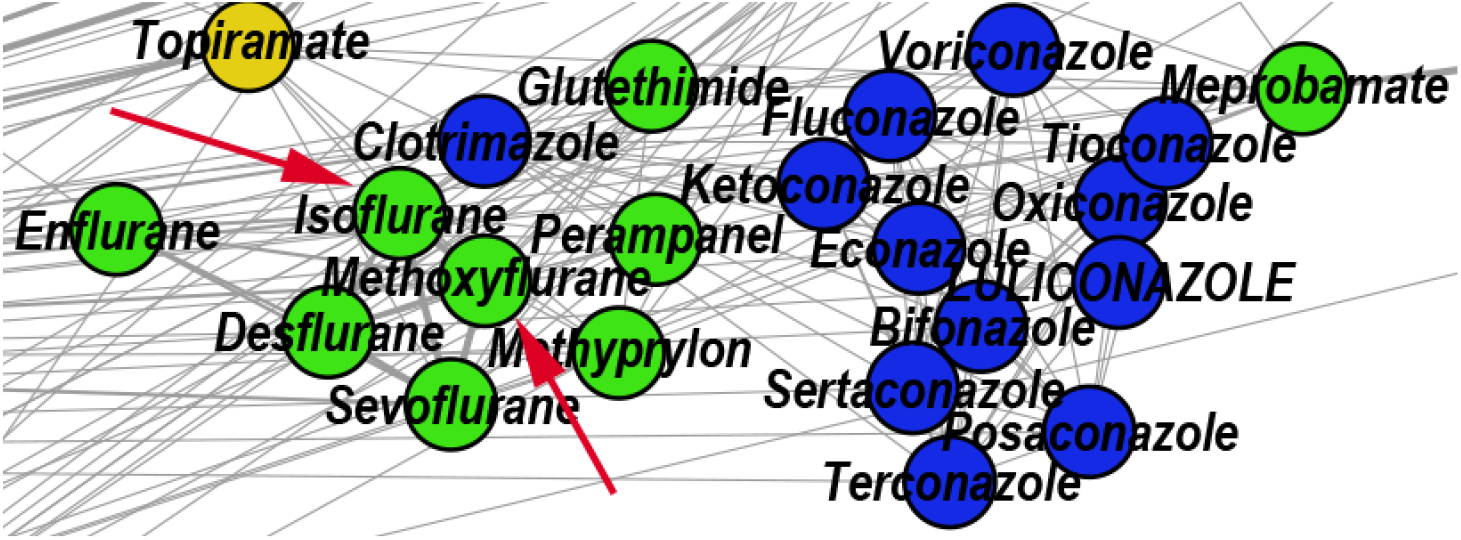
Zoomed DDSN detail of community *C*_25_ (Antifungal agents). The red arrows indicate the reconstructed drug repositionings: Isoflurane and Methoxyflurane (known as general anesthetic drugs) also have antifungal effects.

### DDSN analysis and repositioning hints

In our characterization of drug-drug similarity networks, a high degree node represents a drug with already documented multiple properties. Also, a high betweenness (i.e., the ability to connect network communities) indicates the drug’s propensity for multiple pharmacological functions. By this logic, the high-betweenness, high-degree nodes may have reached their full repositioning potential, whereas the high betweenness, low degree nodes (characterized by high betweenness/degree value 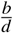) may indicate a significant repositioning potential. However, predicting such high-value cases of degree *d*, weighted degree *d_w_*, betweenness *b*, and betweenness/degree 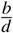 is difficult because the corresponding distributions are fat-tailed^54^. Although all the estimated DDSN centralities are following a power-law distribution (see Figure 8), the betweenness/degree 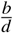 is the most stable parameter and, hence, the most reliable indicator of multiple drug properties.

**Figure 8.**
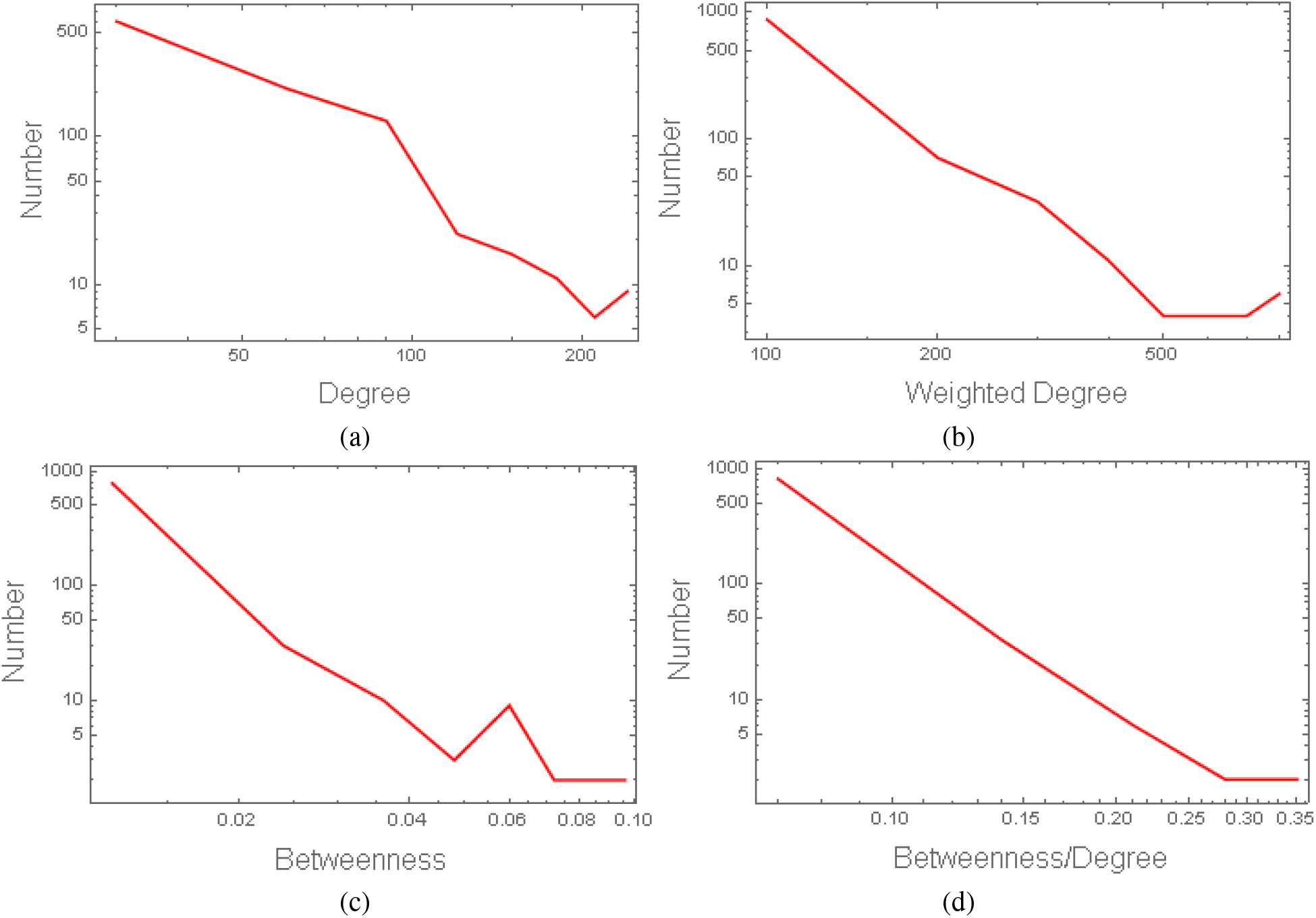
Power-law distributions of centrality parameters in the drug-drug similarity network (DDSN): (a) degree *d*, (b) weighted degree *d_w_*, (c) betweenness *b*, and (c) betweenness/degree 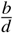. We represent the distributions according to the guidelines in^55^, by using 8 linearly spaced bins for each centrality. The fitting analysis using the *Powerlaw* package in Python^55^ indicates the following values for the distribution slope *α* and cutoff point *x_min_*, respectively: 3.436 and 53 for *d*, 2.598 and 64 for *d_w_*, 2.201 and 0.008 for *b*, 3.093 and 0.088 for 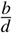. The graphical representations of these centrality distributions show that the betweenness/degree 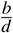 is the most stable parameter; therefore it is the most reliable indicator of multiple drug properties.

To explore the capability of 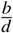 to predict the multiple drug properties, we exploit the community structure of DDSN by following a two-step approach.

1. We uncover the relevant drug properties by generating network communities *C_x_* with 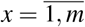 (*m* = 26 in our DDSN). Then, using expert analysis, we assign a dominant property to each community. Figure 3 illustrates the 26 DDSN communities as well as their dominant functionality. The dominant community property can be a pharmacological mechanism, a targeted disease, or a targeted organ. For instance, the community 1 (*C*_1_) consists of antineoplastic drugs which act as mitotic inhibitors and DNA damaging agents; Community 13 (*C*_13_) consists of cardiovascular drugs (antihypertensive, anti-arrhythmic, and anti-angina drugs), mostly beta-blockers.
2. In each cluster *C_x_*, we identify the top *t* drugs according to their 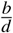 values. From these selected drugs, 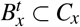, some stand out by not sharing the community property or properties, and thus, can be repositioned as such. To this end, for 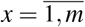 eliminated from 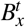 the drugs whose repurposings were already confirmed (i.e., performed by others and found in the recent literature), thus producing *m* = 26 lists of repurposing hints yet to be confirmed by *in silico*, *in vitro*, and *in vivo* experiments, 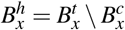. Table 3 presents the lists of 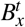 drugs for *t* = 5 and *x* = 26 (i.e., the top 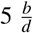 drugs in each community). We chose *t* = 5 to provide a reasonable amount of eloquent information in Table 3; we provide the entire *B_x_* sets in the supplementary file *SupplementaryDDSN.xlsx*.

**Table 3.** Top 5 drugs (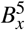 with 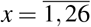) according to their 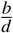 values, for each of the 26 DDSN communities/clusters (*C_x_*). The properties of drugs written in regular fonts match the properties of their respective communities (according to the Drug Bank). The properties of italicized drugs do not match all the properties of their respective communities, but the latest literature confirms them (drugs in regular fonts and italics pertain to 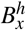). The properties of the drugs written in bold do not match the community properties, and the literature did not confirm them yet; this situation leads to new drug repositioning hints (i.e., the 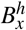 drugs). The positions marked with ‘–’ correspond to drugs with 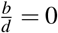.

| $C_x \backslash B_x^5$ | 1 | 2 | 3 | 4 | 5 |
| --- | --- | --- | --- | --- | --- |
| 1 | Amsacrine | *Colchicine* | *Podofilox* | *Lucanthone* | **Besifloxacin** |
| 2 | – | – | – | – | – |
| 3 | *Amiloride* | *Marimastat* | Diclofenac | Thalidomide | Telmisartan |
| 4 | Minocycline | Framycetin | Amikacin<br>Tobramycin<br>Netilmicin | Doxycycline<br>Clomocycline<br>Oxytetracycline | – |
| 5 | Treprostinil | Iloprost | *Captopril* | *Bimatoprost* | *Candoxatril* |
| 6 | *Progesterone* | *Mimosine* | *Fluticasone propionate* | *Danazol* | *Spironolactone* |
| 7 | Vandetanib | *Dalteparin* | *Dehydroepiandrosterone* | *Amlexanox* | *Atorvastatin* |
| 8 | Olopatadine | Terfenadine | Flunarizine | Astemizole | Epinastine |
| 9 | Succinylcholine | Carbachol | Decamethonium | Pilocarpine | Cevimeline |
| 10 | *Nicotine* | *Melatonin* | Amrinone | Dipyridamole | *Naloxone* |
| 11 | *Quinine* | Phenobarbital<br>Secobarbital<br>Pentobarbital | Barbital<br>Hexobarbital<br>Aprobarbital | – | – |
| 12 | *Nimodipine* | Adenosine | Drotaverine | Pentazocine | *Loperamide* |
| 13 | *Ketotifen* | Amiodarone | Sotalol | Bevantolol | Penbutolol |
| 14 | Disopyramide | Scopolamine | Ethopropazine | Paroxetine | Rocuronium |
| 15 | Minaprine | Amitriptyline | *Agomelatine* | Orphenadrine | Imipramine |
| 16 | Cocaine | *Chloroprocaine* | *Procaine* | Phenermine | Milnacipran |
| 17 | Epinephrine | 4-Methoxyamphetamine | Pseudoephedrine | Ephedra | Methamphetamine |
| 18 | *Ginkgo biloba* | **Captodiame** | *Cisapride* | *Bromocriptine* | *Carteolol* |
| 19 | **Acarbose** | Lidocaine | Mexiletine | *Etomidate* | Flecainide |
| 20 | Phenelzine | *Agmatine* | **Quinidine**<br>**Propafenone** | Ephedrine | *Amphetamine* |
| 21 | Zonisamide | **Miconazole** | *Ethanol* | **Quinidine barbiturate** | – |
| 22 | *Felodipine* | *Bepridil* | *Verapamil* | *Dextromethorphan* | *Amlodipine* |
| 23 | Halothane | **Halofantrine** | Tramadol | *Ibutilide* | Tubocurarine |
| 24 | *Thiamylal* | *Valproic Acid* | **Progabide** | **Bethanidine** | *Topiramate* |
| 25 | **Meprobamate** | **Enflurane** | Tioconazole | Clotrimazole | *Methoxyflurane*<br>**Isoflurane**<br>**Sevoflurane** |
| 26 | Flunitrazepam | Eszopiclone | – | – | – |

To facilitate the visual identification of the repositioning hints, in Figure 9, we shape the size of the nodes of our DDSN representation according to the magnitude of the 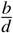 values. We also identify, by arrows, the top 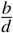 nodes (i.e., drugs) in their respective communities, by indicating their community id. Table 3 shows that 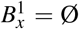 for all *x* except 19 and 25 (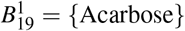 and 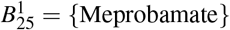), therefore – besides the corresponding community number – we expressly point Acarbose and Meprobamate in Figure 9.

**Figure 9.**
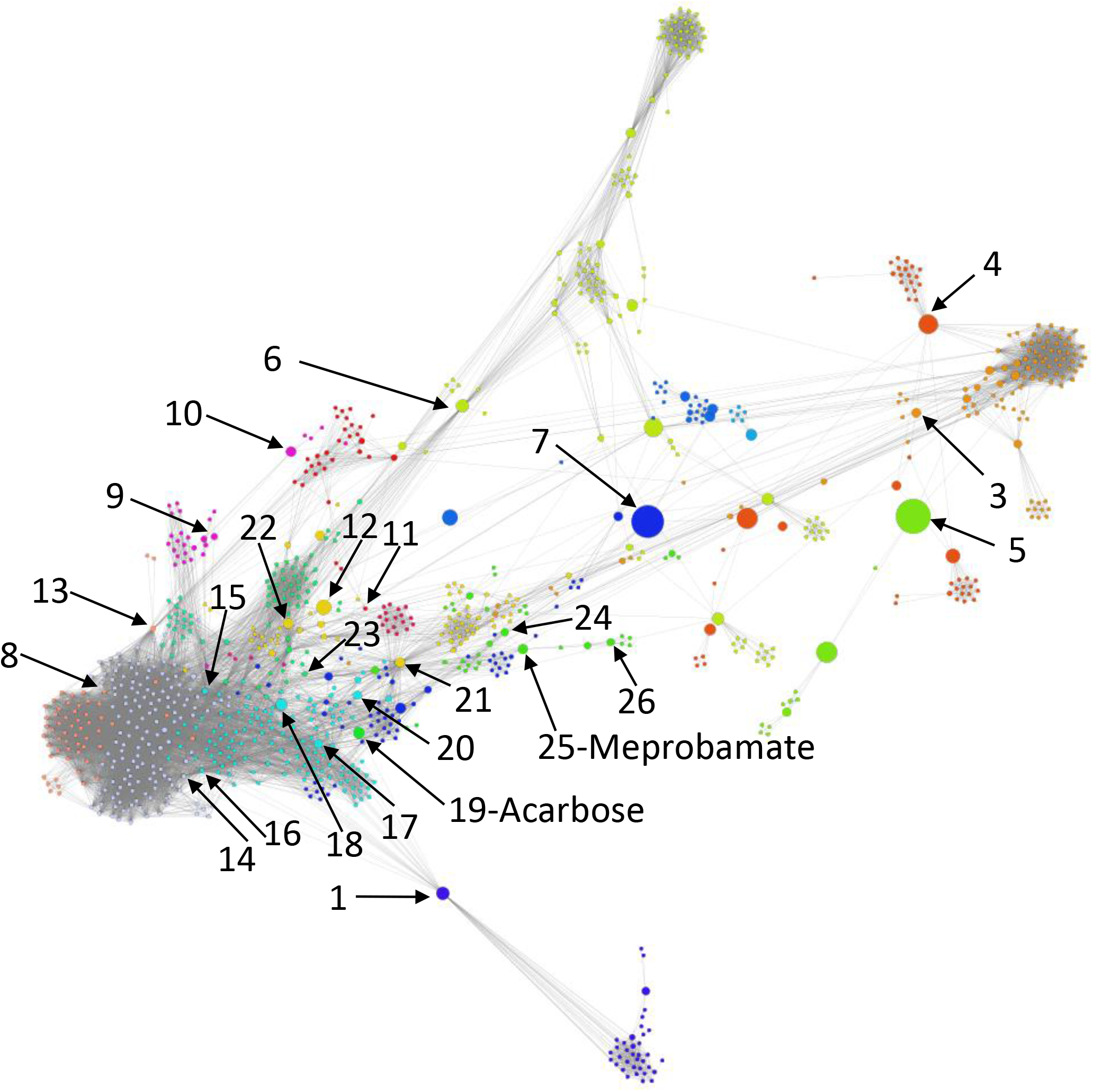
Drug-drug similarity network (DDSN), based on drug-target interactions, where node sizes represent their 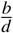 values. The arrows indicate the top 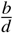 node in each community (for community 2 there is no top node because all drugs have 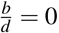). The community index identifies each top 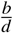 node, excepting Meprobamate (top 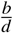 in community 25) and Acarbose (community 19), because these drugs (apparently) do not have their community’s property; this indicates Meprobamate as antifungal (i.e., the property of community 25) and Acarbose as antiarrhythmic, anticonvulsant (i.e., the properties of community 19).

The high percentage of database and literature confirmations of our pharmacological properties predictions are highlighting the robustness of our repurposing method. In the supplementary file *SupplementaryDDSN.xlsx*, we show that the confirmation rate 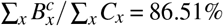. Table 3, presents a similar situation, with only a few unconfirmed drug properties (these repurposing hints 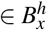 are in bold).

Our data indicate two top 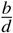 drugs: Meprobamate, in the *C*_25_ antifungal drugs community, and Acarbose, in the *C*_19_ (Antiarrhythmics and Anticonvulsants) community. Both repositionings refer to properties currently unaccounted in the DrugBank version 5.1.4 and the scientific literature we have screened (Table 3 and Figure 9). Meprobamate is a hypnotic, sedative, and mild muscle-relaxing drug, with no reported activities on the antifungal drug targets; thus, the antifungal activities of Meprobamate are not yet investigated in silico (with molecular docking), in vitro, or in vivo. Acarbose is a hypoglycemic drug, with no reported nor investigated antiarrhythmic and anticonvulsant properties. At the same time, one should also consider repurposing hints for drugs that have a high 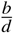, when the highest 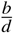 values correspond to drugs already confirmed with the community property. For example, Azelaic acid has the highest 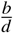 across the not confirmed drugs in *C*_6_.

### Repurposing hints testing

Molecular docking uses the target and ligand structures to predict the lead compound or repurpose drugs for different therapeutic purposes. The molecular docking tools predict the binding affinities, the preferred poses, and the interactions of the ligand-receptor complex with minimum free energy. In this paper, we use the AutoDock 4.2.6 software suite^56^, which consists of automated docking tools for predicting the binding of small ligands (i.e., drugs) to a macromolecule with an established 3D structure (i.e., target). The AutoDock semi-empirical free energy force field predicts the binding energy by considering complex energetic evaluations of bound and unbound forms of the ligand and the target, as well as an estimate of the conformational entropy lost upon binding.

According to the methodology in section *Methods* (subsection *Molecular docking for repurposing testing*), we verify the predicted properties of repurposing hints, by performing molecular docking not only for the hinted drugs but also for the reference drugs (typical drugs having the predicted property) and for some drugs with little probability of having the expected property. This way, we facilitate the comparison between the interaction of the hinted drug with the biological targets relevant for the tested property and the interactions of the reference drugs with the same targets.

Following the methodology in section *Methods*, subsection *Molecular docking for repurposing testing*, we first consider the the property *ϕ* as the anticancer effect with *x* = 6 (corresponding to community *C*_6_), and second *ϕ* as the antifungal effect with *x* = 25 (community *C*_25_). As such, we test the repurposing hints 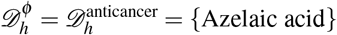 and 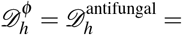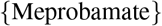.

Accordingly, we define the anticancer reference drug from *C*_6_ as 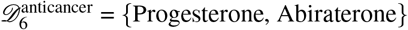, no anticancer reference drug outside *C*_6_ (i.e., 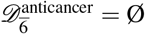), and two reference drugs that have a low probability of anticancer effects 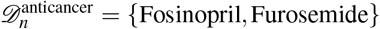 (Fosinopril is an antihypertensive and Furosemide is a diuretic). Here, we test the interaction between the hinted and reference drugs with the targets from DrugBank associated with anticancer drugs in 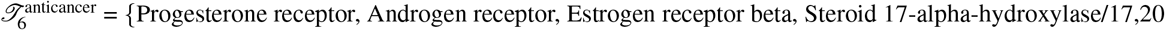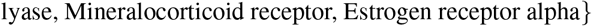.

Similarly, we consider the antifungal references in *C*_25_ as 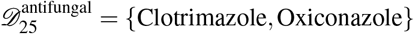, and outside *C*_25_ as 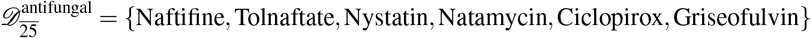. The reference drugs with little probability of having antifungal properties are 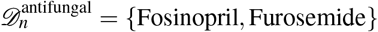. We test the interactions between the hinted and reference drugs with DrugBank antifungal-related targets linked to drugs in *C*_25_ and drugs not in *C*_25_, respectively 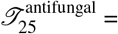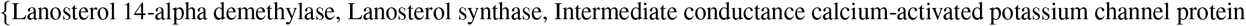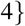, and 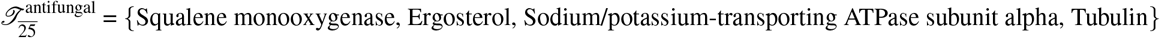.

Figure 10 shows the summary of interactions resulted from the molecular docking analysis of the drug-target pairs generated with equations 6, 7, and 8 (section *Methods*, subsection *Testing procedure*) for the hint 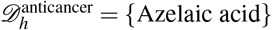. For the hint and the reference drugs 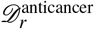, we represent the interactions with the targets 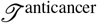 as the number of amino acids from the target interacting with the drug molecule (the maximum is 21). We provide the details related to the molecular docking simulations in the supplementary information material *SupplementaryInformation.pdf* – Tables S1-S6 and Figures S1-S6.

**Figure 10.**
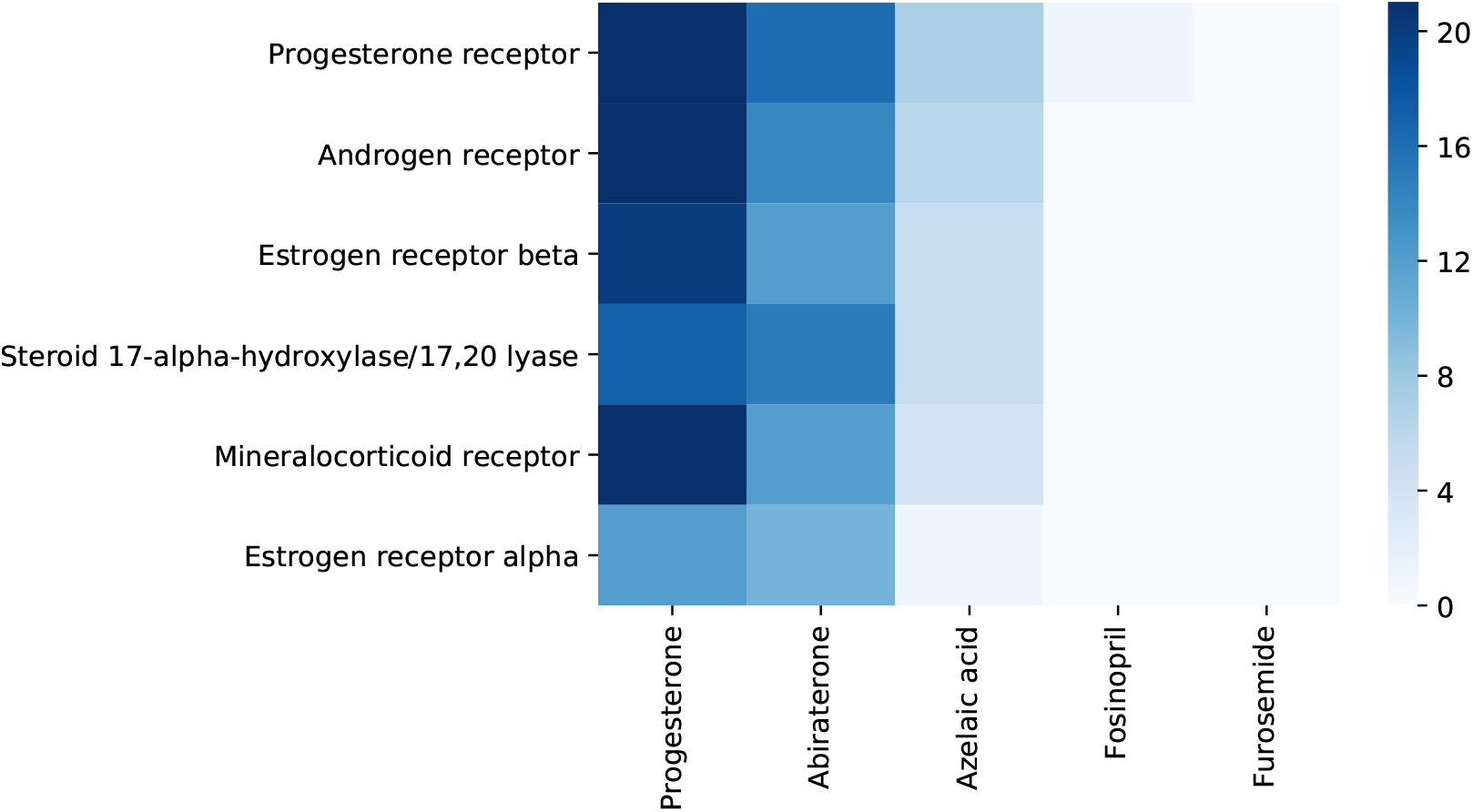
Synthesis of interactions resulted from running molecular docking on the drug-target pairs for 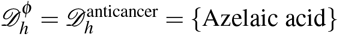. In the left part of the heatmap, we present the interactions between the relevant targets 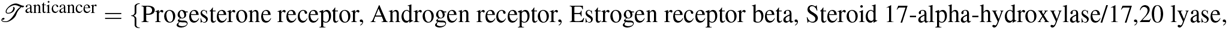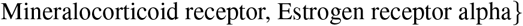 and the reference drugs 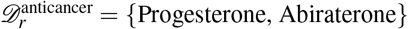. In the right part of the heatmap, we present the interactions between the relevant targets 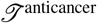 and the tested drugs 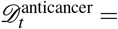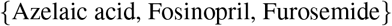. We summarize the interactions with the targets 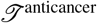 as the number of amino acids from the target interacting with the drug molecule (from 0 to the maximum number in our experiments, namely 21). The heatmap representation indicates interactions between 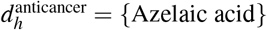 and almost all the targets from 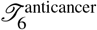. For the drugs in 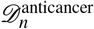. namely Fosinopril and Furosemide, there is no interaction with the targets from 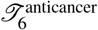.

Figure 11 presents the summary of interactions resulted from the molecular docking analysis of the drug-target pairs generated with equations 6, 7, and 8 (see section *Methods*, subsection *Testing procedure*), for the hint 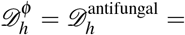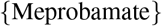. For the reference drugs 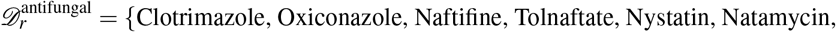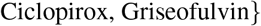 the interaction is represented as the number of amino acids in the target interacting with the drug molecule (the maximum in our molecular docking experiments is 24). Because 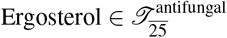 has a steroidal chemical structure, instead of the number of amino acids, we represent interaction strength as the number of hydrophobic alkyl/alkyl interactions. For the tested drugs 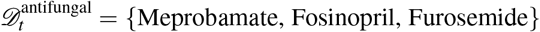, we represent the interaction as the number of amino acids from the target (or hydrophobic alkyl/alkyl interactions for Ergosterol) interacting *in the same way* with both the tested drug 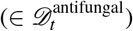 and at least one reference drug 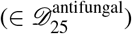. We provide the detailed description of the molecular docking simulations for Meprobamate in the supplementary information material *SupplementaryInformation.pdf* – Tables S7-S13 and Figures S7-S13. The results confirm the interactions between 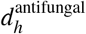 (i.e., Meprobamate) and almost all the targets from both 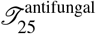 and 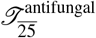. Conversely, for the drugs in 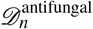, there is no relevant interaction with any arget from 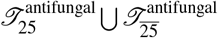.

**Figure 11.**
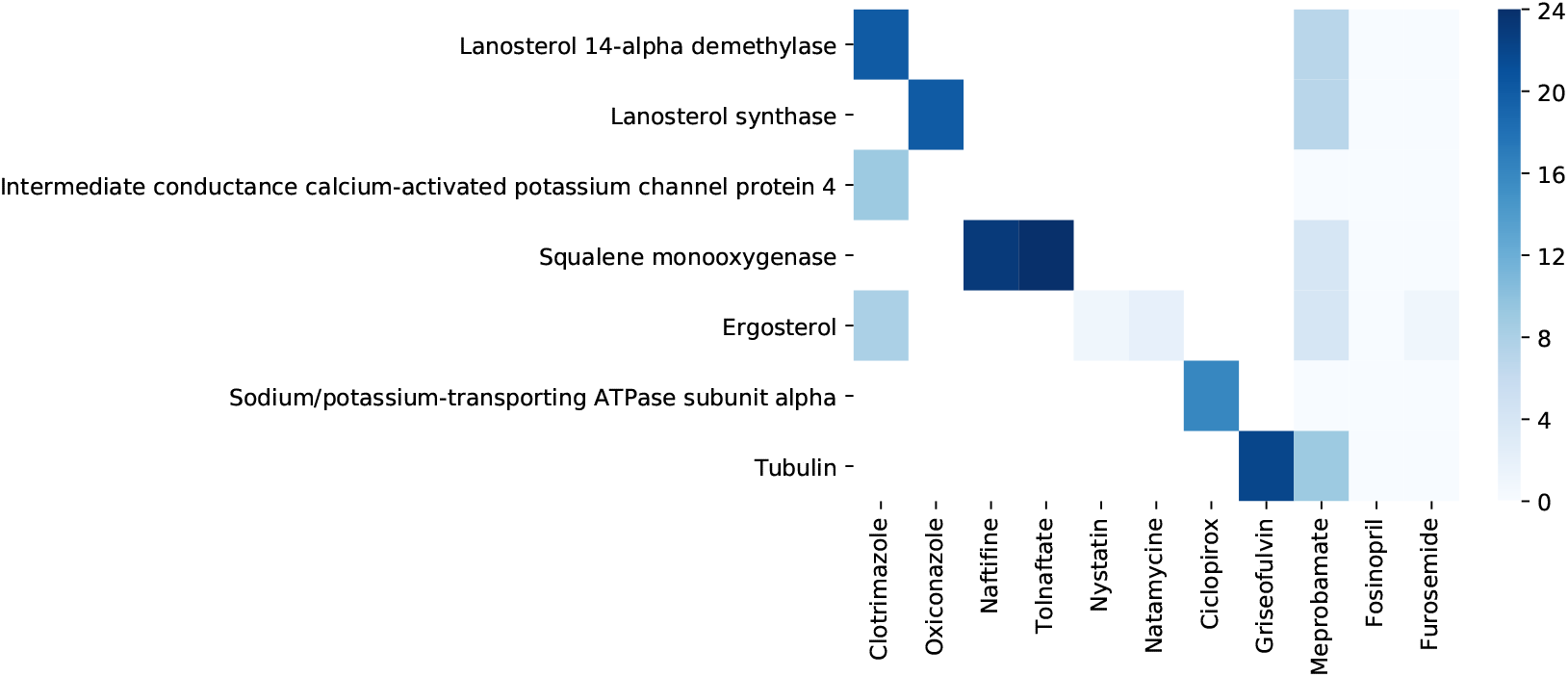
Synthesis of interactions resulted from running molecular docking on the drug-target pairs for 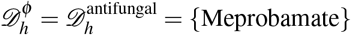. In the left part of the heatmap, we present the interactions between the relevant targets 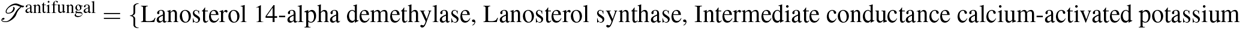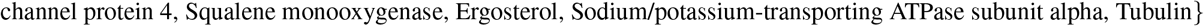 and the reference drugs 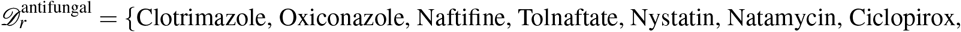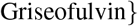. We only test the reference drugs and targets pairs that interact according to DrugBank; all the other pairs are white in our representation because they are not tested. In the right part of the heatmap, we present the interactions between the relevant targets 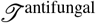 and the tested drugs 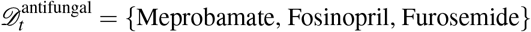. We summarize the interactions with the targets 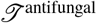 as the number of amino acids from the target interacting with the drug molecule (from 0 to the maximum number in our experiments, namely 24). In the case of 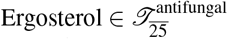, instead of the number of amino acids, we count the number of hydrophobic alkyl/alkyl interactions because this target has a steroidal chemical structure. The heatmap representation indicates interactions between 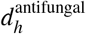 (i.e., Meprobamate) and almost all the targets from both 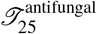 and 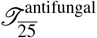. For the drugs in 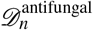, there is no relevant interaction with any target from 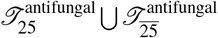.

## Discussion

Drug repurposing represents a promising strategy to accelerate drug discovery in sensitive areas of nowadays medicine, such as antibacterial resistance, complex life-threatening diseases (e.g., cancer), or rare diseases. In this paper, we describe a novel weighted drug-drug similarity network whose weights encode the existing known relationships among drugs (i.e., quantifies the number of biological targets shared by two drugs irrespective of the agonist or antagonist effect).

Then, we demonstrate that the ratio between node betweenness and node degree (i.e., a criterion of combined network metrics) can indicate the drug repositioning candidates better than considering simple network metrics (e.g., degree, weighted degree, betweenness). Indeed, the power-law distributions in Figure 8 suggest that our DDSN is a complex system; thus, the conventional statistical analysis of the DDSN can be misleading. Consequently, we introduce a different approach to deciphering the emerging hidden higher-order functional interactions (i.e., interactions that span multiple orders of magnitude and involve multiple nodes) by visualizing and analyzing the community structure in DDSN and determining the culprits (for such unknown functionalities) through combined network metrics criterion. We use the force-directed energy layout Force Atlas 2 to generate network clusters of drugs^31^ because it emulates the emerging processes of a complex system. More precisely, the force-directed based network layouts use micro-scale interactions (i.e., adjacent nodes attract and non-adjacent nodes repulse) to generate an emergent behavior at the macro-scale (i.e., topological clusters). Once we identify communities, the combined network metrics criterion selects the drug repositioning most likely candidates. Specifically, our weighted drug-drug network analysis encodes not only information about how pairs of drugs interact with biological targets, but also reveals the unknown functional relationship between drugs, such as the unknown effects on the activation/inhibition of a chemical pathway or cellular behavior. We used a similar methodology – underpinned by force-directed layout clustering – to analyze the fundamentally different structures represented by the drug-drug interaction networks (i.e., the DDIN interactome^36, 57^).

As presented in the literature, other computational methods allow for predicting new targets for existing drugs and are currently applied for drug repositioning or explaining off-target effects^58^. Indeed, the computer-assisted methods for predicting alternative targets or similar binding sites are rapidly evolving by making use of the enormous amount of drug data^59^. For instance, Mayr et al. found that the methods based on deep learning are much better than all other in silico approaches, sometimes matching the performance of wet-lab experiments^27^.

Molecular docking represents an alternative, in silico simulation approach to drug discovery, which models the physical interaction between a drug molecule and a target (or a set of targets). With molecular docking, we estimate the free energy values of the molecular interactions to offer a good approximation for the conformation and orientation of the ligand into the protein cavity^60^. Along with many available molecular docking models, DOCK^61^ is a dedicated software tool used in drug repurposing. For example, R. L. Des Jarlais et al. used the Dock computer algorithm to find that haloperidol inhibits HIV-1 and HIV-2 proteases^62^. However, molecular docking cannot work efficiently, unless we have some robust repositioning hints; otherwise, the search space for drug repositionings would be exponentially big. To this end, the methodology proposed in this paper provides strong drug-target interaction hints, such that we can build large-scale drug-target interaction profiles^63^. Our approach integrates the molecular docking with complex networks to hint new pharmacological properties by identifying new sets of biological targets on which the drug acts.

As indicated by Yvonne Martin et al.^64^, the paradigm of chemical similarity – which holds that structurally similar drug molecules exert similar biological effects – cannot fully explain the biological behavior of drugs. They found that only 30% of compounds similar to a particular active compound are themselves active (the compounds are structurally similar if the Tanimoto coefficient is ≥ 0.85 in the Daylight fingerprints). Therefore, behavioral approaches can successfully complement the structural paradigm. To this end, similar interaction profiles are valuable resources in drug repurposing, as drugs with similar target binding patterns may exhibit a similar pharmacologic activity^63, 65, 66^.

From our repurposing hints list, we select Azelaic acid (saturated dicarboxylic acid) and Meprobamate (carbamate derivative) as possible antineoplastic and antifungal, respectively. However, the two hints are not structurally similar to the respective reference drugs (i.e., Progesterone and Abiraterone for antineoplastic, and Clotrimazole, Oxiconazole, Naftifine, Tolnaftate, Nystatin, Natamycin, Ciclopirox, Griseofulvin for antifungal). Indeed, Progesterone and Abiraterone are steroid derivatives, Clotrimazole and Oxiconazole are imidazole derivatives, Ergosterol has a steroidal structure, Terbinafine and Naftifine are allylamine compounds, Griseofulvin is a 3-coumaranone derivative. Nonetheless, as the chemical similarity is not necessarily a reliable predictor of biological similarity^59, 64^, we analyze the binding modes of Azelaic acid and Meprobamate in comparison with the other known reference drugs.

Although the results of the molecular docking analysis are not fully reliable predictors, we highlight the docking simulation results for the interaction between Azelaic acid and Steroid 17-alpha-hydroxylase/17,20 lyase, indicating a high similarity with the interaction of Progesterone and Abiraterone with the same target. Indeed, the lowest free energy of binding is −8.49 kcal/mol for Azelaic acid, −8.72 kcal/mol for Progesterone, −8.99 kcal/mol for Abiraterone, and the inhibition constant at K is 600.71 nM for Azelaic acid, 406.22 nM for Progesterone, 402.33 nM for Abiraterone (see supplementary material SupplementaryInformation.pdf, Table S4). DrugBank lists Progesterone as a substrate and inhibitor, and Abiraterone as an inhibitor of Steroid 17-alpha-hydroxylase/17,20 lyase. All the lowest free energy values suggest very similar stability of the three complexes. The results also indicate high similarity between the repositioning hint (i.e., Azelaic acid) and the reference anticancer drugs (i.e., Progesterone and Abiraterone) in terms of the inhibition constant. Azelaic acid and Progesterone similarly bind to the target, as both drugs interact with the same 8 amino acids in the target, and 5 out of the 8 interactions are of the same type. Azelaic acid and Abiraterone interact with the same 5 amino acids in the target, and 2 interactions are of the same type. Moreover, these docking simulation results are in line with references^67, 68^, which report the covalent bonding of Abiraterone and Steroid 17-alpha-hydroxylase/17,20 lyase (a cysteinato-heme enzyme from the cytochrome P450 superfamily). Precisely, Abiraterone forms a coordinate covalent bond of the pyridine nitrogen at C17 with heme iron of this target^68^.

The overarching conclusion of our molecular docking results is that Azelaic acid represents a promising candidate for further in silico (e.g., molecular dynamics), in vitro, and in vivo investigations of its potential anticancer effects. Although the molecular docking results are not as strong as for Azelaic acid, the antifungal properties of Meprobamate cannot be disregarded or rejected. Meprobamate is a known oral drug; however, we cannot exclude the topical route of administration as an antifungal. To this end, we need further investigations on biopharmaceutical properties to test various pharmaceutical topical formulations with Meprobamate as an active ingredient. The same discussion on the biopharmaceutical properties is valid for Azelaic acid, knowing that its route of administration may change as an anticancer drug.

## Methods

### Databases

We build our Drug-Drug Similarity Network (DDSN) using drug-target interaction information from the older Drug Bank version 4.2^30^, such that we can use the latest Drug Bank 5.1.4^69^ for testing the accuracy of our drug property prediction.

### Network analysis

Our Drug-Drug Similarity Network (DDSN) is an undirected weighted graph *G* = (*V, E*), where *V* is the vertex (or node) set, and *E* is the edge (or link) set. As such, we have |*V*| vertices *v_i_*∈*V* and |*E*| edges *e_j,k_* ∈ *E*, with *i*, *j*, *k* ∈ {1, 2, …|*V*|} and *j*≠*k*. Each edge *e_j,k_* is characterized by a weight *w*(*e_j,k_*)≠0 (in our DDSN, *w*(*e_j,k_*)∈ℕ^*^, ∀*e_j,k_*∈*E*). In an unweighted network, *w*(*e_j,k_*)=1, ∀*e_j,k_*∈*E*.

### Network centralities

Node centralities are complex network parameters, which are characterizing the importance of a vertex/node in a graph^70^. In our analysis, we used the weighted degree, degree, betweenness, and betweenness/degree node centralities.

The weighted degree of a node *v_i_* is the sum of the weights characterizing the links/edges incident to *v_i_*,

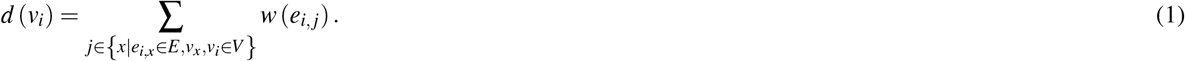

We compute the degree of a node *v_i_* with equation 1, assuming that *w*(*e_i,j_*) = 1, *e_ij_* ∈ *E*.

To compute the node betweenness, we must find the shortest paths between all node pairs (*v_j_*, *v_k_*) in graph *G*, namely *σ_j,k_*. As such, the betweenness of node *v_i_* is the number of minimal paths in graph *G* that cross node *v_i_*, divided by the total number of minimal paths in *G*,

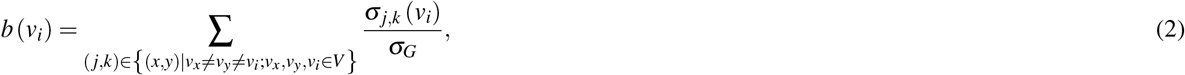

where the total number of shortest paths in *G* is 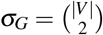.

The betweenness/degree of node *v_i_* is the ratio

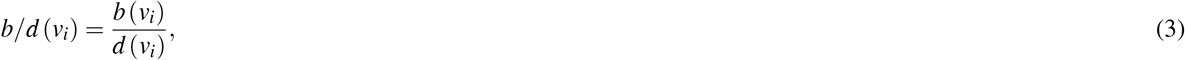

where equation 1 computes *d*(*v_i_*) in the unweighted version (i.e., considering *w*(*e_i,j_*) = 1, ∀*e_ij_* ∈ *E*).

### Community detection

The network layout algorithm we use in this paper places each vertex *v_i_* in a 2D space ℝ×ℝ = ℝ^2^. Therefore, each node *v_i_* ∈ *V* has its 2D coordinates *γ_i_* = (*x_i_, y_i_*) ∈ ℝ^2^, and each edge *e_i,j_* ∈ *E* has a Euclidian distance *δ_i,j_* = |*γ_i_* − *γ_j_*|.

In an energy-model, force-directed layout, we have a force of attraction between any two adjacent nodes *v_i_* and *v_j_*, and a repulsion force between any two non-adjacent nodes. The expression of these forces is 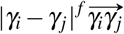 where *f* = *a* for attraction and *f* = *r* for repulsion. The attraction force between adjacent nodes (*v_i_* and *v_j_* such that ∃*e_i,j_* ∈ *E*) decreases, whereas the repulsion force between non-adjacent nodes (*v_i_, v_j_* such that ∃!*e_i,j_* ∈ *E*) increases with the Euclidian distance. Therefore, we must have *a* ≥ 0 and *r* ≤ 0.

In this paper, we use the energy-model force-directed layout Force Atlas 2 to assign node positions in the 2D (i.e., ℝ^2^) space, based on interactions between attraction and repulsion forces, such that we attain minimal energy in the network layout,

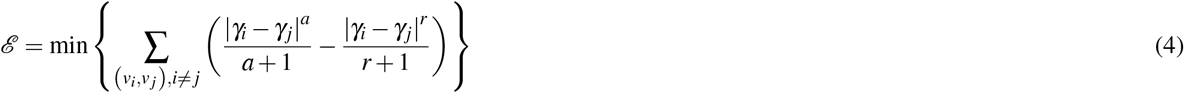

The energy-based layouts generate topological communities because there are specific regions in the network with larger than average link densities. Noack^34^ demonstrated that the energy-based topological communities are equivalent to the network clusters based on modularity classes^32^, when *a* > −1 and *r* > −1.

The network clustering classifies each node *v_i_* ∈ *V* in one of the disjoint sets of nodes (cluster *C_i_* ⊂ *V*, with 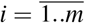, *C*_1_⋃*C*_2_…⋃*C_m_* = *V*). In^32^, the authors use modularity to define the node membership to one of the clusters. To this end, the modularity of a clustering 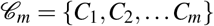 is

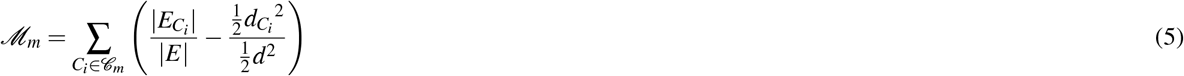

where |*E*| is the total number of edges in *G*, 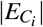 is the total number of edges between nodes in cluster *C_i_*, *d* is the total degree of nodes in *G*, and 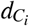 is the total degree of nodesm in cluster *C_i_*. Thus, 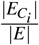 represents the relative edge density of cluster *C_i_* relative to the density of the entire network *G*, where as 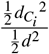 is the expected relative density of *C_i_*.

### Molecular docking for repurposing testing

#### Testing procedure

To verify the predicted properties of any repurposing hint, we perform molecular docking not only for the hint but also for the reference drugs (typical drugs having the predicted property) and for some drugs with little probability of having the predicted property. To this end, we formalize the following testing procedure.

1. We define the drug sets to enter the docking process, consisting of drugs hinted as having the pharmacological property *ϕ* 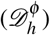, well-documented drugs with property *ϕ* (reference drugs 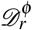), and drugs with little probability of having propert 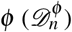. Our goal is to explore the similarity (in terms of relevant target activity) between the reference drugs 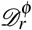 and the tested drugs 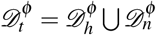.
  a. 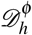 consists of the drugs hinted as repurposed for property/properties *ϕ*.
  b. 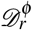 consists of two subsets, reference drugs in the DDSN’s community 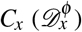 and reference drugs not in 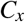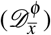, with 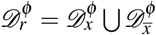.
  c. 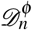 contains typical drugs for other pharmacological properties, with little probability of having property *ϕ*.
2. We establish the target sets. Specifically, for pharmacological property *ϕ*, we take into consideration the targets from DrugBank that interact with the drugs in the hinted drug 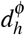 community *C_x_* having property 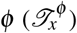, and the targets from DrugBank that interact with the drugs with property *ϕ* not included in DDSN’s 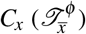.
3. For the set of tested drugs 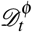, we use molecular docking to check the interactions between all possible drug-target pairs, defined as the Cartesian product of sets 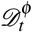 and 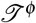 (with 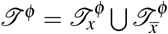),

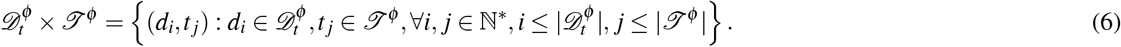
4. For the set of reference drugs, we apply molecular docking on separately designed drug-target pairs for reference drugs in 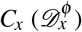, and reference drugs not in 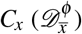 respectively, such that any drug-target pair is well-documented in the literature,

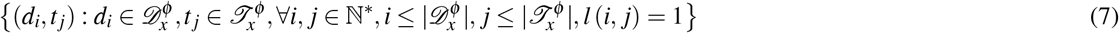

and

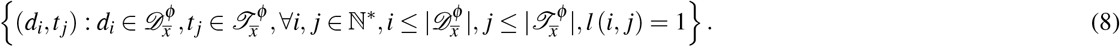

In equations 7 and 8, boolean function *l* is defined as

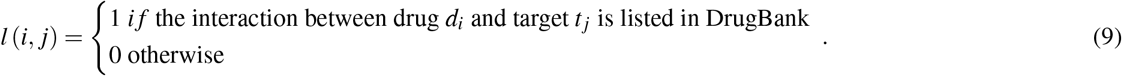

#### Ligands preparation

We generate the three-dimensional coordinates of all ligands using the Gaussian program suite with the DFT/B3LYP/6-311G optimization procedure.

#### Targets preparation

We get the X-ray crystal structure of the targets as target.pdb files from the major protein databases Protein Data Bank ^1^ and optimize them with the ModRefiner software ^2^. The targets and their corresponding codes are Lanosterol 14-alpha demethylase (4LXJ code, resolution 1.9 A), Intermediate conductance calcium-activated potassium channel protein 4 (6D42 code, resolution 1.75 A), Lanosterol synthase (1W6K code, resolution 2.1 A), Squalene monooxygenase (6C6N code, resolution 2.3 A), Ergosterol (2AIB code, resolution 1.1 A), Sodium/potassium-transporting ATPase subunit alpha (2ZXE code, resolution 2.4 A), and Tubulin (4U3J code, resolution 2.81). The preparation of targets also requires adding all polar hydrogens, removing the water, and computing the Gasteiger charge.

#### Docking protocol

We perform the molecular docking analysis using the Autodock 4.2.6 software suite together with the molecular viewer and graphical support AutoDockTools.

In the docking protocol, for the protein targets, we create the grid box using Autogrid 4 with 120 Å × 120 Å × 120 Å in *x*, *y* and *z* directions, and 1 Å spacing from the target molecule’s center. For steroidal target Ergosterol, the grid box is 30 Å × 30 Å × 30 Å in *x*, *y* and *z* directions, with 0.375 Å spacing from the target molecule’s center.

For the docking process, we chose the Lamarckian genetic algorithm (Genetic Algorithm combined with a local search), with a population size of 150, a maximum of 2.5·10^6^ energy evaluations, a gene mutation rate of 0.02, and 50 runs. We adopted the default settings for the other docking parameters and performed all the calculations in vacuum conditions. Then, we exported all AutoDock results in the PyMOL ^3^, and the Discovery Studio (Biovia) molecular visualization system ^4^.

## Supporting information

Supplementary information related to the manuscript.

Drug repurposing confirmation database.

## Author contributions statement

L.U., M.U., I.O.S., and P.B. conceived the drug repurposing procedure, A.T. and M.U. performed the network analysis, L.U., A.C., and I.O.S. performed the expert pharmacological and biochemical interpretation of the results, L.U. and R.-M.V. performed the molecular docking analysis, M.U., L.U., P.B., and I.O.S. wrote the paper. All authors reviewed the manuscript.

## Additional information

## Competing financial interests

The authors declare no competing financial interests.

## Supporting information

We present the validation references, and the statistics that support the drug cluster/community labeling in *Supplementary-DDSN.xlsx*. Also, all additional details indicated mentioned in the main manuscript are given in *SupplementaryInformation.pdf*. The Gephi file with our DDSN, entitled *DDSN.gephi* is available for download at the following URL: https://sites.google.com/site/analizamedicamentuluiumft/datasets. Visualizing Gephi files requires Gephi https://gephi.org/.

## Footnotes

1 http://www.rcsb.org/pdb/home/home.do

2 https://zhanglab.ccmb.med.umich.edu/ModRefiner/

3 The PyMOL Molecular Graphics System, Version 2.0 Schrödinger, LLC

4 Dassault Systémes BIOVIA, BIOVIA Workbook, Release 2017; BIOVIA Pipeline Pilot, Release 2017, San Diego: Dassault Systémes, 2019

## Notes

### Competing Interest Statement

The authors have declared no competing interest.

