## Supplementary information related to the manuscript. for "Uncovering new drug properties in target-based drug-drug similarity networks"

Lucreția Udrescu[1], Paul Bogdan[2], Aimée Chiș[3], Ioan Ovidiu Sîrbu[3], Alexandru Topîrceanu[4], Renata-Maria Văruț[5], and Mihai Udrescu[4]*

[1]"Victor Babeș" University of Medicine and Pharmacy Timișoara, Department of Drug Analysis, Timișoara 300041, Romania
[2]University of Southern California, Ming Hsieh Department of Electrical Engineering, Los Angeles, CA 90089-2563, USA
[3]"Victor Babeș" University of Medicine and Pharmacy Timișoara, Department of Biochemistry, Timișoara 300041, Romania
[4]University Politehnica of Timișoara, Department of Computer and Information Technology, Timișoara 300223, Romania
[5]University of Medicine and Pharmacy of Craiova, Faculty of Pharmacy, Craiova 200349, Romania

*

### 1. Introduction

Supplementary Information contains tables and figures with the molecular docking results for drug-target interaction simulations. The tables provide the details of the molecular drug-target interactions, in terms of the lowest free energy of binding, the estimated inhibition constant, and the type of bonds/interactions between the target amino acids and the molecular fragments of the drug. The figures illustrate the 2D and 3D maps of interactions between the tested drugs and their targets, highlighting the amino acid residues in targets that establish specific interactions with the drug molecule.

The first six tables (Tables S1-S6) and six figures (Figures S1-S6) illustrate the molecular docking results for testing the interactions between six hormone-related cancer targets and three categories of drugs: the repositioning hint (*i.e.*, **Azelaic acid**), the anticancer reference drugs (*i.e.*, Progesterone and Abiraterone), and reference drugs with no reported interaction with the target (*i.e.*, Fosinopril and Furosemide).

The next seven tables (Tables S7-S13) and figures (Figures S7-S13) show the molecular docking results for testing the interactions between seven fungal-related targets and four categories of drugs: the repositioning hint (*i.e.*, **Meprobamate**), the reference drugs with a known antifungal activity which belong to the topological community 25 (*i.e.*, Clotrimazole, Oxiconazole), the reference drugs with a known antifungal activity which do not belong to the topological community 25 (*i.e.*, Naftifine, Tolnaftate, Nystatin, Natamycine, Ciclopirox, and Griseofulvin), and reference drugs with no reported interaction with the target (*i.e.*, Fosinopril and Furosemide).

### 2. Molecular docking results for Azelaic acid

Table S1. The molecular interactions between Azelaic acid (i.e., the repositioning hint – $d_h^{anticancer}$ ), Progesterone and Abiraterone drugs from $\mathscr{D}_6^{anticancer}$ reference drugs with already accounted anticancer activity), Fosinopril and Furosemide (drugs from $\mathscr{D}_n^{anticancer}$ reference drugs, with no reported anticancer activity) with Estrogen receptor alpha. The residues shown in bold represent the common Estrogen receptor alpha amino acids involved in the same type of interaction with the tested drugs (Estrogen receptor alpha is a target from $\mathscr{T}_6^{anticancer}$ ).

| $\mathscr{T}_6^{anticancer}$ **Estrogen receptor alpha** | | | | | | | | | |
|---|---|---|---|---|---|---|---|---|---|
| Drug name | Drug role | Lowest free energy of binding [kcal/mol] | Estimated inhibition constant [Temp 298.15 K] | Conventional hydrogen bond | Carbon hydrogen bond | Alkyl interaction | Pi-alkyl interaction | Pi-sigma interaction | Interactive amino acid residues (Van der Waals interaction) |
| **Azelaic acid** | Repositioning hint (No reported interaction with the target) | -4.48 | 524.50 µM | ALA$_{A307}$<br>ARG$_{A363}$ (2)<br>ASP$_{A369}$ | - | - | - | - | ALA$_{A318}$, ASP$_{A321}$, ALA$_{A322}$, VAL$_{A364}$, PRO$_{A365}$, **GLY$_{A366}$**, VAL$_{A368}$ |
| Progesterone | Reference drug (agonist, inhibitor, downregulator) | -5.89 | 48.06 µM | VAL$_{A364}$<br>VAL$_{A367}$<br>VAL$_{A368}$ | - | ALA$_{A318}$, (2)<br>ARG$_{A363}$ (2)<br>PRO$_{A365}$ | - | - | ALA$_{A307}$, LEU$_{A310}$, GLN$_{A314}$, LYS$_{A362}$, **GLY$_{A366}$**, ASP$_{A369}$ |
| Abiraterone | Reference drug (No reported interaction with the target) | -7.11 | 6.09 µM | LYS$_{A362}$<br>VAL$_{A368}$ | | ALA$_{A307}$<br>LEU$_{A310}$<br>ALA$_{A318}$<br>PRO$_{A365}$ | | | GLN$_{A314}$, ARG$_{A363}$, **GLY$_{A366}$**, PHE$_{A367}$ |
| Fosinopril | Reference drug (No reported interaction with the target) | -2.03 | 32.30 mM | ILE$_{A326}$ | PRO$_{A325}$ | ILE$_{A326}$ (2)<br>TRP$_{A393}$ (2)<br>ARG$_{394}$ | ILE$_{A326}$ (2)<br>TRP$_{A393}$ (2)<br>ARG$_{394}$ | - | LEU$_{A320}$, GLU$_{A323}$, PRO$_{A324}$, GLY$_{A422}$, PHE$_{A44}$5, VAL$_{A446}$ |
| Furosemide | Reference drug (No reported interaction with the target) | -3.62 | 2.20 mM | ARG$_{A434}$<br>LEUV$_{509}$ | - | ALA$_{A430}$<br>ARG$_{A434}$<br>HIS$_{A513}$ | ALA$_{A430}$<br>ARG$_{A434}$<br>HIS$_{A513}$ | HIS$_{A513}$ | THR$_{431}$, GLN$_{506}$, ILE$_{A510}$,SER$_{A512}$ |

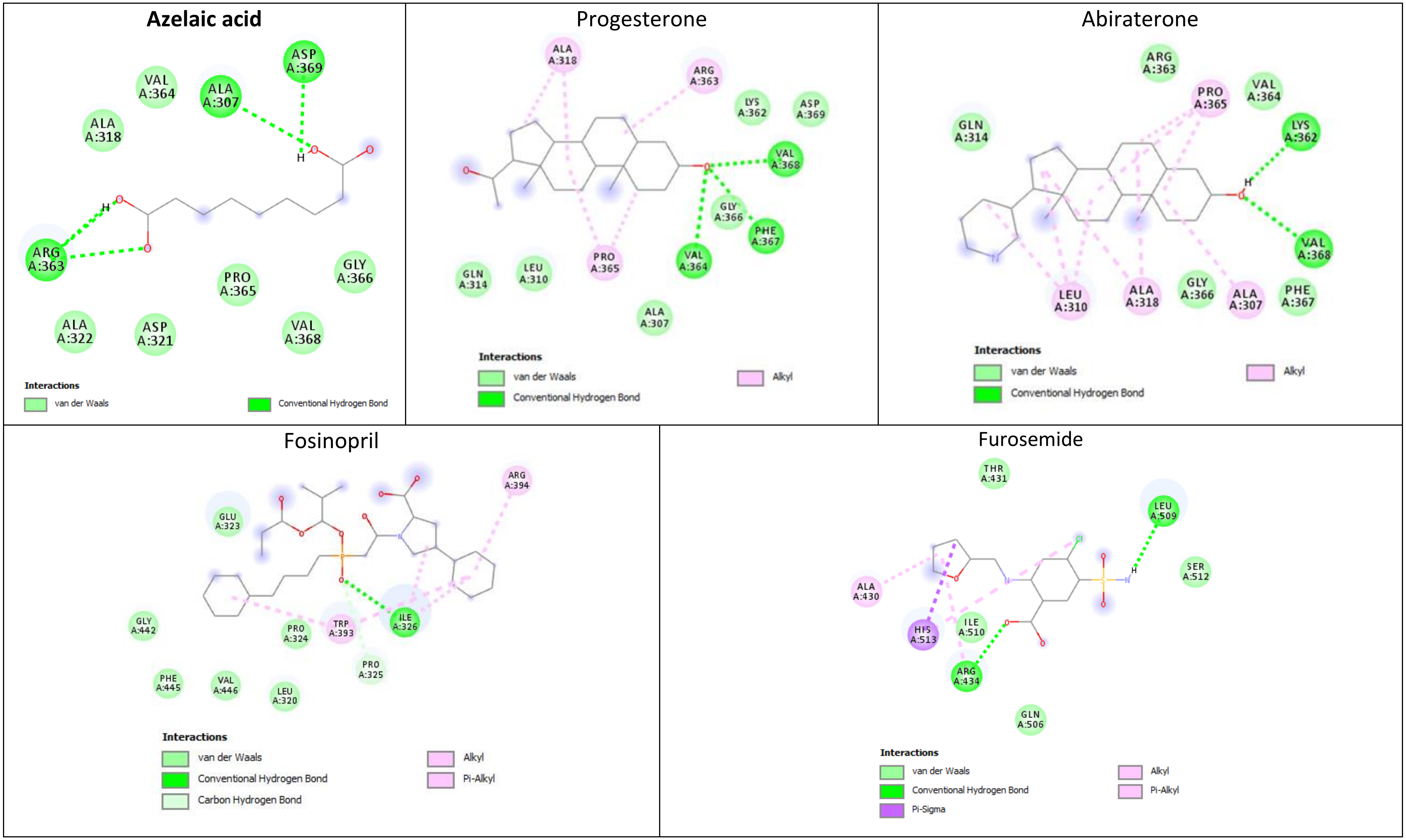


Figure S1. 2D molecular model of interactions between Azelaic acid, Progesterone, Abiraterone, Fosinopril and Furosemide (drugs from $\mathscr{D}_t^{anticancer}$ ) and the amino acid residues in Estrogen receptor alpha (a target from $\mathscr{T}_6^{anticancer}$ ). The docking software places the 2D chemical representation of the drug molecule in the center of each square. The colored disks represent the amino acids surrounding the drug molecule, while the dotted lines represent the interactions between the target's amino acids and the drug molecule. which is surrounded by the interacting amino acids (represented as colored disks) of the target (shown as dotted-lines); the maps indicates the target's amino acids that establish van der Waals interactions with the drug, but these interactions are not represented as dotted-lines. The diagrams also indicate the amino acids that establish van der Waals interactions with the drug, but these interactions are not represented.

Table S2. The molecular interactions between Azelaic acid (i.e., the repositioning hint – $d_h^{\text{anticancer}}$ ), Progesterone and Abiraterone (drugs from $\mathscr{D}_6^{\text{anticancer}}$ reference drugs, with already accounted anticancer activity), Fosinopril and Furosemide (drugs from $\mathscr{D}_n^{\text{anticancer}}$ reference drugs, with no reported anticancer activity) with Estrogen receptor beta (a target from $\mathscr{T}_6^{\text{anticancer}}$ ). The residues shown in bold represent the common Estrogen receptor beta amino acids involved in the same type of interaction with the tested drugs.

| $\mathscr{T}_6^{\text{anticancer}}$ **Estrogen receptor beta** | | | | | | | | | |
|---|---|---|---|---|---|---|---|---|---|
| Drug name | Drug role | Lowest free energy of binding [kcal/mol] | Estimated inhibition constant [Temp 298.15 K] | Conventional hydrogen bond | Carbon hydrogen bond | Alkyl interaction | Pi-Alkyl interaction | Halogen bond | Interactive amino acid residues (Van der Waals interaction) |
| **Azelaic acid** | Repositioning hint (No reported interaction with the target) | -3.11 | 5.29 µM | $GLU_{A305}$<br>$\mathbf{ARG_{A346}}$<br>$\mathbf{GLY_{A472}}$ | - | - | - | | $\mathbf{MET_{A295}}$, $LEU_{A298}$, $LEU_{A301}$, $\boldsymbol{MET_{A336}}$, $LEU_{A339}$, $MET_{A340}$, $LEU_{A343}$, $PHE_{A356}$, $\mathbf{ILE_{A373}}$, $\mathbf{MET_{A473}}$, $HIS_{A475}$, $LEU_{A476}$ |
| Progesterone | Reference drug (agonist, downregulator) | -8.68 | 435.53nM | $\mathbf{ARG_{A346}}$<br>$PHE_{A356}$<br>$\mathbf{GLY_{A472}}$<br>$HIS_{A475}$<br>$LEU_{A476}$ | | $LEU_{A298}$<br>$LEU_{A301}$<br>$ALA_{A302}$<br>$MET_{A336}$<br>$LEU_{A339}$<br>$MET_{A340}$<br>$LEU_{A343}$<br>$ILE_{A376}$ | $PHE_{A356}$ (2) | | $\mathbf{MET_{A295}}$, $GLU_{A305}$, $\mathbf{ILE_{A373}}$, $LEU_{A380}$, $\mathbf{MET_{A473}}$, $MET_{A479}$ |
| Abiraterone | Reference drug (No reported interaction with the target) | -7.9 | 1.62 µM | $SER_{A333}$<br>$GLU_{A337}$ | | $MET_{A473}$ | $TRP_{A335}$,<br>$TYR_{A488}$ | | $GLU_{A332}$, $CYS_{A334}$, $\boldsymbol{MET_{A336}}$, $ARG_{A466}$, $ASN_{A470}$, $LYS_{A471}$, $HIS_{B467}$ |
| Fosinopril | Reference drug (No reported interaction with the target) | -2.54 | 13.68 mM | $LEU_{B263}$<br>$HIS_{B428}$ | $SER_{B264}$<br>$PRO_{B265}$ | $VAL_{B438}$<br>$MET_{A453}$ | - | | $MET_{B261}$, $ASP_{B431}$, $ALA_{B432}$, $ASP_{B435}$, $TRP_{B439}$, $GLN_{A449}$, $GLN_{A450}$ |
| Furosemide | Reference drug (No reported interaction with the target) | -3.51 | 2.66 mM | $PRO_{A285}$<br>$ALA_{A287}$<br>$PHE_{A289}$ | | | $PHE_{A289}$ | $GLU_{A366}$ | $SER_{A283}$, $ARG_{A284}$, $PRO_{A288}$ |

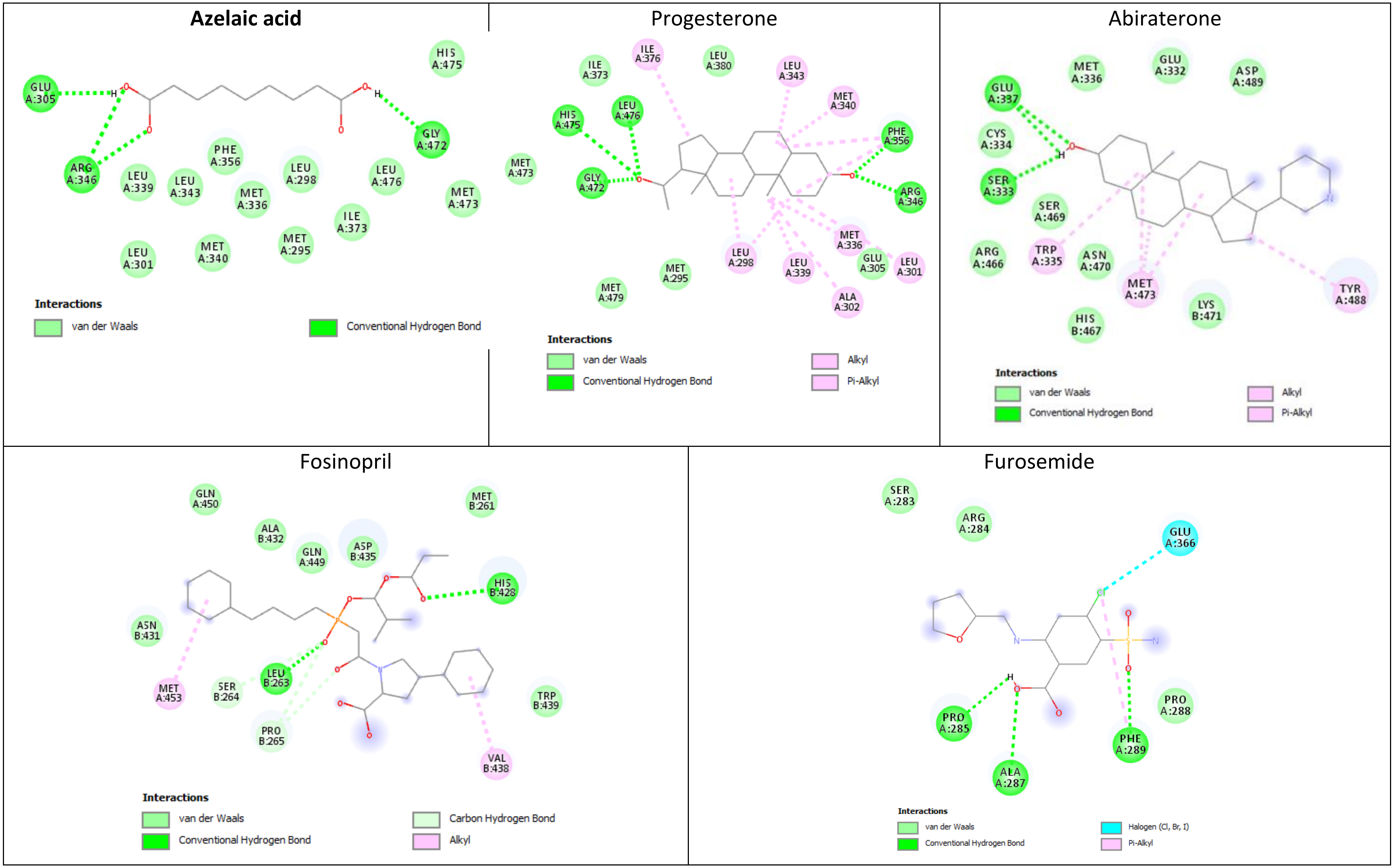


Figure S2. The 2D molecular model generated by the molecular docking simulation, for interactions between Azelaic acid, Progesterone, Abiraterone, Fosinopril, and Furosemide ($d_t^{\text{anticancer}}$ drugs), and the amino acid residues in Estrogen receptor beta (a target from $\mathscr{T}_6^{\text{anticancer}}$). The docking software places the 2D chemical representation of the drug molecule in the center of each square. The colored disks represent the amino acids surrounding the drug molecule, while the dotted lines represent the interactions between the target's amino acids and the drug molecule. The diagrams also indicate the amino acids that establish van der Waals interactions with the drug, but these interactions are not represented.

Table S3. The molecular interactions between Azelaic acid (i.e., the repositioning hint – $d_h^{\text{anticancer}}$ ), Progesterone and Abiraterone (a drug from $\mathscr{D}_6^{\text{anticancer}}$ reference drug with already accounted anticancer activity), Fosinopril and Furosemide (drugs from $\mathscr{D}_n^{\text{anticancer}}$ reference drugs, with no reported anticancer activity) with Progesterone receptor. The residues shown in bold represent the common Progesterone receptor (a target from $\mathscr{T}_6^{\text{anticancer}}$ ) amino acids involved in the same type of interaction with the tested drugs.

| $\mathscr{T}_6^{\text{anticancer}}$ **Progesterone receptor** | | | | | | | | |
|---|---|---|---|---|---|---|---|---|
| Drug name | Drug role | Lowest free energy of binding [kcal/mol] | Estimated inhibition constant [Temp 298.15 K] | Conventional hydrogen bond | Carbon hydrogen bond | Alkyl interaction | Pi-Alkyl interaction | Interactive amino acid residues (Van der Waals interaction) |
| **Azelaic acid** | Repositioning hint (No reported interaction with the target) | -4.54 | 471.16 µM | **GLN$_{B725}$**<br>**ARG$_{B766}$**<br>LEU$_{B887}$ | - | - | - | ***LEU$_{B721}$***, MET$_{B756}$, MET$_{B759}$, **VAL$_{B760}$, LEU$_{B763}$, PHE$_{B778}$**, LEU$_{B797}$, **MET$_{B801}$**, HIS$_{B888}$, TYR$_{B890}$, **CYS$_{B891}$** |
| Progesterone | Reference drug (agonist) | -11.17 | 6.47 nM | ASP$_{B719}$<br>**GLN$_{B725}$**<br>**ARG$_{B766}$** | | LEU$_{B718}$<br>LEU$_{B721}$<br>MET$_{B756}$<br>MET$_{B759}$<br>LEU$_{B797}$ | TYR$_{B890}$ | LEU$_{B715}$, **VAL$_{B760}$**, MET$_{B722}$, TRP$_{B755}$, **LEU$_{B763}$, PHE$_{B778}$, MET$_{B801}$, CYS$_{B891}$**, THR$_{B894}$, VAL$_{B903}$, PHE$_{B905}$, MET$_{B909}$ |
| Abiraterone | Reference drug (No reported interaction with the target) | -11.90 | 1.89 nM | MET$_{B759}$,<br>***ARG$_{B766}$*** | | LEU$_{B715}$<br>LEU$_{B718}$<br>LEU$_{B797}$<br>LEU$_{B887}$<br>CYS$_{B891}$ | PHE$_{B778}$ | ASN$_{B719}$, ***LEU$_{B721}$***, GLY$_{B722}$, TRP$_{B755}$, THR$_{B894}$, VAL$_{B903}$, PHE$_{B905}$, MET$_{B909}$ |
| Fosinopril | Reference drug (No reported interaction with the target) | -2.97 | 6.69 mM | ASP$_{B697}$,<br>LYS$_{B731}$ | SER$_{B728}$ | PRO$_{B696}$<br>TRP$_{B732}$ | | SER$_{B693}$, ILE$_{B694}$, GLU$_{B695}$, ILE$_{B699}$, ARG$_{B724}$, GLN$_{B725}$, LEU$_{B727}$, SER$_{B728}$, SER$_{B735}$ |
| Furosemide | Reference drug (No reported interaction with the target) | -4.88 | 265.75 µM | GLN$_{A682}$ (2)<br>LEU$_{A683}$<br>ILE$_{A684}$ | | | | ASN$_{A689}$, MET$_{A692}$ |

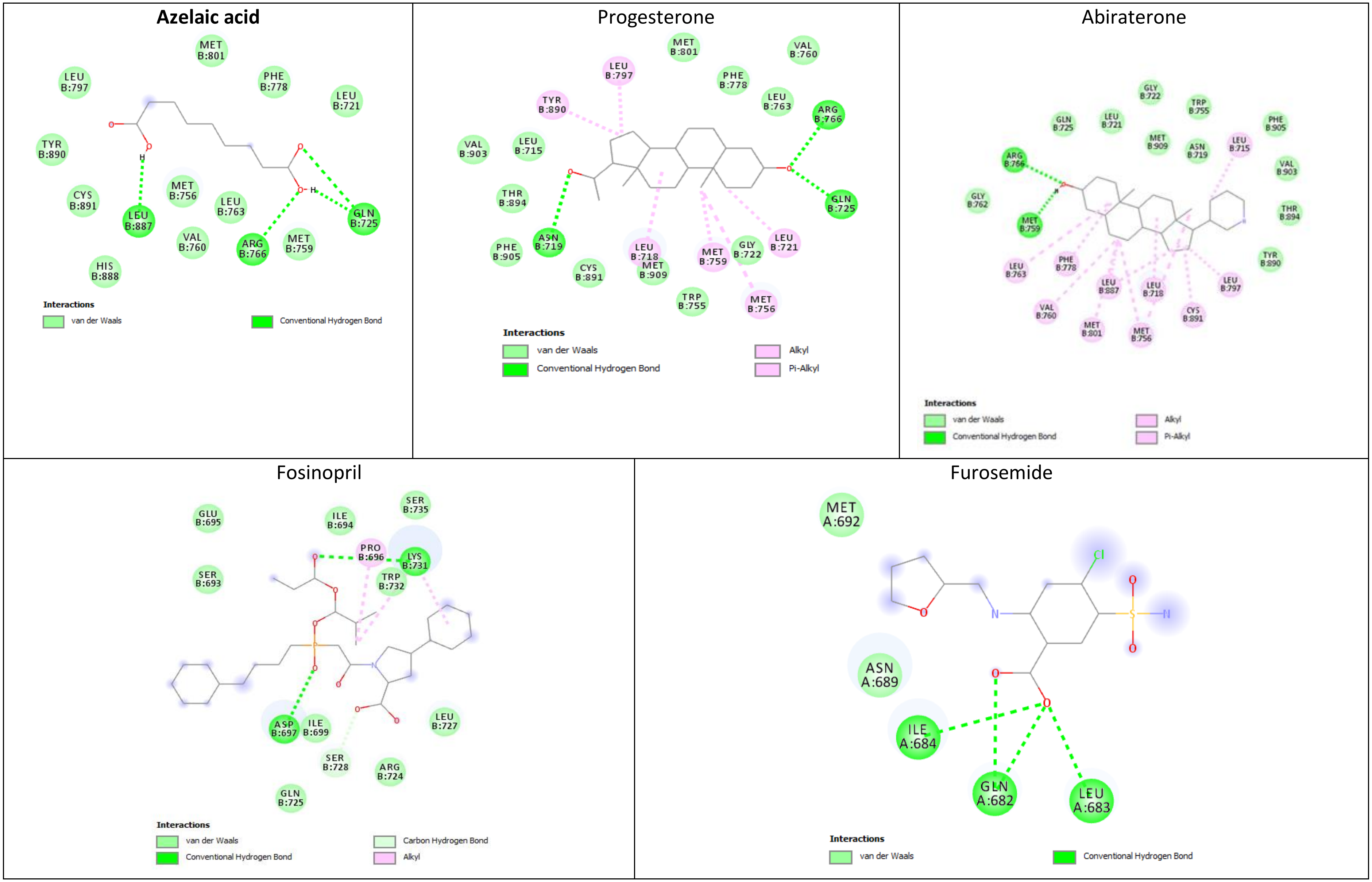


Figure S3. The 2D diagrams generated by the molecular docking simulation, for interactions between Azelaic acid, Progesterone, Abiraterone, Fosinopril, and Furosemide (i.e., $d_t^{\text{anticancer}}$ drugs), and the amino acid residues in Progesterone receptor (a target from $\mathscr{T}_6^{\text{anticance}}$). The docking software places the 2D chemical representation of the drug molecule in the center of each square. The colored disks represent the amino acids surrounding the drug molecule, while the dotted lines represent the interactions between the target's amino acids and the drug molecule. The diagrams also indicate the amino acids that establish van der Waals interactions with the drug, but these interactions are not represented.

Table S4. The molecular interactions between Azelaic acid (i.e., the repositioning hint – $d_h^{\text{anticancer}}$), Progesterone and Abiraterone ($d_6^{\text{anticancer}}$ drugs from $\mathscr{D}_6^{\text{anticancer}}$ reference drugs with known anticancer activity), Fosinopril and Furosemide (drugs from $\mathscr{D}_n^{\text{anticancer}}$ reference drugs, with no reported anticancer activity) with Steroid 17-alpha-hydroxylase/17,20 lyase. The residues shown in bold represent the common Steroid 17-alpha-hydroxylase/17,20 lyase amino acids involved in the same type of interaction with the test and reference drugs (Steroid 17-alpha-hydroxylase/17,20 lyase is a target from $\mathscr{T}_6^{\text{anticancer}}$).

| $\mathscr{T}_6^{\text{anticancer}}$ **Steroid 17-alpha-hydroxylase/17,20 lyase** | | | | | | | | | |
|---|---|---|---|---|---|---|---|---|---|
| Drug name | Drug role | Lowest free energy of binding [kcal/mol] | Estimated inhibition constant [Temp 298.15 K] | Covalent bond | Conventional hydrogen bond | Carbon hydrogen bond | Alkyl interaction | Pi-Alkyl interaction | Interactive amino acid residues (Van der Waals interaction) |
| **Azelaic acid** | Reference drug (No reported interaction with the target) | **-8.49** | **600.71 nM** | | $ARG_{B96}$, $ILE_{B112}$, $TRP_{B121}$,$ARG_{B12}$, **$ILE_{B371}$**, **$HIS_{B373}$**, $ARG_{B440}$ | - | - | - | ***$ALA_{B113}$***, **$LEU_{B370}$**, **$SER_{B441}$**, $CYS_{B442}$, **$GLY_{B436}$** |
| Progesterone | Reference drug (substrate, inhibitor) | -8.72 | 406.22 nM | | **$ILE_{B371}$, $HIS_{B373}$** | | $VAL_{B366}$ (3) $ALA_{B367}$ $CYS_{B442}$ (2) HEM | $PHE_{B435}$ | $LEU_{B86}$, $ARG_{B96}$, $THR_{B306}$, $LEU_{B361}$, **$LEU_{B370}$**, $PRO_{B434}$, **$GLY_{B436}$**, $ALA_{B437}$, $ARG_{B440}$, **$SER_{B441}$**, $ALA_{B448}$ |
| Abiraterone | Reference drug (inhibitor) | -8.99 | 402.33nM | HEM | $ASN_{202}$ | $ALA_{B113}$, $GLY_{B436}$ | $ILE_{B205}$ $ILE_{B206}$ $LEU_{B209}$ $ALA_{B367}$ $LEU_{B370}$ $VAL_{B482}$ | $PHE_{B114}$ | ***$ALA_{B113}$***, $ARG_{B239}$, $GLY_{B301}$, $PRO_{B434}$, **$GLY_{B436}$** |
| Fosinopril | Reference drug (No reported interaction with the target) | -3.86 | 1.47 mM | | $ASN_{A51}$, $ARG_{A364}$ | - | $HIS_{A48}$, $LEU_{A476}$ | $HIS_{A48}$ | $GLY_{A47}$, $HIS_{A50}$, $LYS_{A55}$, $ASP_{C241}$, $LYS_{C245}$, $PHE_{A317}$, $LEU_{A363}$, $TRP_{A397}$, $HIS_{A401}$, $ASP_{A410}$, $GLN_{A411}$, PHEA412, $GLU_{A477}$, $PHE_{A484}$ |
| Furosemide | Reference drug (No reported interaction with the target) | -5.64 | 74.01 µM | | $LYS_{C55}$ $LYS_{C59}$ | $LEU_{C56}$ | $PHE_{C42}$ $ARG_{C45}$ | $PHE_{C42}$ $ARG_{C45}$ | $GLY_{C47}$, $ASN_{C52}$, $LEU_{C56}$, $LYS_{C58}$ |

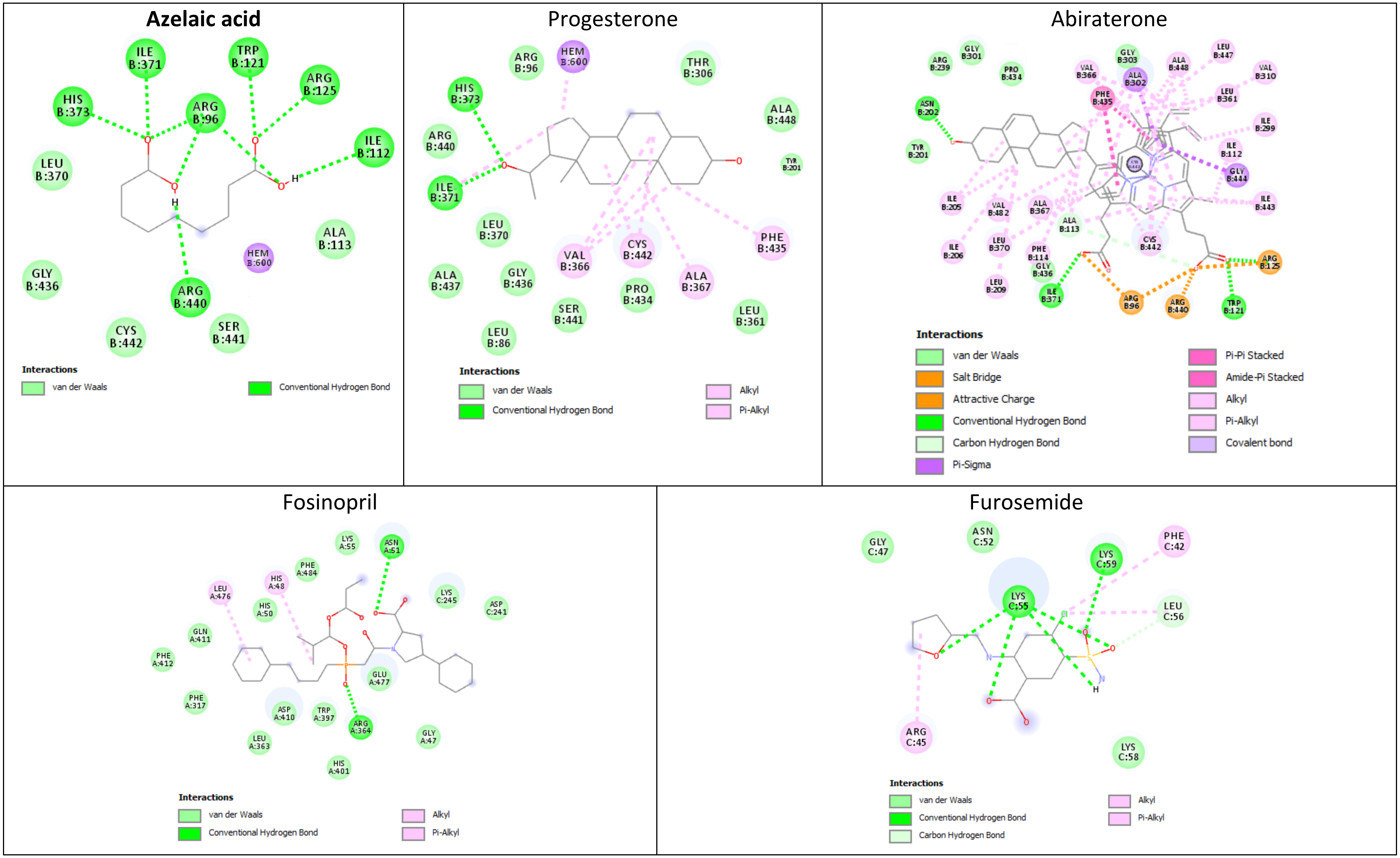


Figure S4. Structure views of the 3D-complexes between Azelaic acid (i.e., the repositioning hint – $d_h^{anticancer}$ ), Progesterone and Abiraterone ( $d_6^{anticancer}$ drugs from $\mathscr{D}_6^{anticancer}$ reference drug with known anticancer activity), Fosinopril and Furosemide (reference drugs from $\mathscr{D}_n^{anticancer}$ , with no reported anticancer activity) with Steroid 17-alpha-hydroxylase/17,20 lyase, which is a target from $\mathscr{T}_6^{anticancer}$ . The docking software places the 2D chemical representation of the drug molecule in the center of each square. The colored disks represent the amino acids surrounding the drug molecule, while the dotted lines represent the interactions between the target's amino acids and the drug molecule. The diagrams also indicate the amino acids that establish van der Waals interactions with the drug, but these interactions are not represented.

Table S5. The molecular interactions between Azelaic acid (i.e., the repositioning hint – $d_h^{\text{anticancer}}$ ), Progesterone and Abiraterone ( $d_6^{\text{anticancer}}$ drugs from $\mathscr{D}_6^{\text{anticancer}}$ reference drugs with known anticancer activity), Fosinopril and Furosemide (drugs from $\mathscr{D}_n^{\text{anticancer}}$ reference drugs, with no reported anticancer activity) with Androgen receptor (a target from $\mathscr{T}_6^{\text{anticancer}}$ ). The residues shown in bold represent the common target's amino acids involved in the same type of interaction with the test and reference drugs.

| $\mathscr{T}_6^{\text{anticancer}}$ **Androgen receptor** | | | | | | | | |
|---|---|---|---|---|---|---|---|---|
| Drug name | Drug role | Lowest free energy of binding [kcal/mol] | Estimated inhibition constant [Temp 298.15 K] | Conventional hydrogen bond | Pi-donor hydrogen bond | Alkyl interaction | Pi-Alkyl interaction | Interactive amino acid residues (Van der Waals interaction) |
| **Azelaic acid** | Repositioning hint (No reported interaction with the target) | -5.01 | 213.54 µM | $LEU_{A704}$ **$MET_{A745}$** **$ARG_{A752}$(2)** $PHE_{A764}$ | $LEU_{A707}$ | - | - | $LEU_{A707}$, **$GLY_{A708}$, $GLN_{A711}$**, $MET_{A742}$, **$VAL_{A746}$**, $ALA_{A748}$, **$MET_{A749}$**, $LEU_{A873}$ |
| Progesterone | Reference drug (agonist, potentiator) | -10.28 | 29.31 nM | **$MET_{A745}$** **$ARG_{A752}$** | - | $LEU_{A704}$ (2), $LEU_{A707}$, $MET_{A742}$, $MET_{A745}$, $MET_{A780}$, $LEU_{A873}$, $THR_{877}$ | - | $LEU_{A701}$, $ASN_{A705}$, **$GLY_{A708}$, $GLN_{A711}$**, $TRP_{A741}$, **$VAL_{A746}$, $MET_{A749}$**, $PHE_{A764}$, $PHE_{A876}$, $THR_{A877}$, $LEU_{A880}$, $PHE_{A891}$ |
| Abiraterone | Reference drug (No reported interaction with the target) | -8.56 | 527.57 nM | $THR_{A755}$ $LYS_{A808}$ | | $PRO_{A682}$, $VAL_{A684}$, $VAL_{A685}$, $ALA_{A748}$, $ARG_{A752}$ | | $GLY_{A683}$, **$GLN_{A711}$**, $HIS_{A714}$, $VAL_{A715}$, $LEU_{A744}$, $TRP_{A751}$, $PHE_{A804}$ |
| Fosinopril | Reference drug (No reported interaction with the target) | --2.91 | 7.41 mM | - | | $LYS_{A912}$, $PRO_{A913}$ | | $PRO_{A868}$, $ARG_{A871}$, $ILE_{A906}$, $GLY_{A909}$, $VAL_{A911}$, $ILE_{A914}$, $HIS_{A917}$, $THR_{A918}$ |
| Furosemide | Reference drug (No reported interaction with the target) | -4.13 | 938.39 µM | $GLU_{A793}$ (2) $LYS_{A861}$ | | $LEU_{A862}$ | $TYR_{A915}$ | $TRP_{A796}$, $LEU_{A797}$, $ASP_{A864}$, $SER_{A865}$, $PRO_{A868}$, $HIS_{A917}$, $THR_{A918}$ |

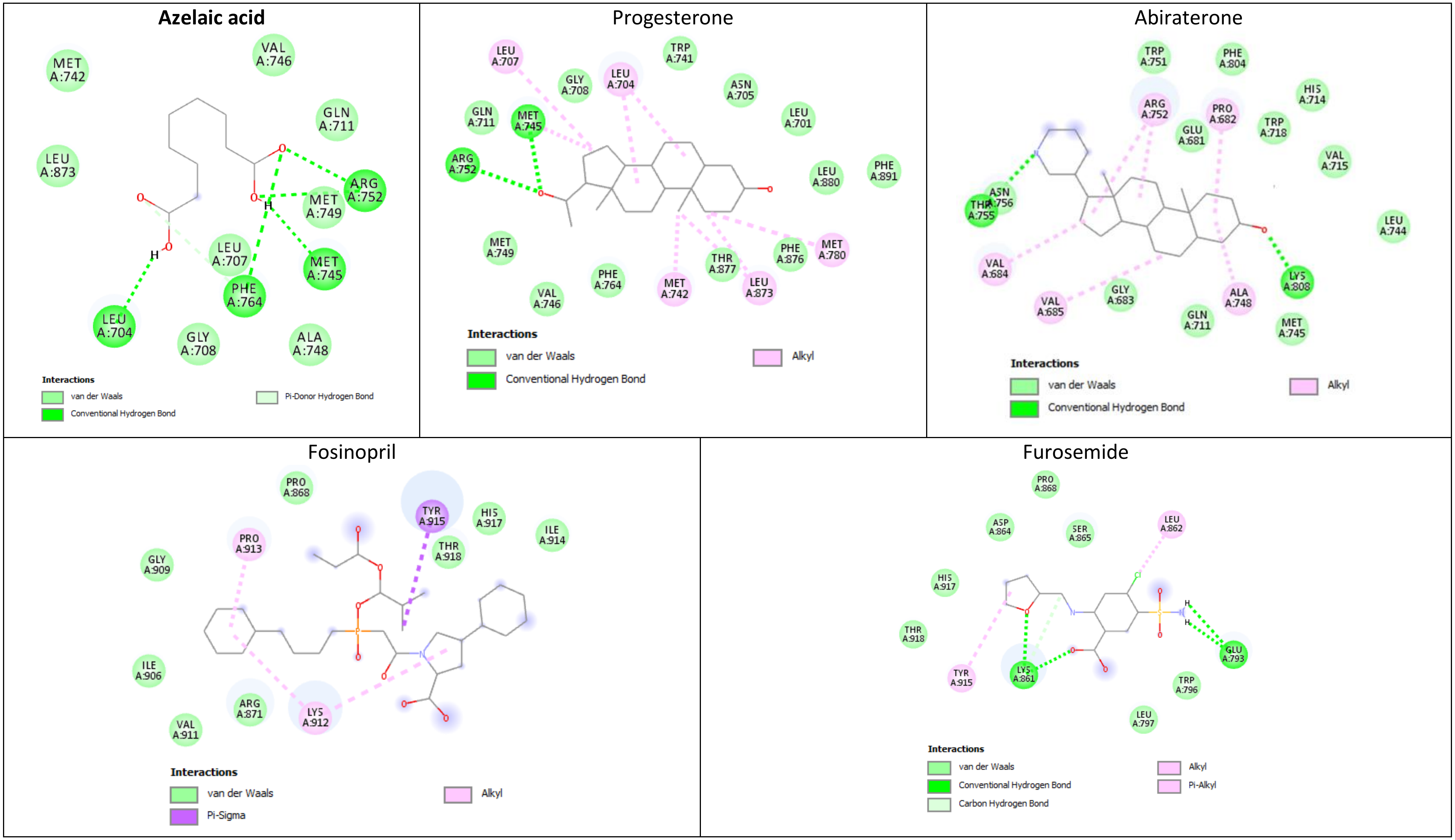


Figure S5. The 2D diagrams generated by the molecular docking simulation for interactions between Azelaic acid, Progesterone, Abiraterone, Fosinopril, and Furosemide (i.e., $d_t^{\text{anticancer}}$ drugs), and the amino acid residues in Androgen receptor (a target from $\mathscr{T}_6^{\text{anticancer}}$). The 2D chemical representation of the drug molecule is in the center of each square. The colored disks represent the amino acids surrounding the drug molecule, while the dotted lines represent the interactions between the target's amino acids and the drug molecule. The diagrams also indicate the amino acids that establish van der Waals interactions with the drug, but these interactions are not represented.

Table S6. The molecular interactions between Azelaic acid (i.e., the repositioning hint – $d_h^{\text{anticancer}}$ ), Progesterone and Abiraterone ( $d_6^{\text{anticancer}}$ drugs from $\mathscr{D}_6^{\text{anticancer}}$ reference drugs with known anticancer activity), Fosinopril and Furosemide (drugs from $\mathscr{D}_n^{\text{anticancer}}$ reference drugs, with no reported anticancer activity) with Mineralocorticoid receptor. The residues shown in bold represent the common Mineralocorticoid receptor amino acids involved in the same type of interaction with the test and reference drugs (Mineralocorticoid receptor is a target from $\mathscr{T}_6^{\text{anticancer}}$ ).

| $\mathscr{T}_6^{\text{anticancer}}$ **Mineralocorticoid receptor** | | | | | | | | |
|---|---|---|---|---|---|---|---|---|
| Drug name | Drug role | Lowest free energy of binding [kcal/mol] | Estimated inhibition constant [Temp 298.15 K] | Conventional hydrogen bond | Carbon hydrogen bond | Alkyl interaction | Pi-Alkyl interaction | Interactive amino acid residues (Van der Waals interaction) |
| **Azelaic acid** | Repositioning hint (No reported interaction with the target) | -4.49 | 507.15 µM | $ASN_{D770}$ $GLN_{D776}$ $\mathbf{ARG_{D817}}$ | | | | $LEU_{D769}$, $LEU_{D772}$, $ALA_{D773}$, $\mathbf{TRP_{D806}}$, $MET_{D807}$, $LEU_{D810}$, $ALA_{D813}$, $\mathbf{LEU_{D814}}$, $PHE_{D829}$, $CYS_{D942}$, $\mathbf{PHE_{D956}}$, $LEU_{D960}$ |
| Progesterone | Reference drug (antagonist, agonist) | -10.77 | 12.72 nM | $\mathbf{ARG_{D817}}$ | $CYS_{D942}$ | $LEU_{D769}$ $LEU_{D772}$ $ALA_{D773}$ $MET_{D807}$ $LEU_{D810}$ $LEU_{D938}$ $CYS_{D942}$ $MET_{D845}$ | $PHE_{D829}$, $PHE_{D941}$ | $ASN_{D770}$, $LEU_{D766}$, $GLN_{D776}$, $\mathbf{TRP_{D806}}$, $SER_{D811}$, $\mathbf{LEU_{D814}}$, $THR_{D945}$, $VAL_{D954}$, $\mathbf{PHE_{D956}}$ |
| Abiraterone | Reference drug (No reported interaction with the target) | -8.08 | 1.19 µM | $GLN_{F776}$ | | $PRO_{F747}$, $VAL_{F750}$, $ALA_{F813}$, $ARG_{F817}$, $LYS_{F820}$, $HIS_{F821}$ | $HIS_{F821}$ | $GLU_{F746}$, $GLU_{F748}$, $ILE_{F749}$, $GLN_{F779}$, $VAL_{F780}$, $TRP_{F816}$, $PHE_{F866}$ |
| Fosinopril | Reference drug (No reported interaction with the target) | -2.28 | 21.22 mM | $LYS_{F901}$ $LYS_{F905}$ | | $PRO_{F738}$ $TYR_{F899}$ | $PRO_{F738}$ $TYR_{F899}$ | $PRO_{F788}$, $ASN_{F898}$, $GLU_{F902}$, $ARG_{F904}$ |
| Furosemide | Reference drug (No reported interaction with the target) | -4.01 | 1.16 mM | $ASP_{D884}$ $LYS_{D977}$ | | $LYS_{D977}$ | | $LYS_{D883}$, $GLY_{D974}$, $PRO_{D978}$ |

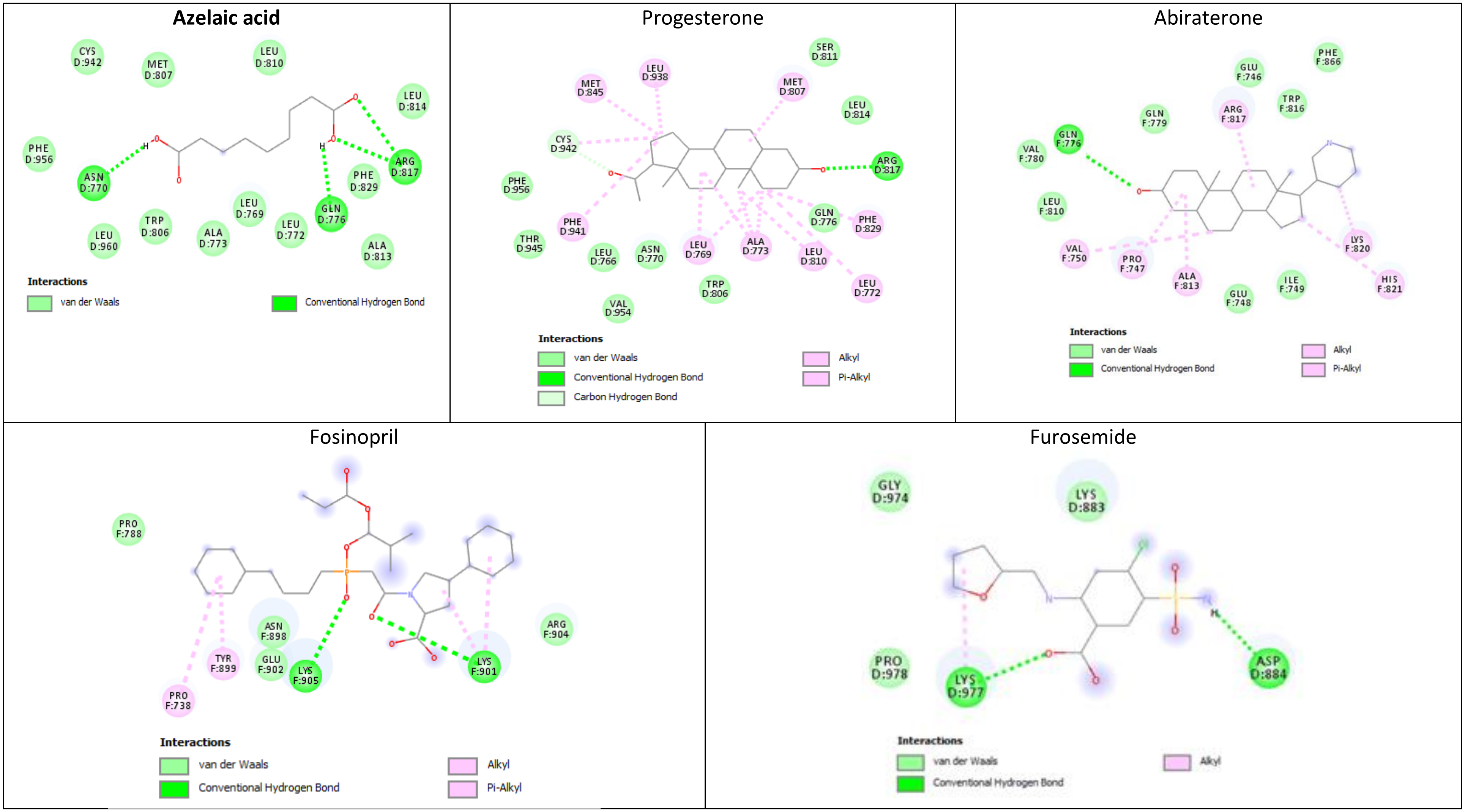


Figure S6. The 2D maps generated by the molecular docking simulation for interactions between Azelaic acid, Progesterone, Abiraterone, Fosinopril, and Furosemide (i.e., $d_t^{anticancer}$ drugs), and the amino acid residues in Mineralocorticoid receptor (a target from $\mathscr{T}_6^{anticancer}$). The 2D chemical representation of the drug molecule is in the center of each square. The colored disks represent the amino acids surrounding the drug molecule, while the dashed lines represent the interactions between the target's amino acids and the drug molecule. The diagrams also indicate the amino acids that establish van der Waals interactions with the drug, but these interactions are not represented.

### 3. Molecular docking results for Meprobamate

Table S7. A comparison of the molecular interactions between Meprobamate (i.e., the repositioning hint $-d_h^{\text{antifungal}}$), Clotrimazole (this drug $\in \mathscr{D}_{25}^{\text{antifungal}}$ reference drugs with documented antifungal activity), Fosinopril and Furosemide (reference drugs from $\mathscr{D}_n^{\text{antifungal}}$, with no reported antifungal activity) and Lanosterol 14-alpha demethylase (Lanosterol 14-alpha demethylase $\in \mathscr{T}_{25}^{\text{antifungal}}$). The residues shown in bold represent the Lanosterol 14-alpha demethylase amino acids involved in the same type of interaction with the tested and reference drugs.

| $\mathscr{T}_{25}^{\text{antifungal}}$ **Lanosterol 14-alpha demethylase** | | | | | | | | |
|---|---|---|---|---|---|---|---|---|
| Drug name | Drug role | Lowest free energy of binding [kcal/mol] | Estimated inhibition constant [Temp 298.15 K] | Conventional hydrogen bond | Carbon hydrogen bond | Alkyl interaction | Pi-alkyl interaction | Interactive amino acid residues (Van der Waals interaction) |
| **Meprobamate** | **Repositioning hint (No reported interaction with the target)** | -2.77 | 9.38 mM | **$MET_{A358}$**<br>$MET_{A360}$<br>$MET_{A460}$ | -- | **$PRO_{A210}$** | -- | $VAL_{A102}$, **$TYR_{A103}$**, **$PHE_{A105}$**, **$VAL_{A213}$**, **$LEU_{A356}$**, $LEU_{A357}$, $LEU_{A359}$, **$VAL_{A461}$** |
| Clotrimazole | Reference drug (Known antagonist, inhibitor) | -7.15 | 0.0058 mM | **$MET_{A358}$** | $MET_{A460}$ | **$PRO_{A210}$**<br>$LEU_{A357}$<br>$MET_{A358}$ | $PHE_{A48}$<br>$PHE_{A214}$ | $GLY_{A49}$, $ILE_{A72}$, **$TYR_{A103}$**, **$PHE_{A105}$**, **$VAL_{A213}$**, $PRO_{A355}$, **$LEU_{A356}$**, $MET_{A360}$, $TYR_{A457}$, $HIS_{A458}$, $THR_{A459}$, **$VAL_{A461}$**, $VAL_{A462}$ |
| Fosinopril | Reference drug (No reported interaction with the target) | -4.85 | 0.278 mM | -- | -- | $ALA_{A131}$<br>$LYS_{A426}$<br>$ILE_{A423}$ | -- | $ARG_{A124}$, $LEU_{A127}$, $ASN_{A128}$, $GLU_{A132}$, $LEU_{A134}$, $THR_{A135}$, $ILE_{A136}$, $PHE_{A139}$, $GLY_{A418}$, $VAL_{A419}$, $HIS_{A420}$, $LYS_{A421}$, $CYS_{A422}$, $GLY_{A424}$, $GLN_{A425}$, $PHE_{A427}$ |
| Furosemide | Reference drug (No reported interaction with the target) | -4.78 | 0.315 mM | $PRO_{A83}$<br>$HIS_{A84}$ (2)<br>$HIS_{A86}$<br>$SER_{A87}$<br>$GLU_{A409}$ (2) | $GLY_{A410}$ | -- | -- | $GLU_{A85}$, $ARG_{A88}$, $LEU_{A91}$, $VAL_{A408}$, $ALA_{A411}$ |

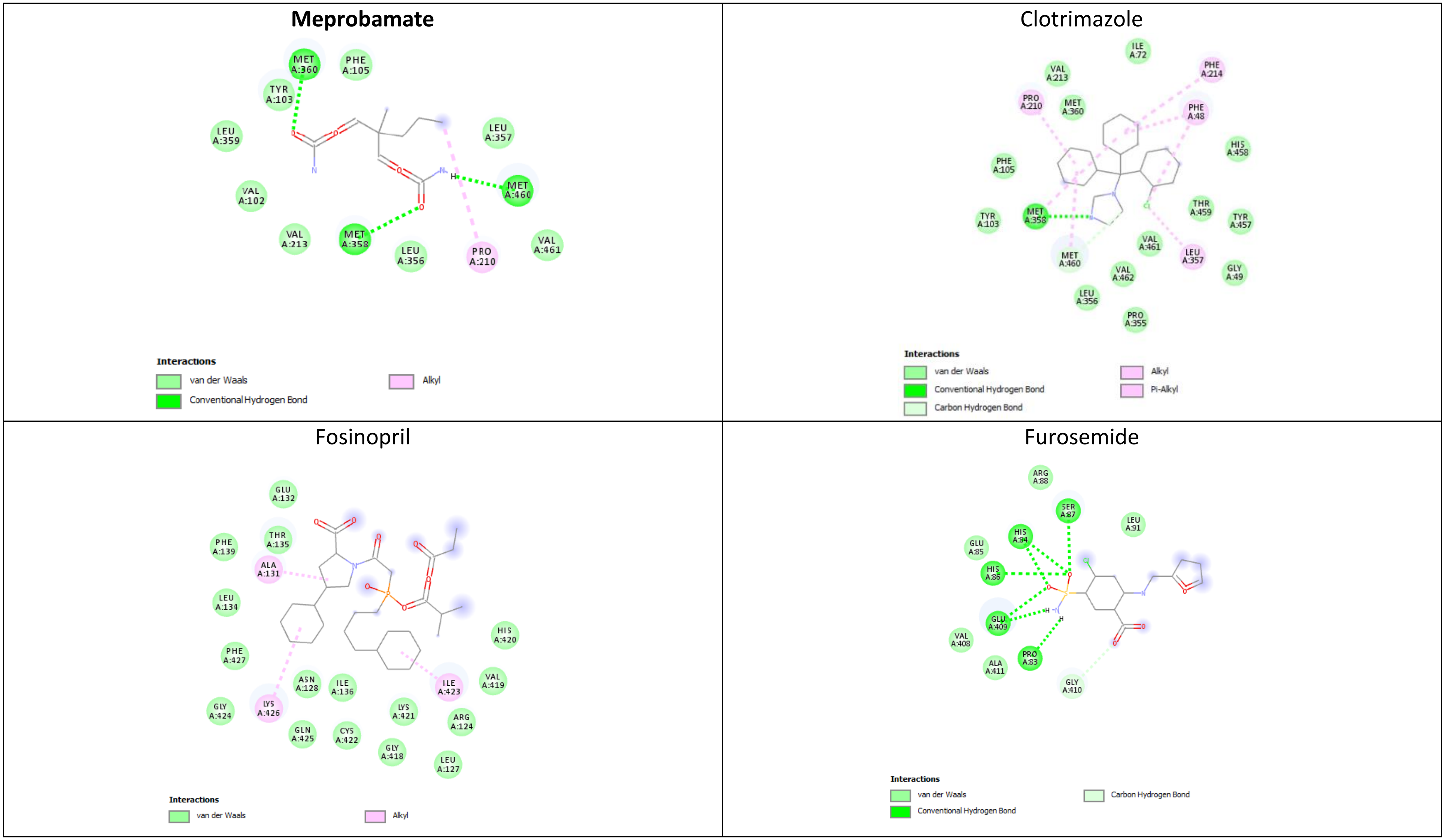


Figure S7. 2D molecular model of interactions between Meprobamate, Fosinopril, Furosemide (tested drugs that $\in \mathscr{D}_t^{\text{antifungal}}$), and Clotrimazole (a reference drug from $\mathscr{D}_{25}^{\text{antifungal}}$) with the amino acid residues in Lanosterol 14-alpha demethylase (a target $\in \mathscr{T}_{25}^{\text{antifungal}}$). The docking software places the 2D chemical representation of the drug molecule in the center of each square. The colored disks represent the amino acids surrounding the drug molecule, while the dotted lines represent the interactions between the target's amino acids and the drug molecule. The maps also indicate the target's amino acids that establish van der Waals interactions with the drug; however, for simplification, this figure does not represent these interactions.

Table S8. The comparison of the molecular interactions between Meprobamate (i.e., the repositioning hint – $d_h^{\text{antifungal}}$), Oxiconazole (this drug ∈ $\mathscr{D}_{25}^{\text{antifungal}}$ reference drugs, with already documented antifungal activity), Fosinopril and Furosemide (reference drugs from $\mathscr{D}_n^{\text{antifungal}}$, with no reported antifungal activity) and Lanosterol synthase (this target ∈ $\mathscr{T}_{25}^{\text{antifungal}}$). The residues shown in bold represent the Lanosterol synthase amino acids involved in the same type of interaction with the tested and reference drugs.

| $\mathscr{T}_{25}^{\text{antifungal}}$ **Lanosterol synthase** | | | | | | | | |
|---|---|---|---|---|---|---|---|---|
| Drug name | Drug role | Lowest free energy of binding [kcal/mol] | Estimated inhibition constant [Temp 298.15 K] | Conventional hydrogen bond | Carbon hydrogen bond | Alkyl interaction | Pi-Alkyl interaction | Interactive amino acid residues (Van der Waals interaction) |
| **Meprobamate** | Repositioning hint (No reported interaction with the target) | -3.23 | 4.26 mM | $THR_{A210}$<br>$LEU_{A211}$<br>$TRP_{A216}$<br>$SER_{A241}$ (2) | | $\mathbf{MET_{A215}}$<br>$\mathbf{ALA_{A224}}$<br>$\mathbf{LEU_{A515}}$ | | $\mathbf{PHE_{A212}}$, $LEU_{A229}$, $CYS_{A233}$, $\mathbf{TYR_{A237}}$, $TYR_{A297}$, $\mathbf{LEU_{A300}}$, $PRO_{A517}$, $\mathbf{MET_{A525}}$ |
| Oxiconazole | Reference drug (inhibitor) | -6.24 | 0.0267 mM | $MET_{A215}$ | | $\mathbf{MET_{A215}}$ (3)<br>$\mathbf{ALA_{A224}}$<br>$LEU_{A229}$<br>$LEU_{A292}$<br>$VAL_{A296}$<br>$\mathbf{LEU_{A515}}$ | $TRP_{A216}$ | $LEU_{A211}$, $\mathbf{PHE_{A212}}$, $PRO_{A213}$, $ALA_{A222}$, $PRO_{A223}$, $PRO_{A226}$, $\mathbf{TYR_{A237}}$, $SER_{A241}$, $LEU_{A299}$, $\mathbf{LEU_{A300}}$, $LEU_{A512}$, $\mathbf{MET_{A525}}$ |
| Fosinopril | Reference drug (No reported interaction with the target) | -5.04 | 0.2018 mM | $GLY_{A86}$ | $THR_{A49}$<br>$GLN_{A88}$ | $LEU_{A27}$<br>$CYS_{A29}$<br>$LEU_{A51}$<br>$ALA_{A89}$ | $PHE_{A64}$ | $ASN_{A28}$, $ARG_{A46}$, $GLY_{A50}$, $GLU_{A52}$, $TYR_{A63}$, $VAL_{A85}$, $LEU_{A87}$, $GLU_{A90}$, $ASP_{A91}$, $GLY_{A92}$, $THR_{A95}$, $GLU_{A406}$, $PHE_{A407}$, $SER_{A409}$, $CYS_{A410}$, $LYS_{A413}$ |
| Furosemide | Reference drug (No reported interaction with the target) | -5.61 | 0.0773 mM | $TYR_{A98}$<br>$GLY_{A380}$ | $TYR_{A704}$ | | $TRP_{A230}$<br>$HIS_{A232}$<br>$PHE_{A696}$ | $PRO_{A101}$, $PHE_{A103}$, $TRP_{A192}$, $GLY_{A336}$, $PRO_{A337}$, $ILE_{A338}$, $SER_{A339}$, $THR_{A381}$, $VAL_{A453}$, $THR_{A502}$, $TYR_{A503}$, $PHE_{A521}$, $TRP_{A581}$, $VAL_{A695}$, $ASN_{A697}$, $ILE_{A702}$ |

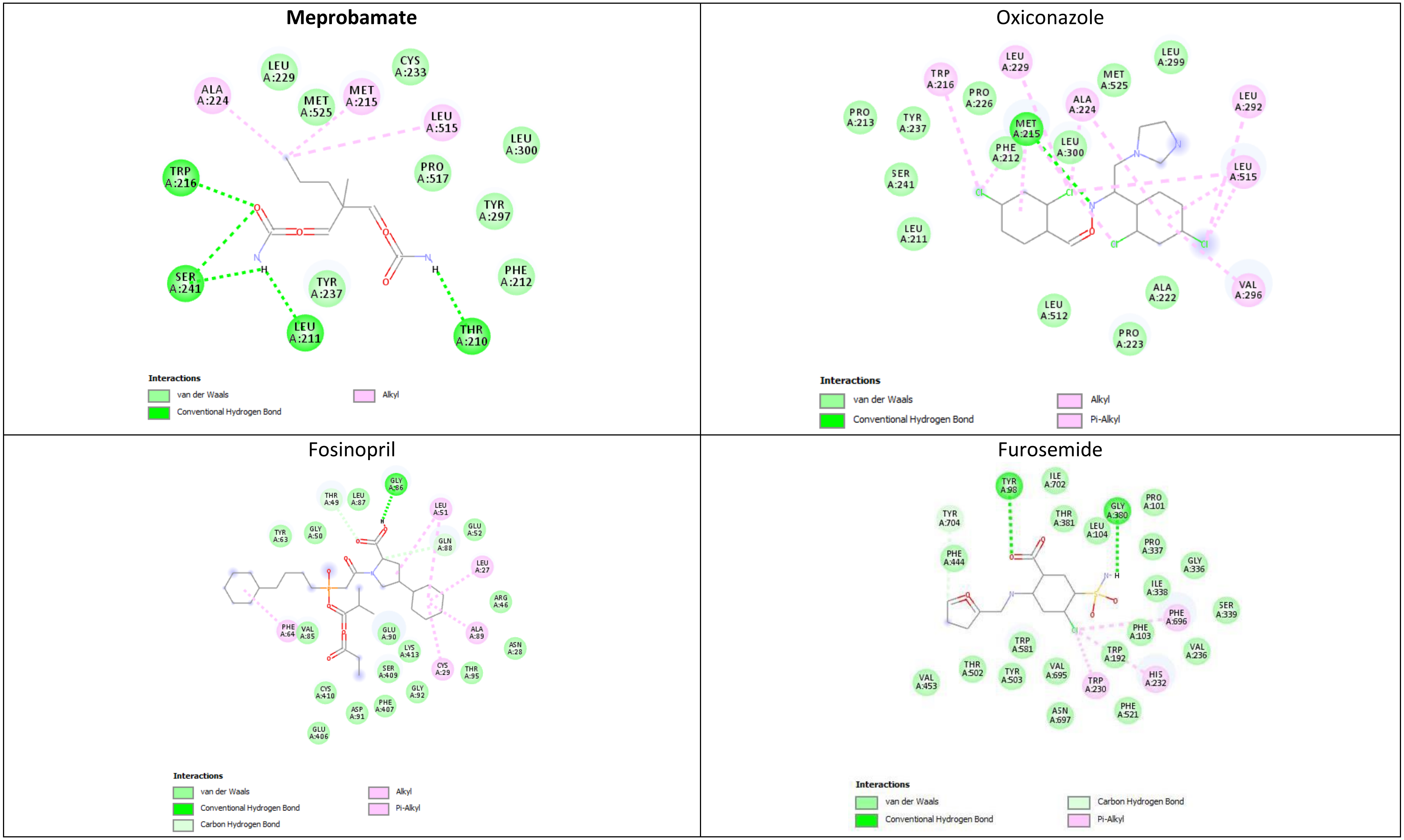


Figure S8. The 2D maps generated by the molecular docking simulation, for interactions between Meprobamate, Fosinopril, and Furosemide (tested drugs from $\mathscr{D}_t^{antifungal}$ drugs), Oxiconazole (the reference drug $\in \mathscr{D}_{25}^{antifungal}$ ) and the amino acid residues in Lanosterol synthase (the target $\in \mathscr{T}_{25}^{antifungal}$ ). The docking software places the 2D chemical representation of the drug molecule in the center of each square. The colored disks represent the amino acids surrounding the drug molecule, while the dotted lines represent the interactions between the target's amino acids and the drug molecule. The diagrams also indicate the amino acids that establish van der Waals interactions with the drug; however, for the sake of clarity, these interactions are not represented.

Table S9. A comparison of the molecular interactions between Meprobamate (i.e., the repositioning hint – $d_h^{\text{antifungal}}$ ), Clotrimazole (this drug ∈ $\mathscr{D}_{25}^{\text{antifungal}}$ reference drug with already documented antifungal activity), Fosinopril and Furosemide (these drugs ∈ $\mathscr{D}_{n}^{\text{antifungal}}$ reference drugs, with no reported antifungal activity) with Intermediate conductance calcium-activated potassium channel protein 4. The residues shown in bold represent the Intermediate conductance calcium-activated potassium channel protein 4 (this target ∈ $\mathscr{T}_{25}^{\text{antifungal}}$ ) amino acids involved in the same type of interaction with the reference and tested drugs.

| $\mathscr{T}_{25}^{\text{antifungal}}$ | **Intermediate conductance calcium-activated potassium channel protein 4** | | | | | | | | |
|---|---|---|---|---|---|---|---|---|---|
| Drug name | Drug role | Lowest free energy of binding [kcal/mol] | Estimated inhibition constant [Temp 298.15 K] | Conventional hydrogen bond | Carbon hydrogen bond | Alkyl interaction | Halogen interaction | Pi-Sulfur interaction | Interactive amino acid residues (Van der Waals interaction) |
| **Meprobamate** | **Repositioning hint (No reported interaction with the target)** | -1.02 | 179.63 mM | $ALA_{A405}$ (2) $GLU_{A408}$ $THR_{A412}$ $THR_{B398}$ | $THR_{A412}$ | $LEU_{A409}$ $ALA_{A413}$ | -- | -- | $GLN_{B395}$, $LYS_{B402}$ |
| Clotrimazole | Reference drug (Known antagonist, inhibitor) | -3.60 | 2.30 mM | -- | -- | $LYS_{A402}$ $ALA_{A405}$ | $GLU_{A408}$ | -- | $ASP_{A404}$, $THR_{A407}$ $ASP_{B397}$, $THR_{B398}$, $GLY_{B401}$, $ASP_{B404}$ |
| Fosinopril | Reference drug, randomly chosen (No reported interaction with the target) | -1.96 | 36.76 mM | -- | -- | $LEU_{B381}$ | -- | -- | $ILE_{B377}$, $LEU_{B378}$, $ASP_{B380}$, $ASN_{B384}$, $LEU_{B385}$ |
| Furosemide | Reference drug, randomly chosen (No reported interaction with the target) | -2.45 | 16.05 mM | $HIS_{A389}$ | $SER_{A386}$ | -- | -- | $HIS_{A389}$ | $LEU_{A378}$, $LEU_{A381}$, $GLN_{A382}$, $LEU_{A385}$, $TYR_{A379}$ |

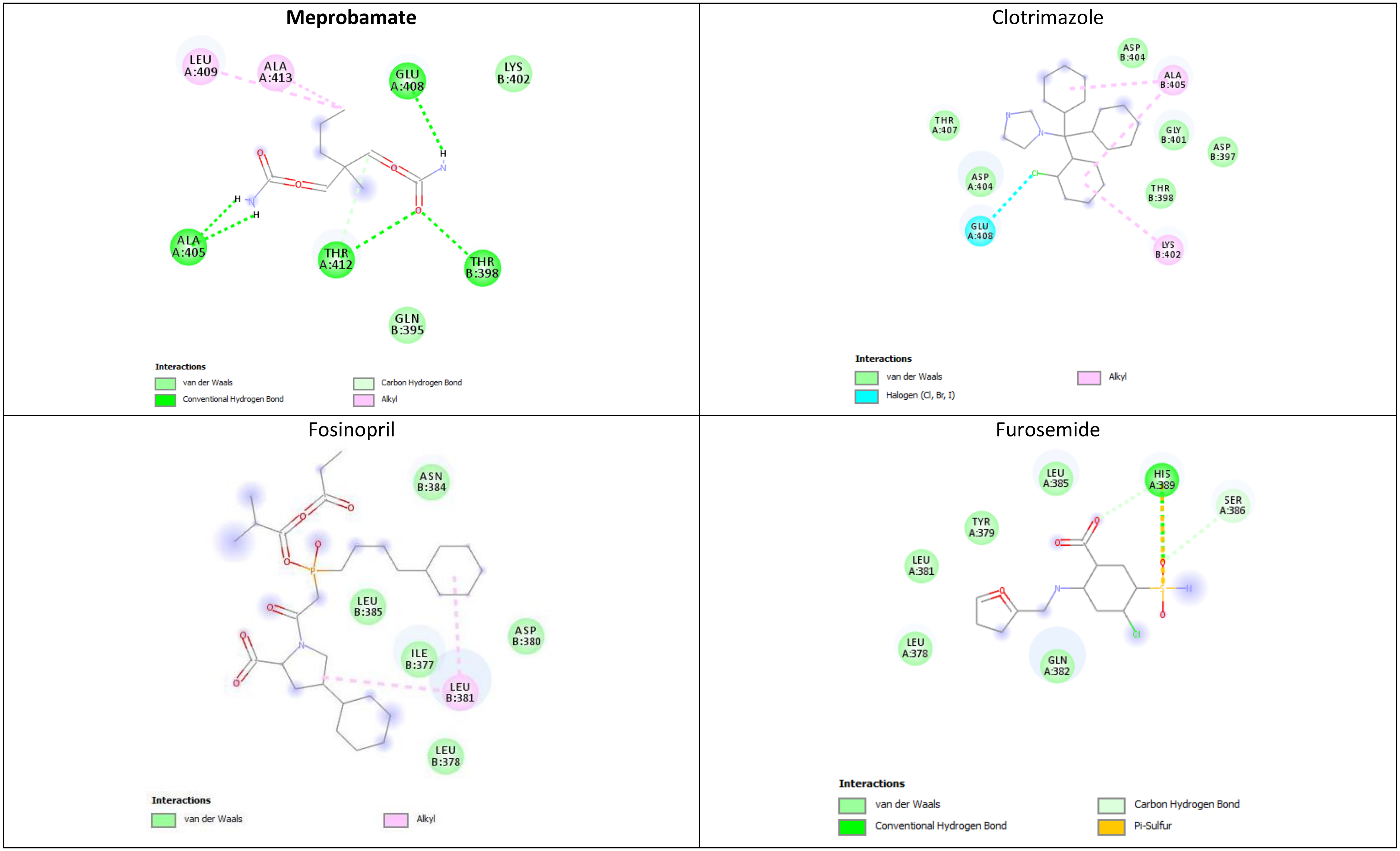


Figure S9. The 2D diagrams generated by the molecular docking simulation, for interactions between Meprobamate (the $d_h^{\text{antifungal}}$ repositioning hint), Fosinopril, and Furosemide (these reference drugs $\in \mathscr{D}_n^{\text{antifungal}}$), and Clotrimazole (the reference drug $\in \mathscr{D}_{25}^{\text{antifungal}}$) with the amino acid residues in Intermediate conductance calcium-activated potassium channel protein 4 (this target $\in \mathscr{T}_{25}^{\text{antifungal}}$). The docking software places the 2D chemical representation of the drug molecule in the center of each square. The colored disks represent the amino acids surrounding the drug molecule, while the dotted lines represent the interactions between the target's amino acids and the drug molecule. The diagrams also indicate the amino acids that establish van der Waals interactions with the drug; however, for the sake of clarity, these interactions are not represented.

Table S10. A comparison of the molecular interactions between Meprobamate (i.e., the repositioning hint – $d_h^{\text{antifungal}}$ ), Naftifine and Tolnaftate (these drugs $\in \mathscr{D}_{25}^{\text{antifungal}}$ reference drugs with already well-documented antifungal activity), Fosinopril and Furosemide (these drugs $\in \mathscr{D}_n^{\text{antifungal}}$ reference drugs, with no reported antifungal activity), and Squalene monooxygenase. The residues shown in bold represent the Squalene monooxygenase (this target $\in \mathscr{T}_{25}^{\text{antifungal}}$ ) amino acids involved in the same type of interaction with the tested and reference drugs.

| $\mathscr{T}_{25}^{\text{antifungal}}$ **Squalene monooxygenase** | | | | | | | | |
|---|---|---|---|---|---|---|---|---|
| Drug name | Drug role | Lowest free energy of binding [kcal/mol] | Estimated inhibition constant [Temp 298.15 K] | Conventional hydrogen bond | Carbon hydrogen bond | Alkyl interaction | Pi-Alkyl interaction | Interactive amino acid residues (Van der Waals interaction) |
| **Meprobamate** | Repositioning hint (No reported interaction with the target) | -2.67 | 11.01 mM | $GLY_{A164}$ $PHE_{A166}$ $GLY_{A418}$ $GLY_{A420}$ $MET_{A421}$ | | $\mathbf{VAL_{A163}}$ $LEU_{A287}$ | | $ILE_{A162}$, $\mathbf{GLU_{A165}}$, $LEU_{A167}$, $GLN_{A168}$, $PHE_{A306}$, $LEU_{A333}$, $\mathbf{TYR_{A335}}$, $LEU_{A345}$, $MET_{A388}$, $PRO_{A389}$, $\mathbf{ASP_{A408}}$, $\mathbf{ARG_{A413}}$, $PRO_{A415}$, $GLY_{A419}$, $THR_{A422}$ |
| Naftifine | Reference drug (inhibitor) | -7.47 | 0.0034 mM | | $PRO_{B415}$ | $VAL_{B163}$ $ALA_{B322}$ $LEU_{B333}$ $MET_{B421}$ | | $ILE_{B162}$, $GLY_{A164}$, $\mathbf{GLU_{A165}}$, $PHE_{B166}$, $GLN_{B168}$, $TYR_{B195}$, $PHE_{B306}$, $GLU_{B323}$, $LEU_{B324}$, $ILE_{B334}$, $\mathbf{TYR_{B335}}$, $LEU_{B345}$, $LEU_{B416}$, $THR_{B417}$, $GLY_{B418}$, $GLY_{B419}$, $GLY_{B420}$, $LEU_{B509}$ |
| Tolnaftate | Reference drug (inhibitor) | -6.75 | 0.0113 mM | $PRO_{A389}$ | $PRO_{A389}$ | $VAL_{A133}$ $ILE_{A162}$ $\mathbf{VAL_{A163}}$ $LEU_{A345}$ $MET_{A388}$ | $HIS_{A226}$ $PHE_{A306}$ | $ARG_{A161}$, $GLY_{A164}$, $\mathbf{GLU_{A165}}$, $ILE_{A230}$, $GLY_{A286}$, $LEU_{A287}$, $\mathbf{TYR_{A335}}$, $ALA_{A390}$, $SER_{A391}$, $\mathbf{ASP_{A408}}$, $MET_{A412}$, $\mathbf{ARG_{A413}}$, $HIS_{A414}$, $GLY_{A420}$, $MET_{A421}$ |
| Fosinopril | Reference drug (No reported interaction with the target) | -4.25 | 0.7694 mM | | | $LEU_{A547}$ $LEU_{A554}$ $LEU_{B554}$ $CYS_{A558}$ $CYS_{B558}$ | | $PRO_{A544}$, $ARG_{A545}$, $LEU_{A548}$, $TYR_{B566}$, $GLY_{A551}$, $GLY_{B551}$, $ALA_{A552}$, $SER_{B559}$, $TYR_{A555}$, $TYR_{B555}$, $PRO_{B563}$ |
| Furosemide | Reference drug (No reported interaction with the target) | -5.36 | 0.1186 mM | $HIS_{B198}$ $GLU_{B205}$ $ASP_{B370}$ | | $LYS_{B203}$ | | $ASP_{B199}$, $GLN_{B200}$, $GLU_{B201}$, $SER_{B204}$, $PHE_{B317}$, $PRO_{B366}$, $GLN_{B367}$, $ILE_{B368}$, $PRO_{B369}$, $HIS_{B371}$, $LYS_{B373}$ |

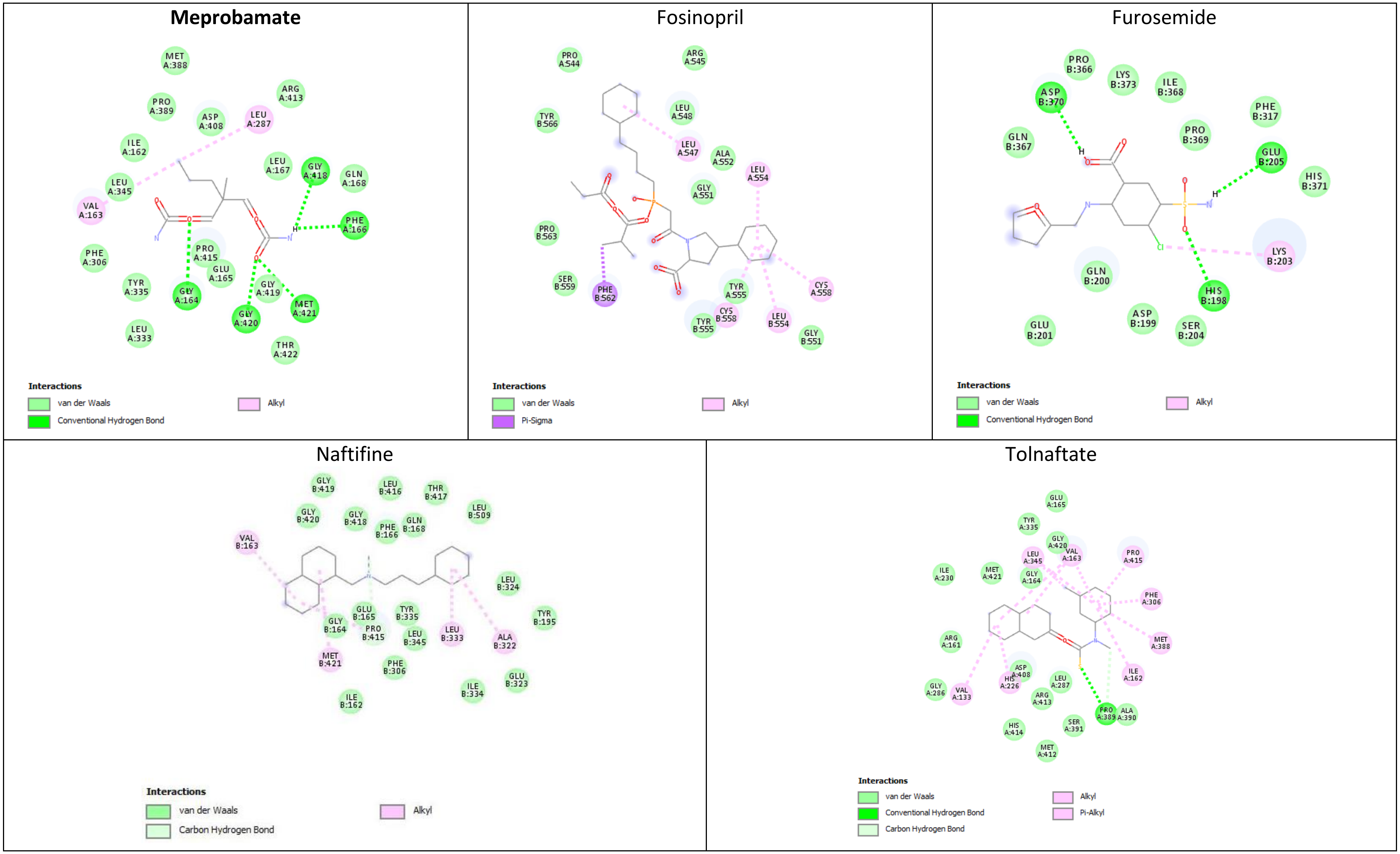


Figure S10. The 2D maps generated by the molecular docking simulation, for interactions of Meprobamate, Fosinopril, and Furosemide (i.e., $d_t^{\text{antifungal}}$ drugs), Naftifine and Tolnaftate (reference drugs from $\mathscr{D}_{25}^{\text{antifungal}}$ with already documented antifungal activity) with the amino acid residues in Squalene monooxygenase (this target $\in \mathscr{T}_{25}^{\text{antifungal}}$). The docking software places the 2D chemical representation of the drug molecule in the center of each square. The colored disks represent the amino acids surrounding the drug molecule, while the dotted lines represent the interactions between the target's amino acids and the drug molecule. The maps also indicate the amino acids that establish van der Waals interactions with the drug; however, for clarity, these interactions are not represented.

Table S11. A comparison of the molecular interactions between Meprobamate (i.e., the repositioning hint – $d_h^{antifungal}$ ), Clotrimazole (a $d_{25}^{antifungal}$ drug from $\mathscr{D}_{25}^{antifungal}$ reference drugs with documented antifungal activity), Nystatin and Natamycine (these drugs $\in \mathscr{D}_{\overline{25}}^{antifungal}$ reference drugs with well-documented antifungal activity), Fosinopril and Furosemide (these drugs $\in \mathscr{D}_n^{antifungal}$ reference drugs, with no reported antifungal activity) and Ergosterol (this target $\in \mathscr{T}_{\overline{25}}^{antifungal}$ ).

| $\mathscr{T}_{\overline{25}}^{antifungal}$ **Ergosterol** | | | | |
|---|---|---|---|---|
| | Drug role | Lowest free energy of binding [kcal/mol] | Estimated inhibition constant [Temp 298.15 K] | Description of drug-target molecular interaction |
| **Meprobamate** | Reference drug (No reported interaction with the target) | **-3.48** | **2.79 mM** | 4 hydrophobic alkyl/alkyl interaction that involves the pentyl radical (meprobamat) and the six-atom cycles (ergosterol) |
| Clotrimazole | Reference drug (inhibitor) | -4.06 | 1.05 mM | 8 hydrophobic alkyl/alkyl interaction that involves the 3 benzene rings (clotrimazole) and the six-atom cycles (ergosterol) |
| Nystatin | Repositioning hint (No reported interaction with the target) | -3.98 | 1.21 mM | 1 hydrogen bond between –1-OH of nystatin (oxanic cycle) and –OH (ergosterol) |
| Natamycin | Reference drug (inhibitor) | -6.24 | 0.0265 mM | 2 hydrogen bond between -3,4—dihydroxy of natamycine (methyloxan cylce) and –OH (ergosterol) |
| Fosinopril | Reference drug (No reported interaction with the target) | -3.79 | 1.67 mM | 10 hydrophobic alkyl/alkyl interaction that involves all three cycles (fosinopril) and the five-atom cycles and 18-$CH_3$ (ergosterol) |
| Furosemide | Reference drug (No reported interaction with the target) | -4.01 | 1.15 mM | 1 hydrophobic alkyl/alkyl interaction between chlorine (furosemide) and -OH (ergosterol) |

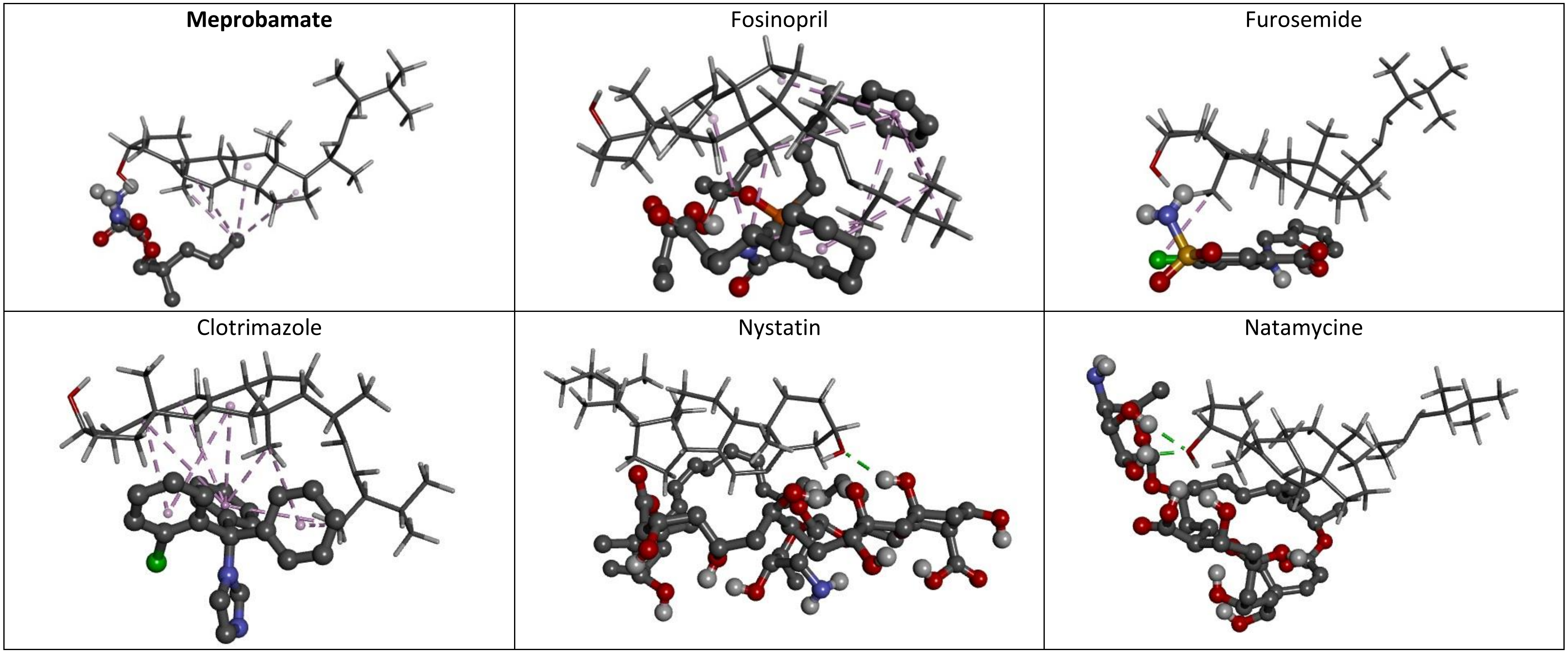


Figure S11. Structure views of the 3D-complexes between Meprobamate (i.e., the repositioning hint – $d_h^{\text{antifungal}}$ ), Clotrimazole (a $d_{25}^{\text{antifungal}}$ drug from $\mathscr{D}_{25}^{\text{antifungal}}$ reference drug with documented antifungal activity), Nystatin and Natamycine (these reference antifungals drugs $\in \mathscr{D}_{\overline{25}}^{\text{antifungal}}$ ), Fosinopril and Furosemide (reference drugs $\in \mathscr{D}_n^{\text{antifungal}}$ , with no reported antifungal activity) with Ergosterol, which is a steroidal target from $\mathscr{T}_{\overline{25}}^{\text{antifungal}}$ . The drug molecules are represented by the ball-and-stick molecular models, while the chemical structure in gray lines is for Ergosterol. The purple dashed lines represent the hydrophobic alkyl/alkyl interactions between Meprobamate, Clotrimazole, Fosinopril, and Furosemide with the target, and the green lines are for the hydrogen bonds of Nystatin and Natamycine with the target.

Table S12. The comparison of the molecular interactions of Meprobamate (i.e., the repositioning hint – $d_h^{\text{antifungal}}$ ), Ciclopirox (a $d_{\overline{25}}^{\text{antifungal}}$ drug from $\mathscr{D}_{\overline{25}}^{\text{antifungal}}$ reference drugs with documented antifungal activity), Fosinopril and Furosemide ($\in \mathscr{D}_n^{\text{antifungal}}$ reference drugs, with no reported antifungal activity) and Sodium/potassium-transporting ATPase subunit alpha (this target $\in \mathscr{T}_{\overline{25}}^{\text{antifungal}}$ ). There are no target amino acid involved in the same type of interaction with the test and reference drugs.

| $\mathscr{T}_{\overline{25}}^{\text{antifungal}}$ **Sodium/potassium-transporting ATPase subunit alpha** | | | | | | | | |
|---|---|---|---|---|---|---|---|---|
| Drug name | Drug role | Lowest free energy of binding [kcal/mol] | Estimated inhibition constant [Temp 298.15 K] | Conventional hydrogen bond | Carbon hydrogen bond | Alkyl interaction | Pi-Alkyl interaction | Interactive amino acid residues (Van der Waals interaction) |
| **Meprobamate** | Repositioning hint (No reported interaction with the target) | -2.47 | 15.37 mM | $GLN_{B69}$ $ASN_{B282}$ | $ALA_{B73}$ | $VAL_{B183}$ | $PHE_{B186}$ | $ASP_{B70}$, $VAL_{B72}$, $PRO_{B74}$, $PRO_{B75}$, $LEU_{B184}$, $GLY_{B185}$, $LYS_{B187}$, $GLU_{B281}$, $ILE_{B283}$ |
| Ciclopirox | Reference drug (binder) | -5.38 | 0.1145 mM | $GLN_{D69}$ $ALA_{D73}$ | | $ALA_{D73}$ $PRO_{D74}$ (3) | $PHE_{E19}$ $PHE_{D186}$ | $TYR_{E20}$, $TYR_{E21}$, $ASP_{D70}$, $ARG_{D71}$, $VAL_{D72}$, $PRO_{D75}$, $VAL_{D183}$, $GLU_{D281}$, $ASN_{D282}$, $ILE_{D283}$ |
| Fosinopril | Reference drug (No reported interaction with the target) | -3.04 | 5.92 mM | $SER_{C988}$ | | $VAL_{C928}$ (2) $VAL_{C937}$ $MET_{C942}$ (2) $CLR_{C1107}$ | | $VAL_{C921}$, $VAL_{C922}$, $TRP_{C924}$, $ALA_{C925}$, $LYS_{C931}$, $PHE_{C938}$, $ILE_{C948}$, $LEU_{C951}$, $PHE_{C952}$, $PHE_{C985}$, $LEU_{C989}$, $PHE_{C992}$ |
| Furosemide | Reference drug (No reported interaction with the target) | -4.11 | 0.9668 mM | $GLN_{C274}$ $THR_{C359}$ $LYS_{C719}$ $ASP_{C722}$ $ASP_{C740}$ (2) | $LEU_{C270}$ $GLU_{C271}$ $PRO_{C276}$ | $ALA_{C356}$ | | $GLY_{C272}$, $THR_{C275}$, $LYS_{C352}$, $ASN_{C353}$, $GLU_{C355}$, $SER_{C718}$, $ALA_{C721}$, $ILE_{C723}$, $GLY_{C724}$, $GLN_{C737}$, $ALA_{C738}$, $ALA_{C739}$ |

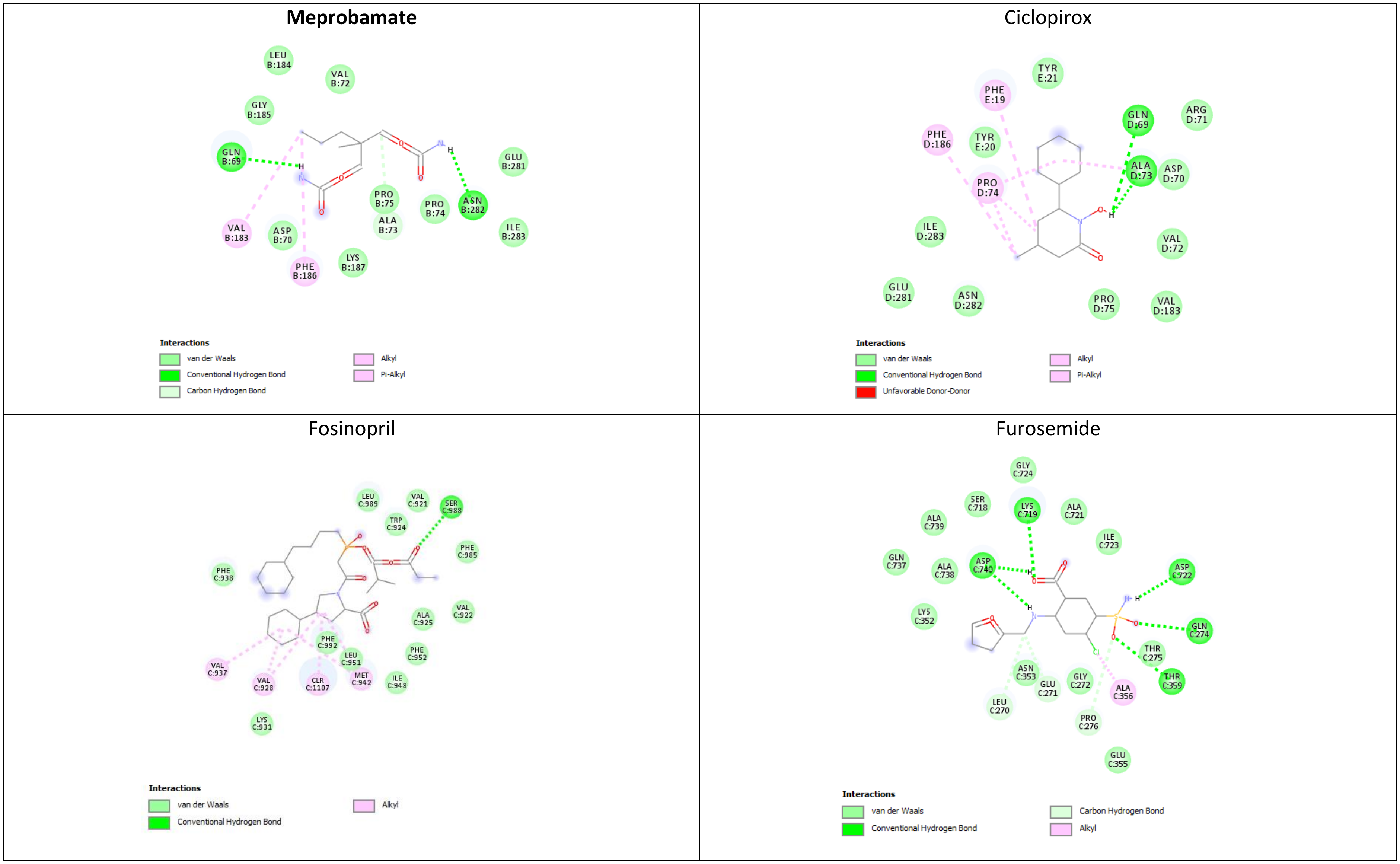


Figure S12. The 2D diagrams generated by the molecular docking simulation for interactions of Meprobamate, Fosinopril, and Furosemide (i.e., $d_t^{\text{antifungal}}$ drugs), Ciclopirox (i.e., a reference drug from $\mathscr{D}_{25}^{\text{antifungal}}$) with the amino acid residues in Sodium/potassium-transporting ATPase subunit alpha (this target $\in \mathscr{T}_{25}^{\text{antifungal}}$). The 2D chemical representation of the drug molecule is in the center of each square. The colored disks represent the amino acids surrounding the drug molecule, while the dotted lines represent the interactions between the target's amino acids and the drug molecule. The diagrams also indicate the amino acids that establish van der Waals interactions with the drug; however, for clarity, these interactions are not represented.

Table S13. The comparison of the molecular interactions between Meprobamate (i.e., the repositioning hint – $d_h^{\text{antifungal}}$ ), Griseofulvin (a $d_{\overline{25}}^{\text{antifungal}}$ drug from $\mathscr{D}_{\overline{25}}^{\text{antifungal}}$ reference drugs with documented antifungal activity), Fosinopril and Furosemide ($\in \mathscr{D}_n^{\text{antifungal}}$ reference drugs, with no reported antifungal activity), and Tubulin (this target $\in \mathscr{T}_{\overline{25}}^{\text{antifungal}}$ ). The residues shown in bold represent the Tubulin amino acids involved in the same type of interaction with the test and reference drugs.

| $\mathscr{T}_{\overline{25}}^{\text{antifungal}}$ **Tubulin** | | | | | | | | |
|---|---|---|---|---|---|---|---|---|
| Drug name | Drug role | Lowest free energy of binding [kcal/mol] | Estimated inhibition constant [Temp 298.15 K] | Conventional hydrogen bond | Carbon hydrogen bond | Alkyl interaction | Pi-Alkyl interaction | Interactive amino acid residues (Van der Waals interaction) |
| **Meprobamate** | Repositioning hint (No reported interaction with the target) | -2.99 | 6.42 mM | VAL$_{B260}$ TRP$_{B346}$ | | PRO$_{B261}$ **ILE$_{B347}$** | | ALA$_{B256}$, VAL$_{B257}$, **ASN$_{B258}$**, **MET$_{B259}$**, **LEU$_{B313}$**, THR$_{B314}$, **PRO$_{B348}$**, **ASN$_{B349}$**, LYS$_{A401}$, **ALA$_{A403}$**, **PHE$_{A404}$**, **TYR$_{B435}$** |
| Griseofulvin | Reference drug (inhibitor) | -6.14 | 0.0316 mM | THR$_{B314}$ LYS$_{A401}$ | VAL$_{A181}$ VAL$_{B257}$ | **ILE$_{B347}$** | | VAL$_{A182}$, PRO$_{A184}$, **ASN$_{B258}$**, **MET$_{B259}$**, VAL$_{B260}$, PRO$_{B261}$, **LEU$_{B313}$**, TRP$_{B346}$, **PRO$_{B348}$**, **ASN$_{B349}$**, ASN$_{B350}$, LEU$_{A397}$, MET$_{A398}$, **ALA$_{A403}$**, **PHE$_{A404}$**, TYR$_{B432}$, **TYR$_{B435}$** |
| Fosinopril | Reference drug (No reported interaction with the target) | -4.28 | 0.7279 mM | THR$_{B145}$ GLY$_{B146}$ ASP$_{B179}$ THR$_{B180}$ GTP$_{B502}$ | THR$_{B145}$ | PRO$_{B173}$ | HIS$_{B139}$ | ALA$_{B9}$, GLY$_{B10}$, CYS$_{B12}$, ASN$_{B101}$, LEU$_{B137}$, GLY$_{B142}$, GLY$_{B143}$, GLY$_{B144}$, SER$_{B147}$, GLY$_{B150}$, SER$_{B170}$, VAL$_{B171}$, VAL$_{B172}$, GLU$_{B183}$, ASN$_{B206}$ |
| Furosemide | Reference drug (No reported interaction with the target) | -5.09 | 0.1861 mM | HIS$_{A309}$ GLN$_{A342}$ (2) | LYS$_{A311}$ | | | PHE$_{A296}$, PRO$_{A307}$, ARG$_{A308}$, GLY$_{A310}$, TYR$_{A312}$, ILE$_{A335}$, LYS$_{A338}$, THR$_{A340}$, ILE$_{A341}$, PHE$_{A343}$ |

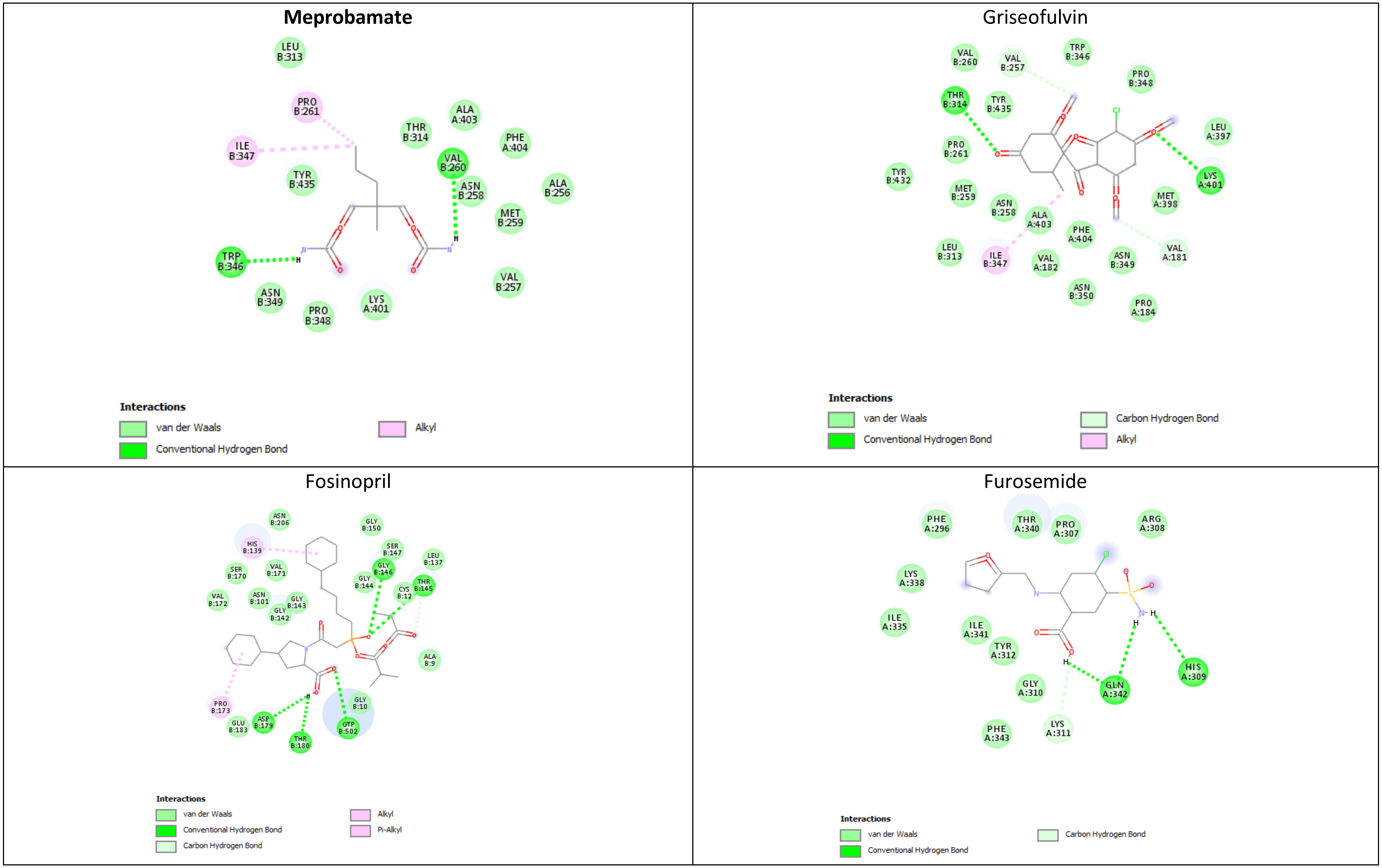


Figure S13. The 2D maps generated by the molecular docking simulation for interactions between Meprobamate, Fosinopril, Furosemide (i.e., $d_t^{\text{antifungal}}$ drugs), Griseofulvin (this reference drug $\in \mathscr{D}_{25}^{\text{antifungal}}$) and the amino acid residues in Tubulin (this target $\in \mathscr{T}_{25}^{\text{antifungal}}$). The 2D chemical representation of the drug molecule is in the center of each square. The colored disks represent the amino acids surrounding the drug molecule, while the dashed lines represent the interactions between the target's amino acids and the drug molecule. The maps also indicate the amino acids that establish van der Waals interactions with the drug; however, for clarity, these interactions are not represented.

### 4. Molecular docking results interpretation

#### 4.1. Azelaic acid

**Table S1 & Figure S1**

The free energy of the complex between Progesterone and **Estrogen receptor alpha** is -5.89 kcal/mol, Abiraterone and **Estrogen receptor alpha** is -7.11 kcal/mol, whereas that of the complex between Azelaic acid and Estrogen receptor alpha is -4.48 kcal/mol, showing that the Progesterone and Abiraterone complexes have higher stability than that of the Azelaic acid complex. Azelaic acid and Progesterone bind to the target through 8 amino acids, but Azelaic acid establishes an identical type of interaction with only one amino acid. Azelaic acid and Abiraterone bind to the target through 7 amino acids, but Azelaic acid establishes an identical type of interaction with only one amino acid. Fosinopril and Furosemide interact with no amino acids in the target. The results indicate a low similarity between Progesterone (i.e., a reference drug in $\mathscr{D}_6^{\text{anticancer}}$) and $d_h^{\text{anticancer}}$ Azelaic acid. We also notice a clear difference between the $\mathscr{D}_n^{\text{anticancer}}$ = {Fosinopril, Furosemide} and reference anticancer drugs in terms of interaction at the active binding site of Estrogen receptor beta.

**Table S2 & Figure S2**

The lowest free energy of the Azelaic acid-**Estrogen receptor beta** complex is -3.11 kcal/mol, Abiraterone-Estrogen receptor beta complex is -7.90 kcal/mol, and of the Progesterone-Estrogen receptor beta complex is -8.68 kcal/mol, indicating that the Progesterone and Abiraterone complexes have higher stability than that of Azelaic acid. Both Progesterone and Azelaic acid bind the same 15 amino acids, and 5 out of 15 interactions are of the same type. Azelaic acid and Abiraterone bind to the target through the same 4 amino acids, and Azelaic acid establishes identical type of interactions with 3 amino acids. Fosinopril and Furosemide interact with no amino acid in the active site of the Estrogen receptor beta.

**Table S3 & Figure S3**

The free energy of the complex between Azelaic acid and **Progesterone receptor** is -4.54 kcal/mol, that of the complex between Progesterone and Progesterone receptor is -11.17 kcal/mol, and that of the complex between Abiraterone and Progesterone receptor is -11.90 kcal/mol, thus indicating higher stability for the Abiraterone and Progesterone complexes. However, Azelaic acid and Progesterone similarly bind to the target, as both drugs interact with the same 12 amino acids in the target, and 7 out of 12 interactions are of the same type; Azelaic acid and Abiraterone bond to this target with the same 7 amino acids, but only 2 out of these 7 interactions are of the same type. Fosinopril establishes only one van der Waals interaction with Progesterone receptor, and Furosemide do not interact with any amino acids in the target. These results indicate a similarity between Progesterone (i.e., a reference drug $\mathscr{D}_6^{\text{anticancer}}$) and $d_h^{\text{anticancer}}$ Azelaic acid. On the other hand, we notice a clear difference between the $\mathscr{D}_n^{\text{anticancer}}$ = {Fosinopril, Furosemide} and Progesterone in terms of interaction at the active binding site of Progesterone receptor.

**Table S4 & Figure S4**

The lowest free energy of the complex between Azelaic acid and **Steroid 17-alpha-hydroxylase/17,20 lyase** is -8.49 kcal/mol, that of the complex between Progesterone and Steroid 17-alpha-hydroxylase/17,20 lyase is -8.72 kcal/mol, and that of the complex between Abiraterone and Steroid 17-alpha-hydroxylase/17,20 lyase is -8.99 kcal/mol, suggesting a very similar stability of the three complexes. DrugBank lists Progesterone as a substrate and inhibitor, and Abiraterone as an inhibitor of Steroid 17-alpha-hydroxylase/17,20 lyase. On the other hand, our results indicate a high similarity between the repositioning hint (i.e., Azelaic acid) and the reference anticancer drugs (i.e., Progesterone and Abiraterone) in terms of the inhibition constant, as these values rendered by the docking simulation are 600.71 nM, 406.22 nM, and 402.33 nM for Azelaic acid, Progesterone, and Abiraterone, respectively. Azelaic acid and Progesterone similarly bind to the target, as both drugs interact with the same 8 amino acids in the target, and 5 out of 8 interactions are of the same type. Azelaic acid and Abiraterone interact with the same 5 amino acids in the target, and 2 interactions are of the same type. Moreover, our docking simulation results are in line with results of C. Avendaño (Carme nAvendaño, J. CarlosMenéndez. Chapter 3 - Anticancer Drugs That Modulate Hormone Action. Medicinal Chemistry of Anticancer Drugs, Second Edition, 2015, p. 81-131) and **N.M. DeVore** (Natasha M. DeVore & Emily E. Scott. Structures of cytochrome P450 17A1 with prostate cancer drugs abiraterone and TOK-001. Nature. 2012; 482(7383): 116–119), which report the covalent bonding of Abiraterone and Steroid 17-alpha-hydroxylase/17,20 lyase (a cysteinato-heme enzyme that belongs to the cytochrome P450 superfamily). Precisely, Abiraterone forms a coordinate covalent bond of the pyridine nitrogen at $C_{17}$ with heme iron of this target (Natasha M. DeVore & Emily E. Scott. Structures of cytochrome P450 17A1 with prostate cancer drugs abiraterone and TOK-001. Nature. 2012; 482(7383): 116–119). Fosinopril and Furosemide do not interact with any amino acids in this target. We notice a similarity between Progesterone (i.e., a reference drug $\mathscr{D}_{6}^{\text{anticancer}}$ ) and $d_{h}^{\text{anticancer}}$ Azelaic acid in terms of number and type of interactions with the amino acids in the target. We notice a clear difference between the $\mathscr{D}_{n}^{\text{anticancer}}$ = {Fosinopril, Furosemide} and the anticancer reference drugs (i.e., Progesterone and Abiraterone) in terms of interaction at the active binding site of Steroid 17-alpha-hydroxylase/17,20 lyase. These results suggest that Azelaic acid is a promising candidate for further in silico, in vitro and in vivo investigations of its potential anticancer effects.

**Table S5 & Figure S5**

The lowest free energy of the Azelaic acid - **Androgen receptor** complex is -5.01 kcal/mol, whereas the Progesterone - Anrogen receptor complex is more stable than the Azelaic acid - Androgen receptor complex (-10.28 kcal/mol). Azelaic acid and Progesterone interact similarly with Androgen receptor through a common set of 10 amino acids; 6 out of these interactions are of the same type. Fosinopril and Furosemide interact with none of the 11 amino acids in the active site of the target. Again, we notice a difference between the $\mathscr{D}_{n}^{\text{anticancer}}$ = {Fosinopril, Furosemide} and Progesterone in terms of interactions at the active binding site of Androgen receptor.

**Table S6 & Figure S6**

The lowest free energy of the Azelaic acid - **Mineralocorticoid receptor** complex is -4.49 kcal/mol, of the Progesterone - Mineralocorticoid receptor complex is -10.77 kcal/mol, and of the Abiraterone - Mineralocorticoid receptor is -8.08 kcal/mol, thus indicating that the Progesterone complex has higher stability than that of Abiraterone and Azelaic acid complexes. Progesterone and Azelaic acid bind the same 13 amino acids in the target, and 4 out of 13 interactions are of the same type. Abiraterone, Fosinopril and Furosemide interact with no amino acid in the active site of the Mineralocorticoid receptor.

### 4.2 Meprobamate

**Table S7 & Figure S7**

The free energy of the complex between Meprobamate and **Lanosterol 14-alpha demethylase** is -2.77 kcal/mol, whereas that of the complex between Clotrimazole and Lanosterol 14-alpha demethylase is -7.15 kcal/mol, showing a higher stability for the Clotrimazole complex. However, Clotrimazole and Meprobamate similarly bind to the target, as both drugs interact with the same 10 amino acids in the target (7 out of 10 interactions are of the same type). Conversely, Fosinopril and Furosemide interact with none of these 10 amino acids in the target. On the other hand, we notice a clear difference between the $\mathscr{D}_n^{\text{antifungal}}$ = {Fosinopril, Furosemide} and Clotrimazole in terms of interaction at the active binding site of Lanosterol 14-alpha demethylase.

**Table S8 & Figure S8**

The lowest free energy of the Meprobamate - **Lanosterol synthase** complex is -3.23 kcal/mol, whereas the Oxiconazole - Lanosterol synthase complex is more stable than the Meprobamate-Lanosterol synthase (-6.24 kcal/mol). Oxiconazole and Meprobamate interact similarly with Lanosterol synthase through a common set of 11 amino acids, interactions of which 7 are of the same type. Fosinopril and Furosemide interact with none of the 11 amino acids in the active site of the target. Again, we have a clear difference between the $\mathscr{D}_n^{\text{antifungal}}$ = {Fosinopril, Furosemide} and Oxiconazole regarding the interaction at the active binding site of Lanosterol synthase.

**Table S9 & Figure S9**

The molecular docking results reveal that the complex Meprobamate - **Intermediate conductance calcium-activated potassium channel protein 4** has the lowest free energy of -1.02 kcal/mol. The complex between the reference antifungal drug Clotrimazole and the same target has the lowest free energy -3.60 kcal/mol. Examining the drug-target interaction, we notice that Meprobamate, Fosinopril, and Furosemide are different from Clotrimazole in the way they interact with the amino acids in the target active site.

**Table S10 & Figure S10**

For the Meprobamate - **Squalene monooxygenase** complex, the lowest free energy is -2.67 kcal/mol, while for the complexes of Tolnaftate and Naftifine with Squalene monooxygenase are -6.75 kcal/mol and 7.47 kcal/mol, respectively; this means that the reference drug-target complexes are more stable than the Meprobamate complex. Meprobamate and Tolnaftate interact with the same 15 amino acids in the target (of which 5 are of the same type). Concurrently, the interaction of both Meprobamate and Naftifine with Squalene monooxygenase shares only 3 amino acids, of which 2 are of the same type. Fosinopril and Furosemide interact with no amino acid in the active site of the target. According to our docking results, Meprobamate's behavior in terms of binding to the target Squalene monooxygenase is more similar to Tolnaftate than to Naftifine.

**Table S11 & Figure S11**
The interaction with the (non-protein) target **Ergosterol** is similar for Meprobamate (the repositioning hint) and Clotrimazole ($\in \mathscr{D}^{antifungal}_{25}$), through weak hydrophobic interactions with the Ergosterol six-atoms cycles. The Clotrimazole-Ergosterol complex (-4.06 kcal/mol) is more stable than the Meprobamate-Ergosterol complex (-3.48 kcal/mol). Of note, the reference antifungal drugs Nystatin and Natamycine ($\mathscr{D}^{antifungal}_{\overline{25}}$) interact differently with Ergosterol, as they form hydrogen bonds with the target's hydroxyl group, while Fosinopril and Furosemide establish hydrophobic interactions with the five-atom cycle of Ergosterol.

**Table S12 & Figure S12**
The molecular docking of Meprobamate with **Sodium/potassium-transporting ATPase subunit alpha**, as well as of Ciclopirox with the same target, reveals that the former complex has lower stability (-2.47 kcal/mol) than the latter (-5.38 kcal/mol). Furthermore, Meprobamate and Ciclopirox do not interact with common amino acids in the active site of the target. Fosinopril and Furosemide do not interact with the amino acids in the active site of the target.

**Table S 13 & Figure S13**
The lowest free energy of the Meprobamate - **Tubulin** complex is -2.99 kcal/mol, and of the Griseofulvin - Tubulin complex is -6.14 kcal/mol, indicating that the Griseofulvin complex has much higher stability than that of Meprobamate. Griseofulvin and Meprobamate similarly bind to the target, as both drugs interact with the same 15 amino acids (9 out of 15 interactions are of the same type). Fosinopril and Furosemide interact with none of the 15 amino acids in the active site of the target.

### 5. The graphical representations of the docked complexes

**Figure 5.1.a.** The graphical representation of the docked complex between the antifungal reference drug Clotrimazole and the target Lanosterol 14-alpha demethylase

**Figure 5.1.b.** The graphical representation of the docked complex between the antifungal repositioning hint Meprobamate and the target Lanosterol 14-alpha demethylase

**Figure 5.2.a.** The graphical representation of the docked complex between the anticancer reference drug Progesterone and the target Progesterone receptor

**Figure 5.2.b.** The graphical representation of the docked complex between the anticancer repositioning hint Azelaic acid and the target Progesterone receptor

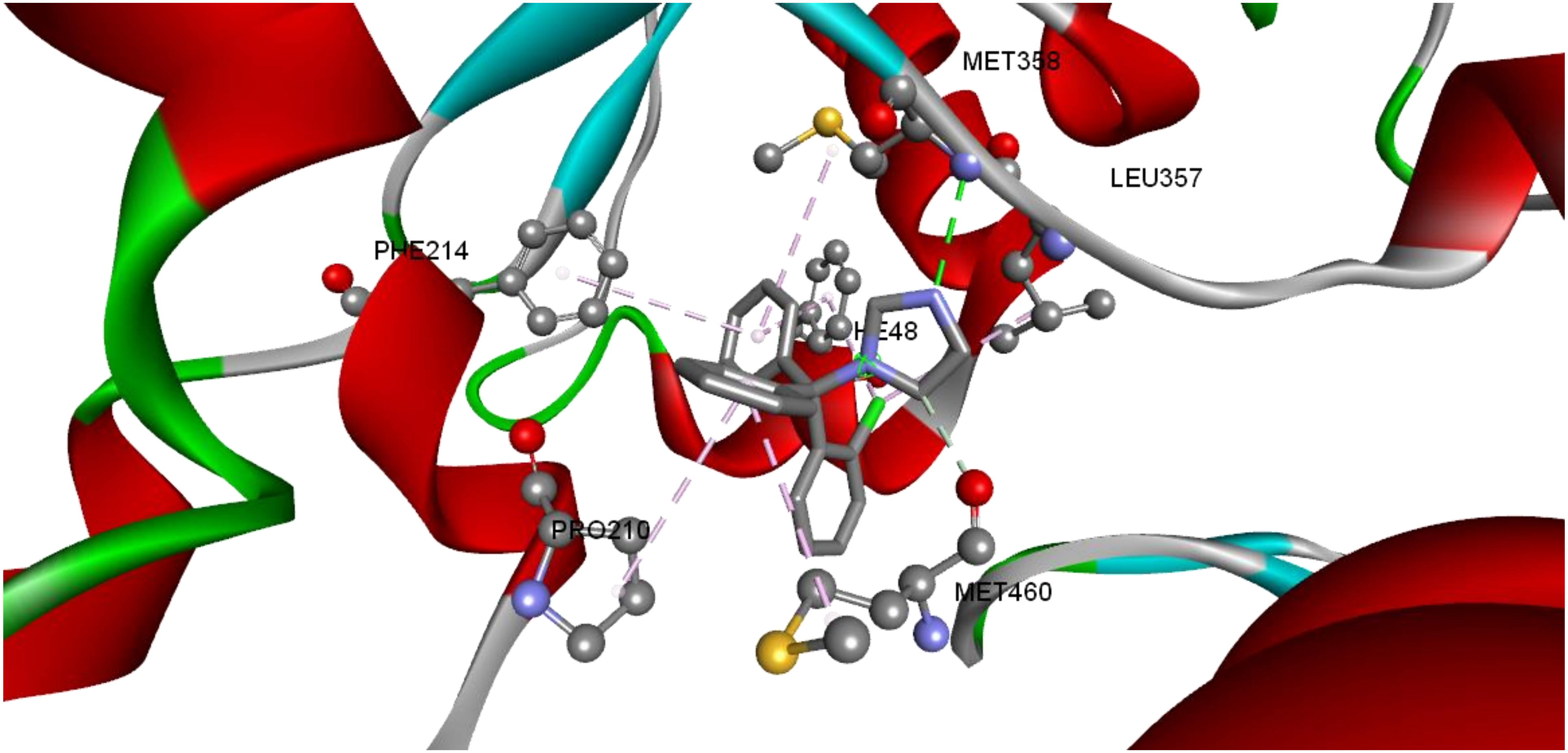


**Figure 5.1.a.** Molecular interactions analysis of Clotrimazole (the reference antifungal drug) with target Lanosterol 14-alpha demethylase. The docked complex of Lanosterol 14-alpha demethylase with Clotrimazole emphasize the molecular interactions of Meprobamate towards the active site of Lanosterol 14-alpha demethylase; the green dashed lines represent conventional and carbon hydrogen bonds, and the pink dashed lines represent the alkyl and pi-alkyl interactions. The flat ribbon represents the protein target, from which the interacting amino acid residues append. The figure displays the drug molecules (figured as ball-and-stick models) in the binding pocket of the target.

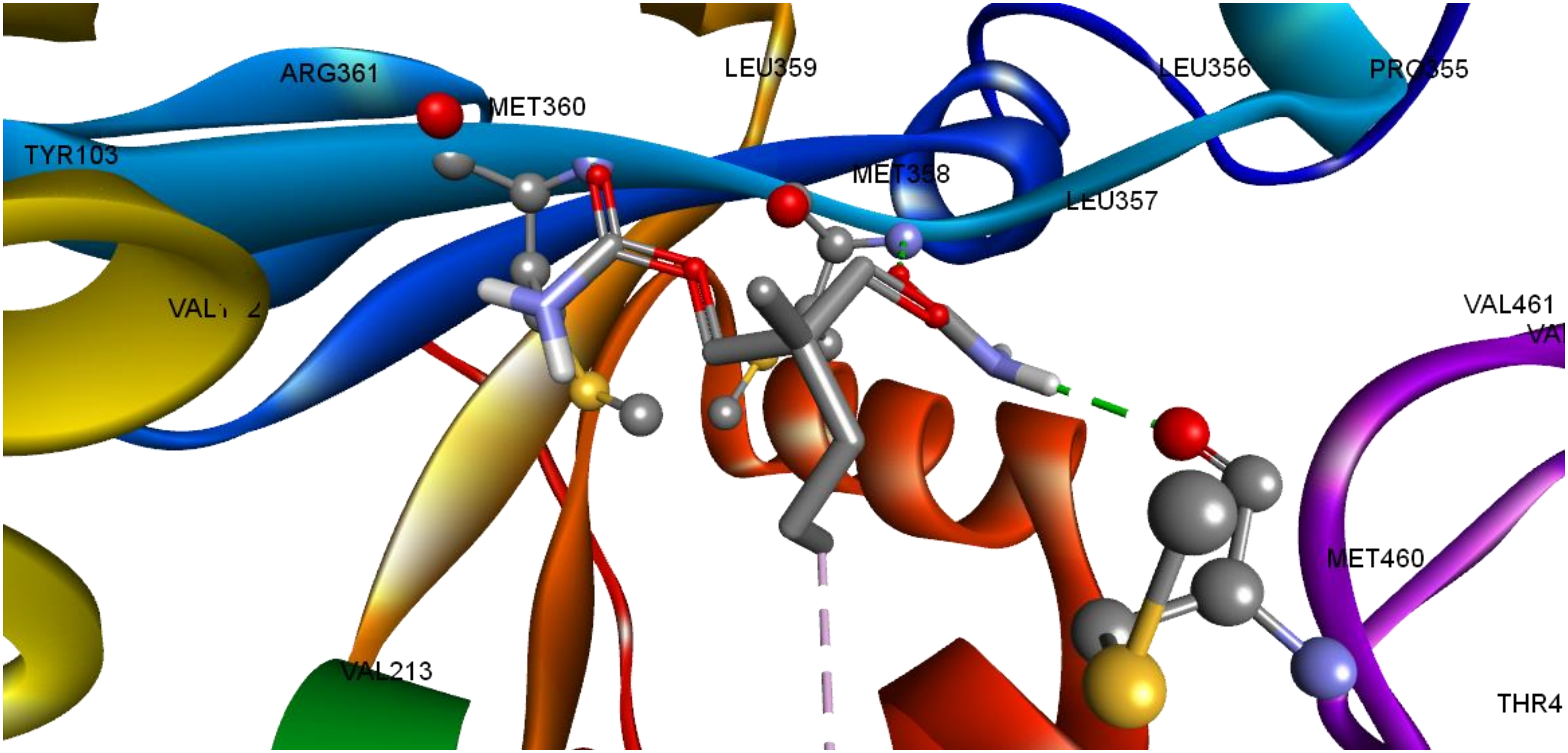


**Figure 5.1.b.** Molecular interactions analysis of Meprobamate (the repositioning hint) with target Lanosterol 14-alpha demethylase. The docked complex of Lanosterol 14-alpha demethylase with Meprobamate emphasize the molecular interactions of Meprobamate towards the active site of Lanosterol 14-alpha demethylase. The green dashed lines represent conventional hydrogen bonds, and the pink dashed lines represent the alkyl interactions. The flat ribbon represents the protein target, from which the interacting amino acid residues append. The figure displays the drug molecule (figured as ball-and-stick models) in the binding pocket of the target.

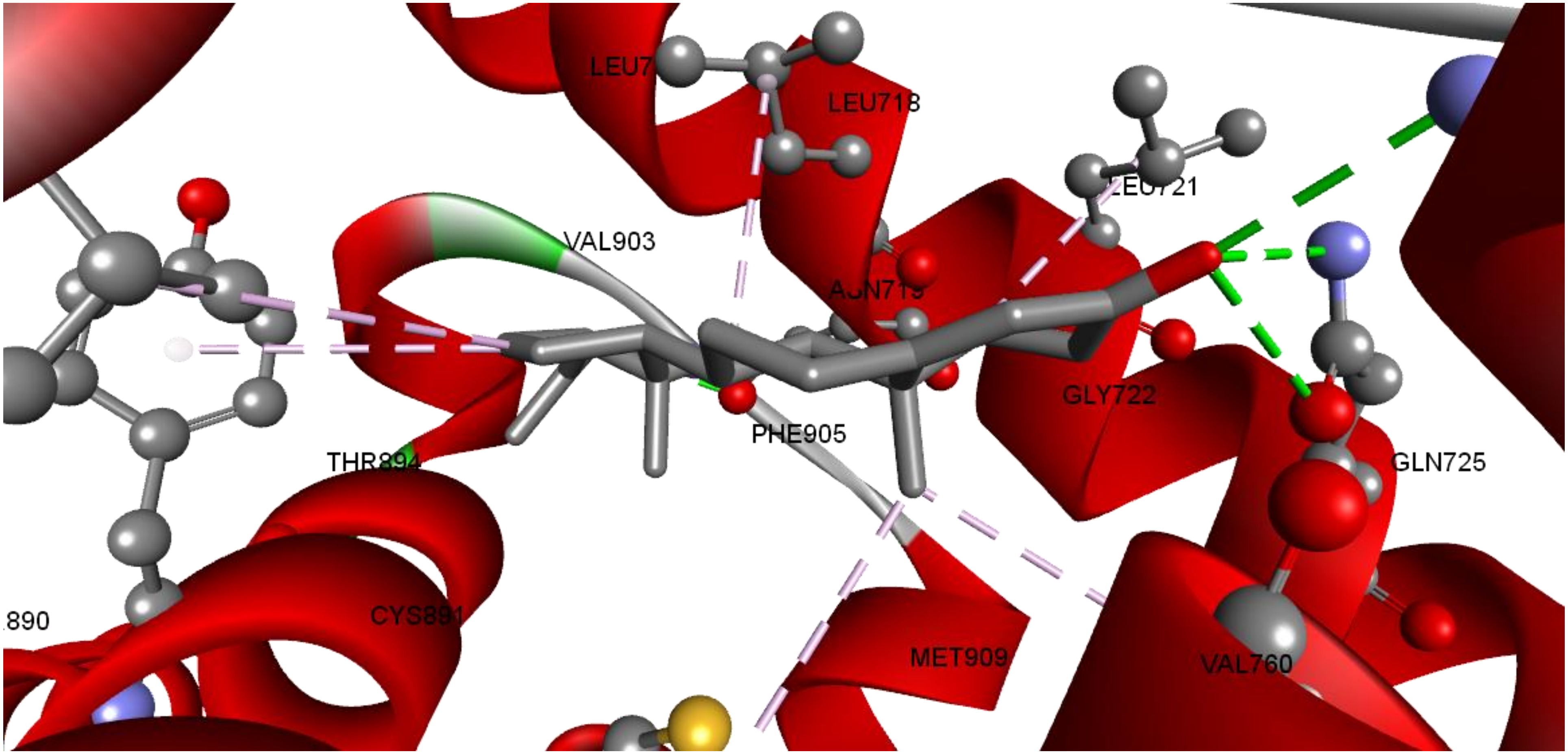


**Figure 5.2.a.** Molecular interactions analysis of Progesterone (the reference anticancer drug) with target Progesterone receptor. The docked complex of Progesterone receptor with Progesterone emphasize the molecular interactions of Progesterone towards the active site of Progesterone receptor; the green dashed lines represent conventional hydrogen bonds, and the pink dashed lines represent the alkyl and pi-alkyl interactions. The flat ribbon represents the protein target, from which the interacting amino acid residues append. The figure displays the drug molecules (figured as ball-and-stick models) in the binding pocket of the target.

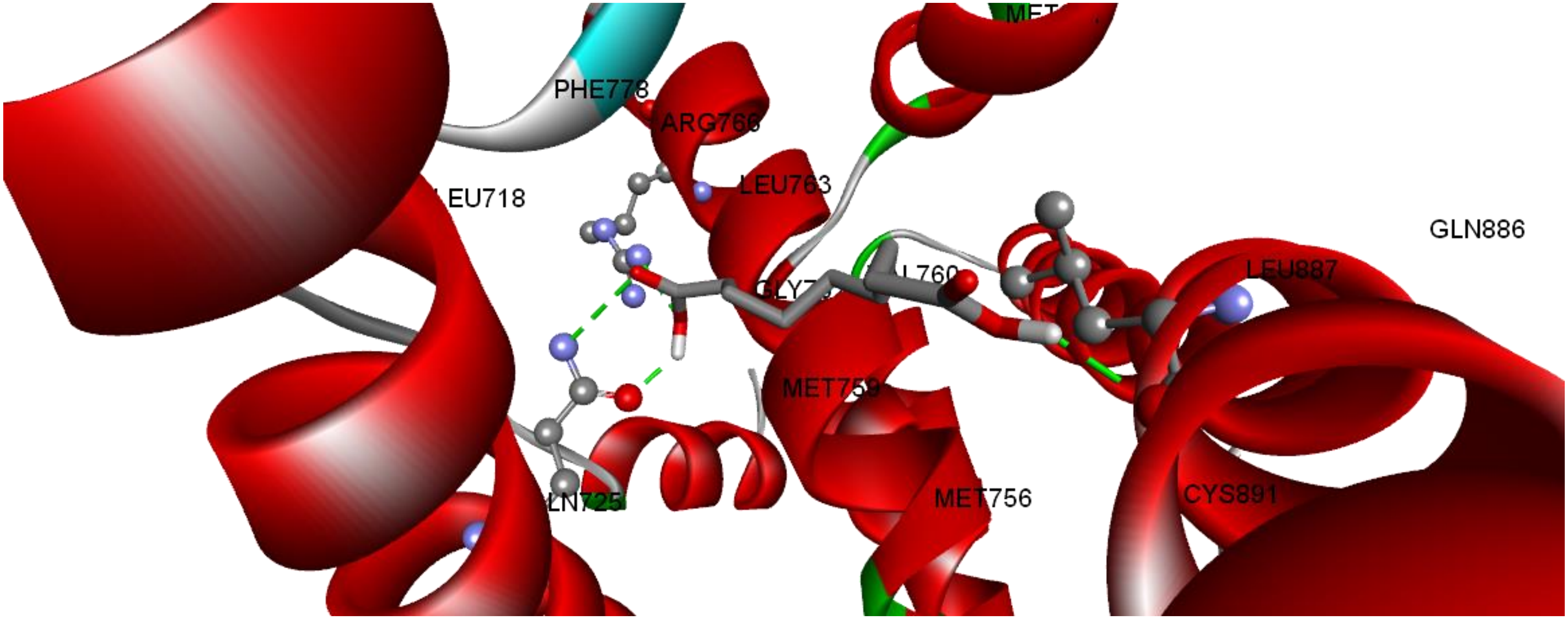


**Figure 5.2.b.** Molecular interactions analysis of Azelaic acid (the repositioning hint) with target Progesterone receptor. The docked complex of Progesterone receptor with Azelaic acid emphasize the molecular interactions of Azelaic acid towards the active site of Progesterone receptor. The green dashed lines represent conventional hydrogen bonds. The flat ribbon represents the protein target, from which the interacting amino acid residues append. The figure displays the drug molecule (figured as ball-and-stick models) in the binding pocket of the target.

### 6. Quantum chemical calculation

### 6.1 HOMO-LUMO energies for all ligands (i.e., drugs) in this manuscript

| Graphical representation of HOMO orbitals in the studied molecules | Graphical representation of LUMO orbitals in the studied molecules | $E_{HOMO}$ (eV) | $E_{LUMO}$ (eV) | $\Delta E$ [(1)] (eV) | $\lambda$ [(2)] (eV) | $\eta$ [(3)] (eV) |
|---|---|---|---|---|---|---|
| **Azelaic acid** | | | | | | |
| 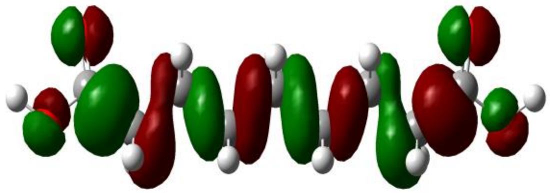 | 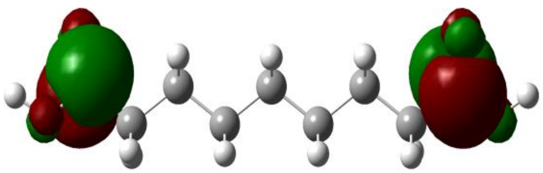 | - 11.38467 | 0.917639 | 12.302309 | 5.2335 | 6.15115 |
| **Progesterone** | | | | | | |
| 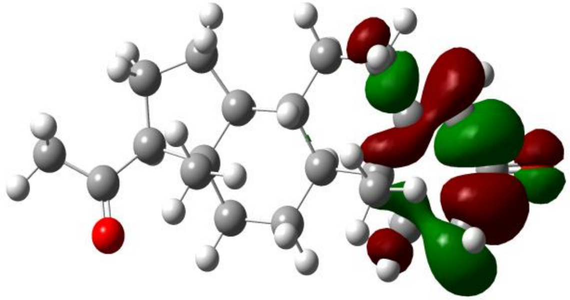 | 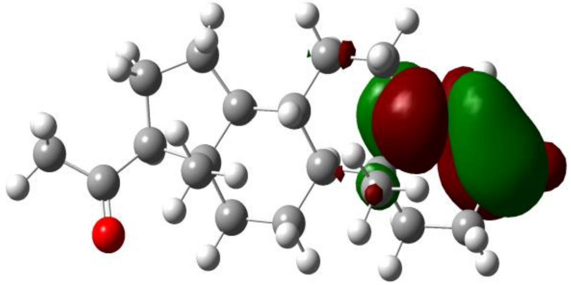 | - 10.15986 | - 0.097347 | 10.062512 | 5.1286 | 5.031255 |
| **Abiraterone** | | | | | | |
| 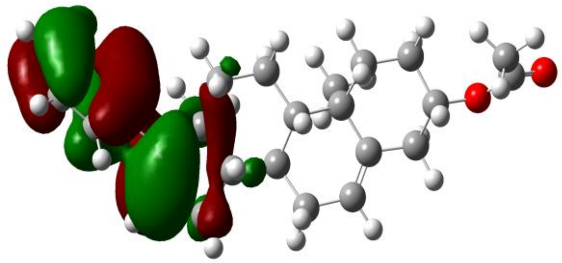 | 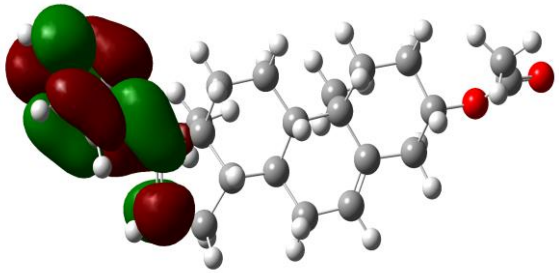 | - 9.249274 | - 0.309326 | 8.939947 | 4.7793 | 4.469973 |

| | | | | | | |
|---|---|---|---|---|---|---|
| **Meprobamate** | | | | | | |
| 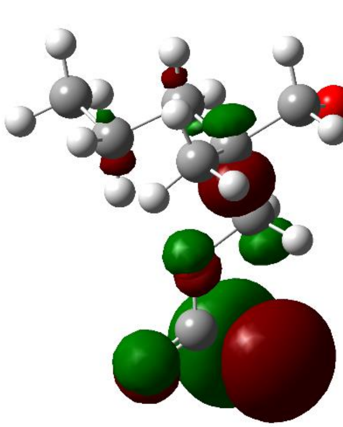 | 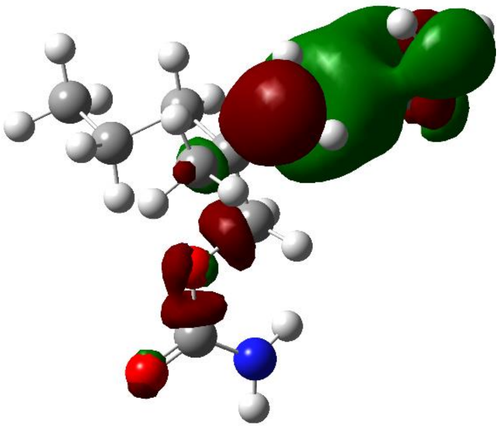 | **-10.48988** | **0.867030** | **11.35691** | **4.811426** | **5.6784** |
| **Clotrimazole** | | | | | | |
| 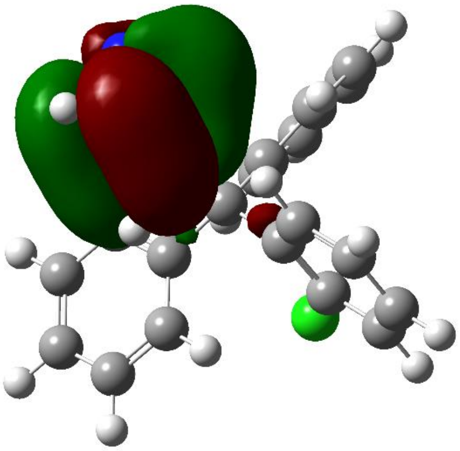 | 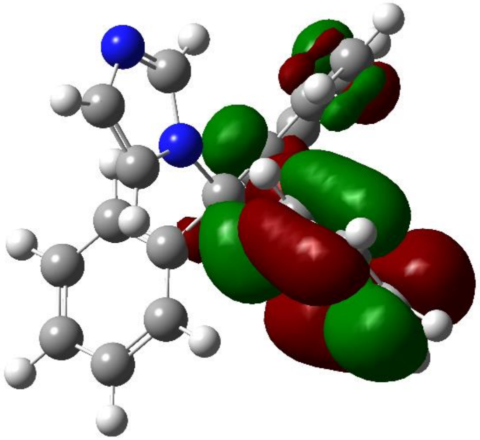 | **- 9.280731** | **- 0.347823** | **8.932907** | **4.814275** | **4.292541** |

| | | | | | | |
|---|---|---|---|---|---|---|
| **Oxiconazole** | | | | | | |
| 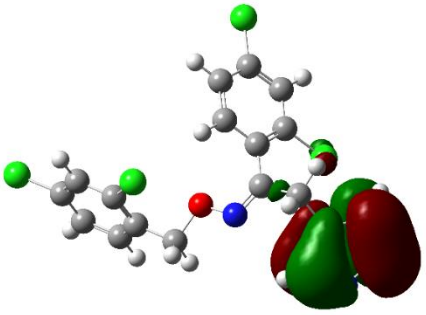 | 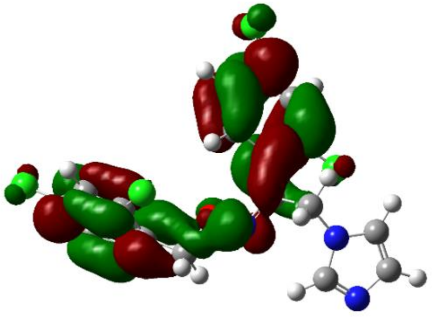 | **- 9.344505** | **- 0.631010** | **8.713494** | **4.987755** | **4.356747** |
| **Naftifine** | | | | | | |
| 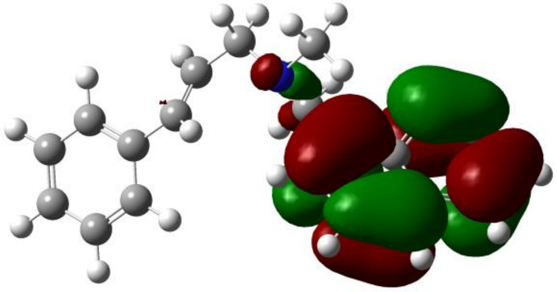 | 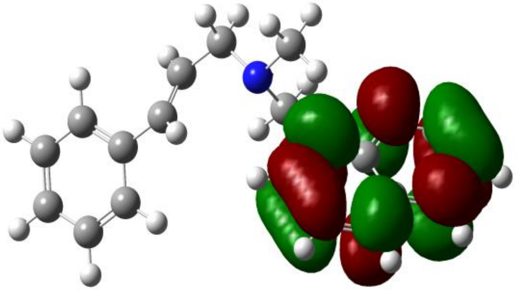 | **- 8.649096** | **- 0.351178** | **8.297918** | **4.50013** | **4.148959** |
| **Tolnaftate** | | | | | | |
| 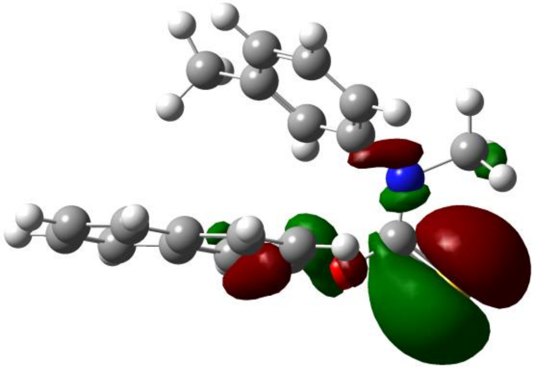 | 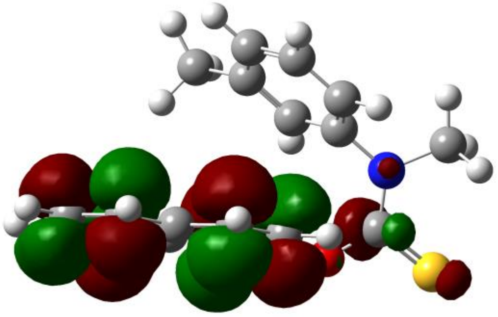 | **- 8.685643** | **- 0.760219** | **7.925423** | **4.72293** | **3.962711** |

| | | | | | | |
|---|---|---|---|---|---|---|
| **Nystatin** | | | | | | |
| 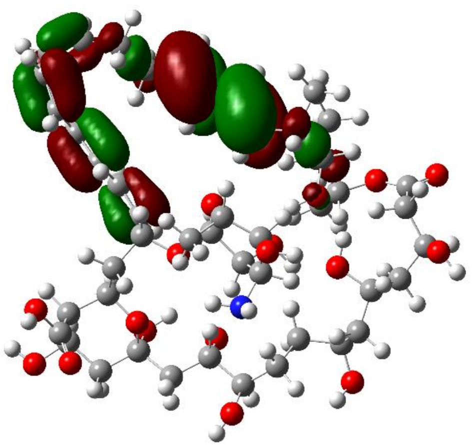 | 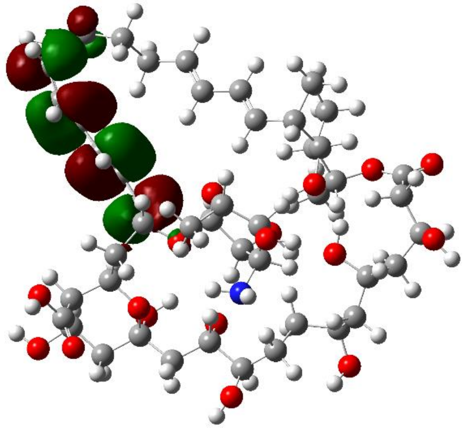 | **- 8.714139** | **- 0.626526** | **8.087619** | **4.67033** | **3.7305** |
| **Natamycin** | | | | | | |
| 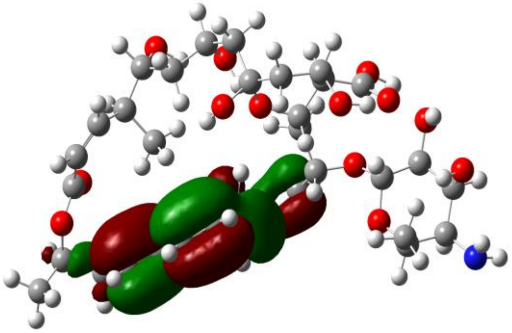 | 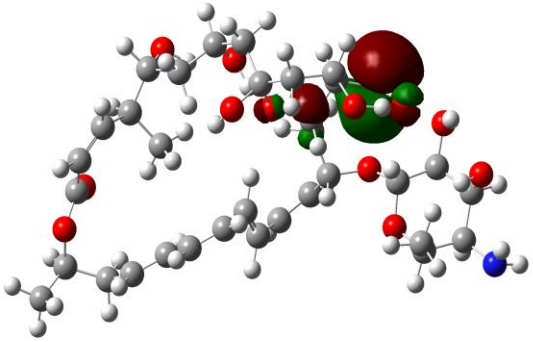 | **- 9.110476** | **0.0363716** | **9.146847** | **4.53705** | **4.573423** |
| **Ciclopirox** | | | | | | |
| 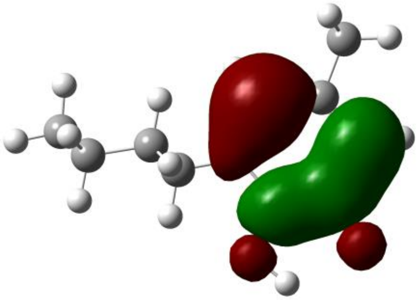 | 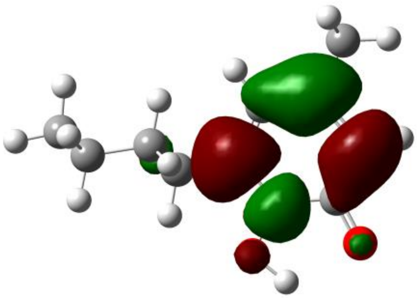 | **- 8.864879** | **- 0.518489** | **8.346389** | **4.69164** | **4.173194** |

| | | | | | | |
|---|---|---|---|---|---|---|
| **Griseofulvin** | | | | | | |
| 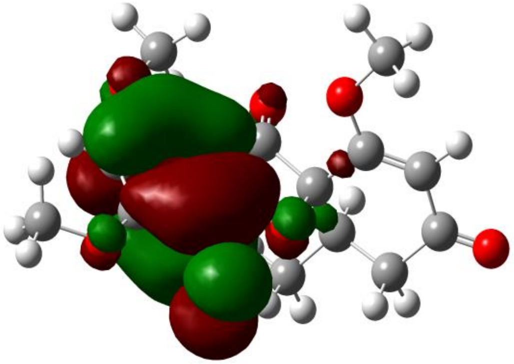 | 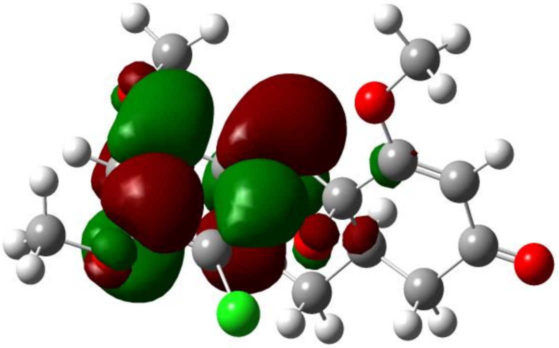 | **- 9.227333** | **- 0.832985** | **8.394347** | **5.03015** | **3.780681** |
| **Fosinopril** | | | | | | |
| 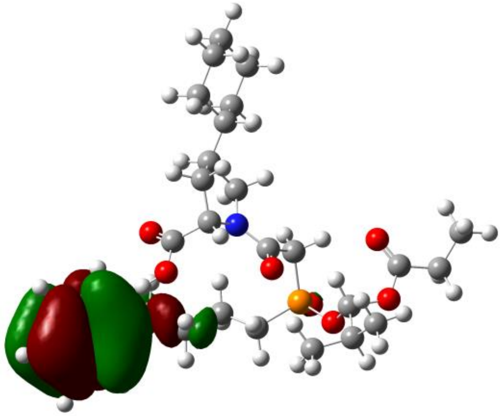 | 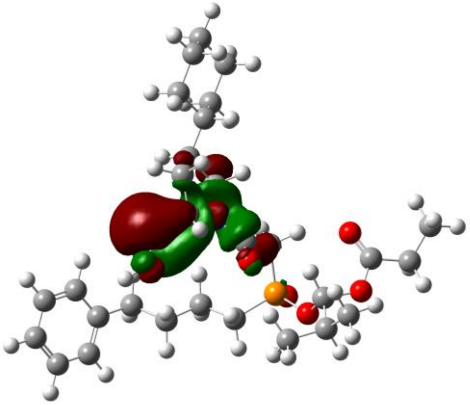 | **- 9.52807** | **- 0.088852** | **9.439217** | **4.80846** | **4.719608** |
| **Furosemide** | | | | | | |
| 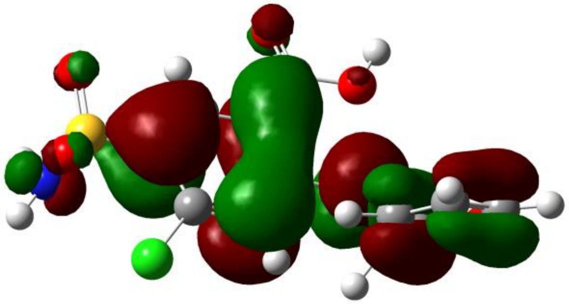 | 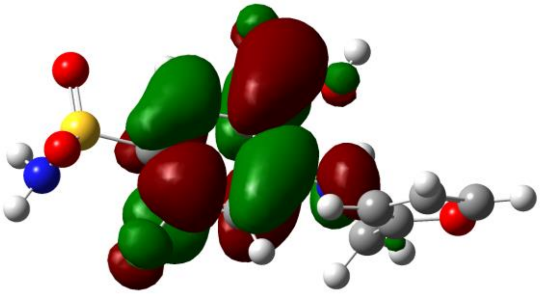 | **- 9.450583** | **- 0.988114** | **8.462468** | **5.2193** | **4.231234** |

The HOMO and LUMO energies are calculated in eV by DFT B3LYP.

Green and dark red isosurfaces of HOMO and LUMO indicate positive and negative values, respectively.

(1) $\Delta E = E_{LUMO} - E_{HOMO}$

(2) $\lambda = \frac{E_{LUMO} + E_{HOMO}}{2}$

(3) $\eta = \frac{E_{LUMO} - E_{HOMO}}{2}$

### *6.2* Mulliken population analysis for partial atomic charges for all ligands (*i.e.*, drugs) in this manuscript

| **Meprobamate** | **Ciclopirox** | **Clotrimazole** |
|---|---|---|
| 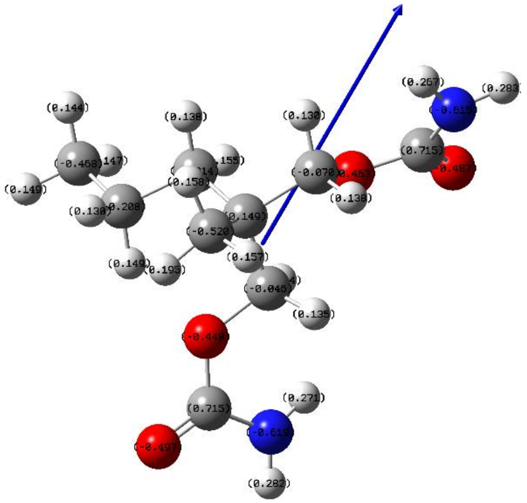 | 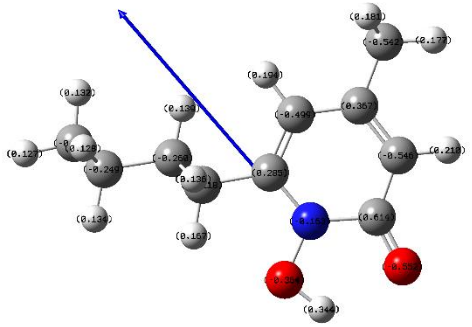 | 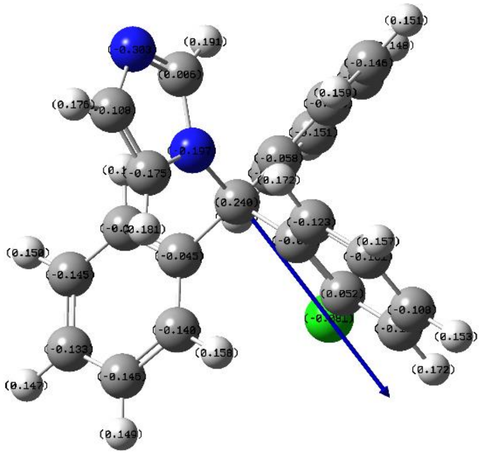 |

| Griseofulvine | Naftifine | Natamycin |
| --- | --- | --- |
| 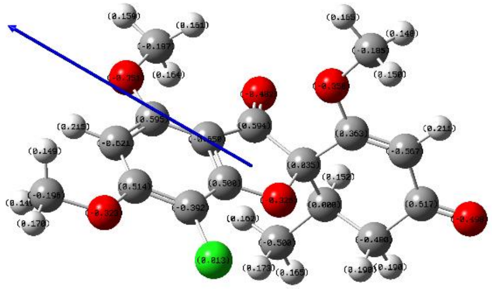 | 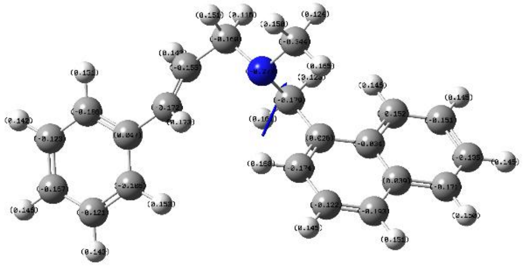 | 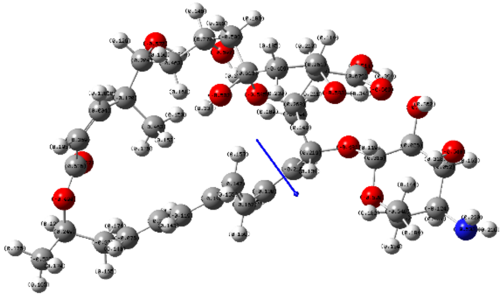 |
| **Nystatin** | **Oxiconazole** | **Tolnaftate** |
| 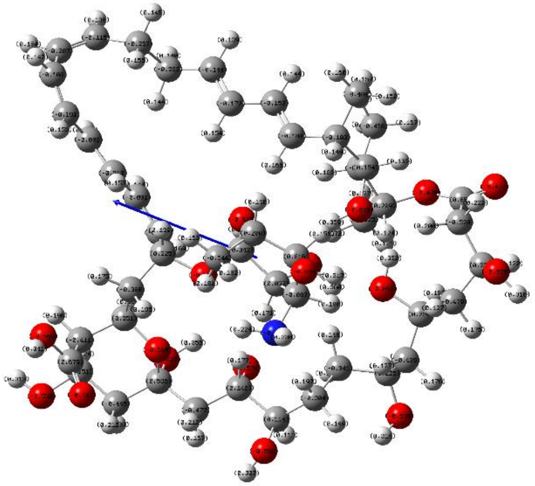 | 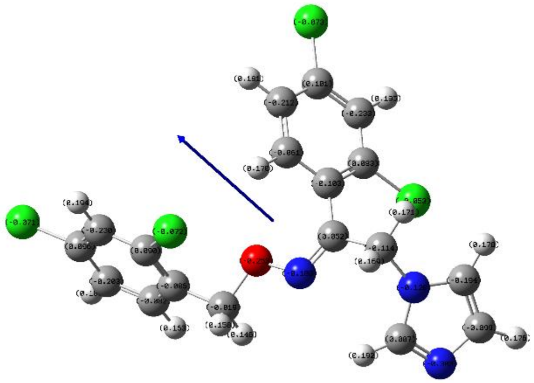 | 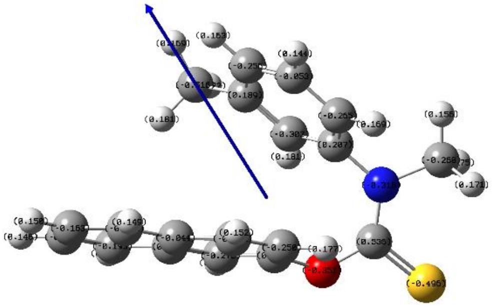 |

| Azelaic acid | Abiraterone | Progesterone |
|---|---|---|
| 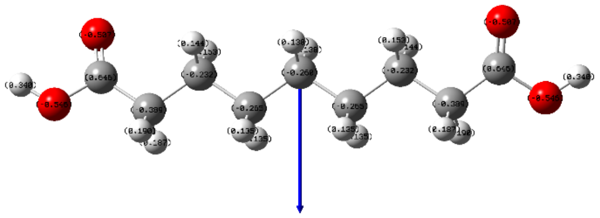 | 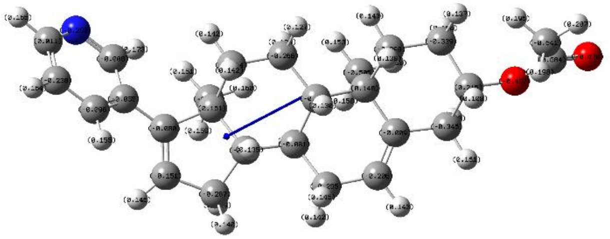 | 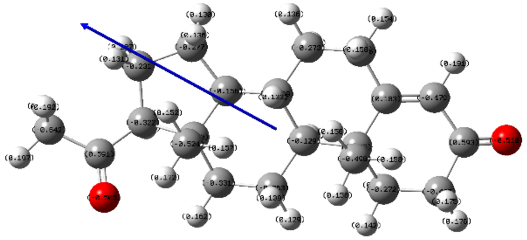 |
| **Fosinopril** | **Furosemide** | |
| 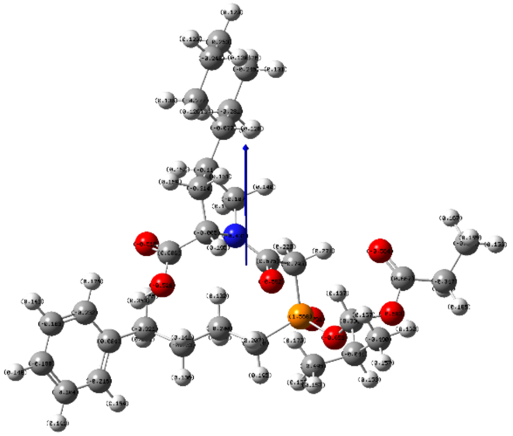 | 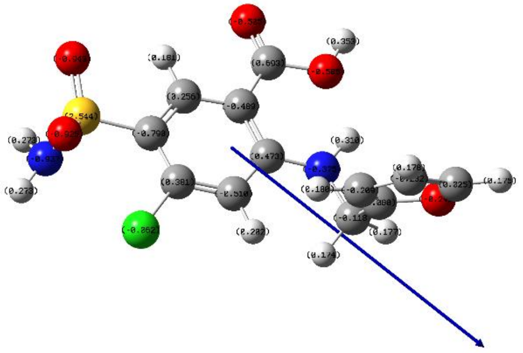 | |

### 6.3. Molecular electrostatic potential surfaces for all ligands (*i.e.*, drugs) in this manuscript

| **Meprobamate** | **Ciclopirox** | **Clotrimazole** |
| --- | --- | --- |
| 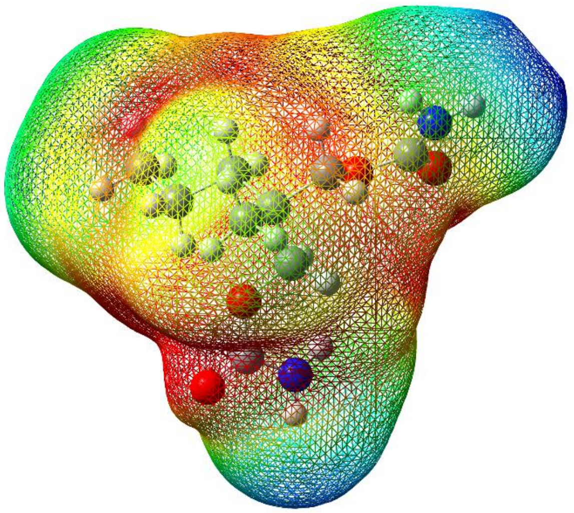 | 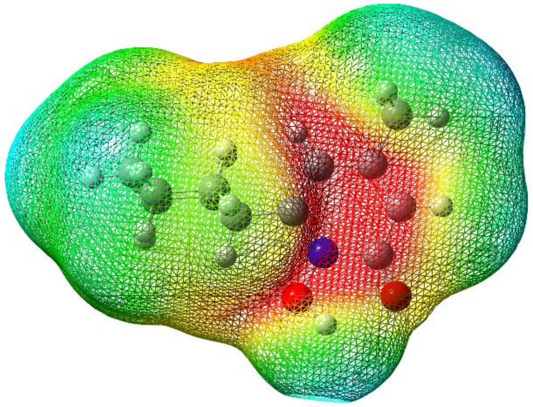 | 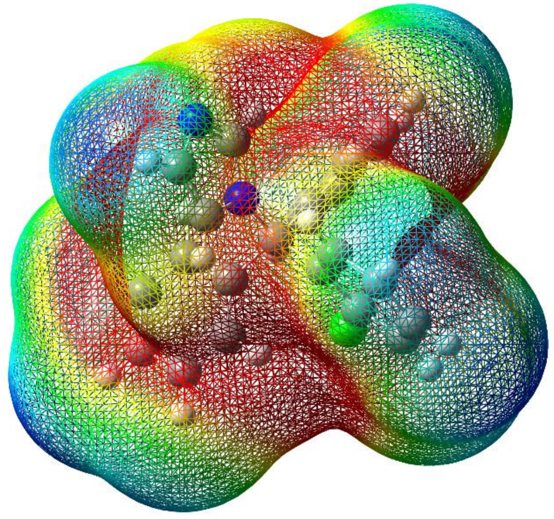 |
| **Griseofulvine** | **Naftifine** | **Natamycin** |
| 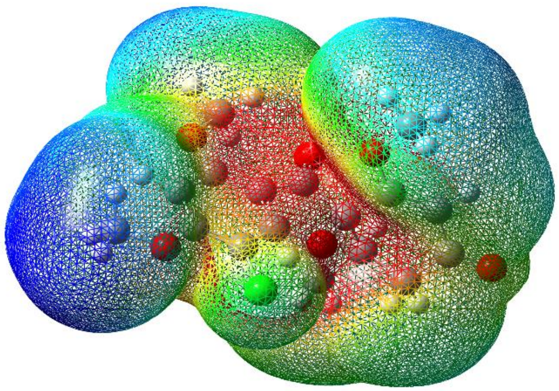 | 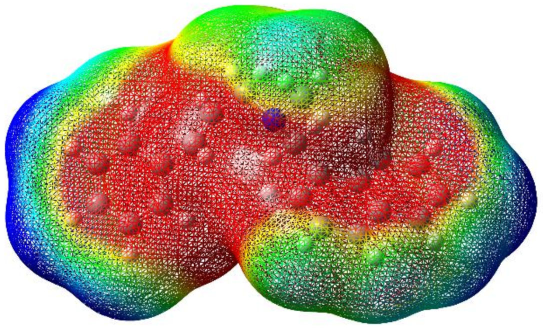 | 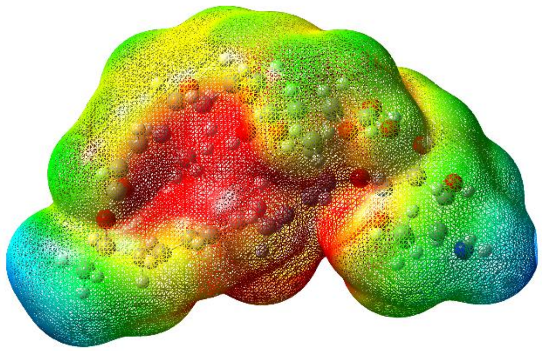 |

**Nystatin**

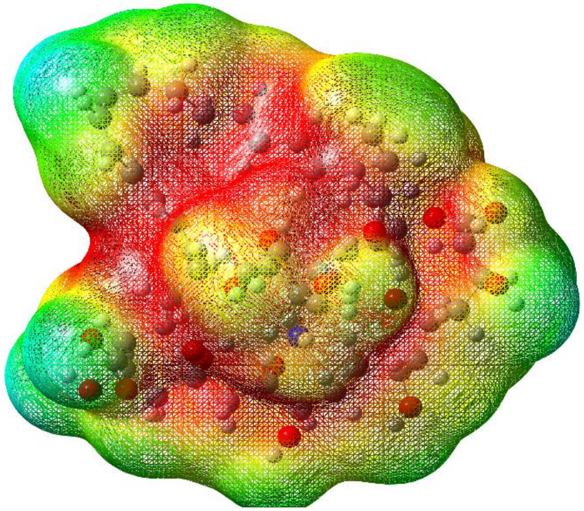

**Oxiconazole**

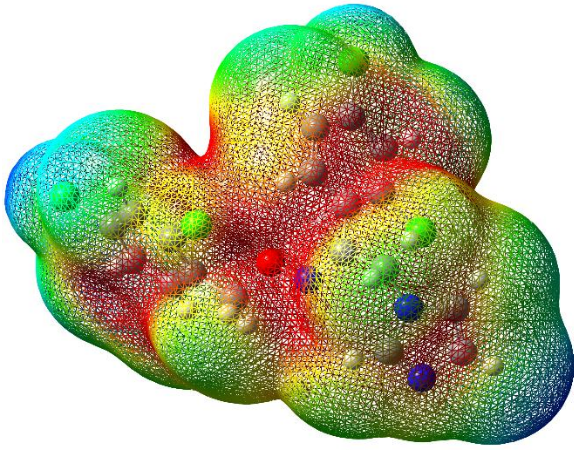

**Tolnaftate**

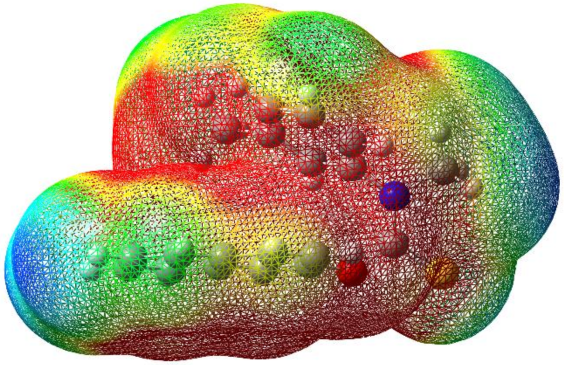

**Azelaic acid**

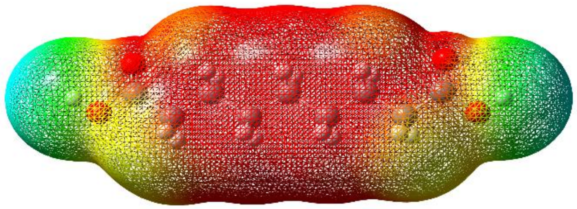

**Abiraterone**

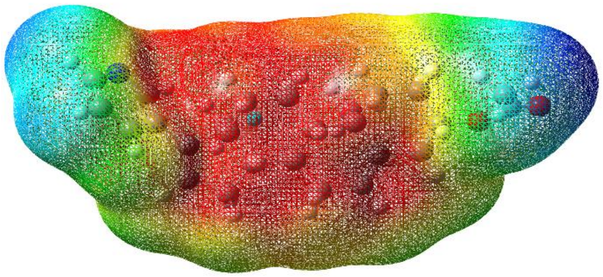

**Progesterone**

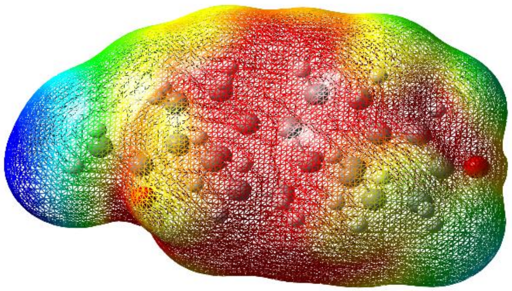

| **Fosinopril** | **Furosemide** |
|---|---|
| 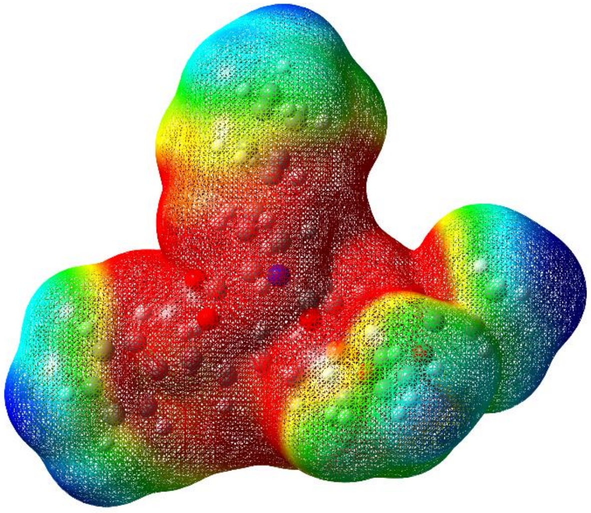 | 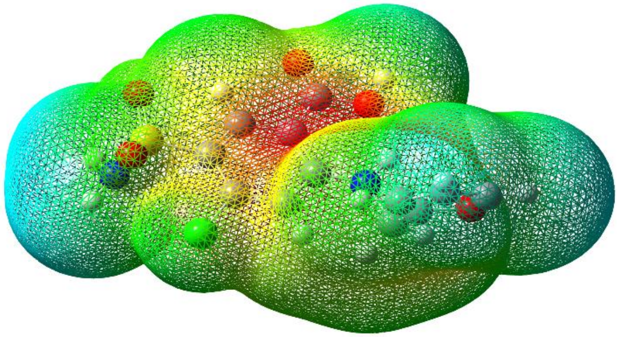 |
